# Spatial transcriptomic brain imaging reveals the effects of immunomodulation therapy upon specific regional brain cells in mouse dementia model

**DOI:** 10.1101/2023.01.20.524845

**Authors:** Eun Ji Lee, Sungwoo Bae, Minseok Suh, Hongyoon Choi, Yoori Choi, Do Won Hwang, Dong Soo Lee

## Abstract

Increasing evidence of brain-immune crosstalk raises expectations for the efficacy of novel immunotherapies in Alzheimer’s disease (AD), but the lack of methods to understand brain tissues make it difficult to examine therapeutics. Here, we investigated the changes of spatial transcriptomic signatures and brain cell type using the 10x Genomics Visium platform in immune modulated AD models by various treatments. To proceed with an analysis suitable for a single spot-based transcriptomics, we first organized a workflow for segmentation of neuroanatomical regions, establishment of appropriate gene combinations, and comprehensive review of altered brain cell signatures. Ultimately, we investigated spatial transcriptomic changes following administration of immunomodulators, NK cell supplements and anti-CD4 antibody, that ameliorate behavior impairment, and designated brain cells and regions showing probable associations with behavior changes. We provided the customized analytic pipeline into an application named STquantool. Thus, we anticipate that our approach can help researchers to interpret real action of drug candidate by simultaneously investigating the dynamics of all transcripts for development of novel AD therapeutics.

## Introduction

Central nervous system (CNS) and central immune system (bone marrow: BM) interaction, more specifically, brain-immune cross-talk can happen by a pathway of skull BM, meninges and their lymphatics, and cerebrospinal fluid (CSF) to brain parenchyma^1–14^ and/or by another pathway of choroidal plexus (CP) capillary-stroma-epithelium, CSF to brain parenchyma^15–19^ in addition to by the classic pathway of crossing blood-brain barrier (BBB)^20–23^. In an explicit neuroinflammatory diseases such as multiple sclerosis in humans or experimental autoimmune encephalomyelitis (EAE), immunoglobulins or immune cells have been considered to enter the brain parenchyma via BBB^20^ of brain parenchyma or via brain-CSF-barrier of CP^24,25^, or recently via arachnoid barrier cell (ABC) layer of skull BM-meningeal lymphatics and CSF/perivascular spaces reaching brain parenchyma^3–5,17,26–28^.

Novel immunomodulatory therapy in Alzheimer’s disease (AD) transgenic model such as 5xFAD mouse, if to be noteworthy, should be accompanied by the improvement of cognitive decline associated with aging and/or the amelioration of the transgenes’ adverse effects such as priming brain cells or immune responses during development and aging. When we inadvertently found the effect of anti-CD4 antibody during investigating the effect of aducanumab^29^ and also encountered the probable effect of allogeneic natural killer (NK) cell supplements in AD models^30^, we questioned which of cells or transcriptomic markers along the areas would be the best to predict the outcome of these currently unaccounted new therapy candidates. In AD mouse models including 5xFAD, the surrogate effect-markers of previous findings/trials of systemic or intraventricular administration of CD8+ T cells^31^, anti-CD8^32^ or anti-CD3^33^ antibodies, Treg cells^34–36^ (or for stroke model^37^ or DEREG model for traumatic brain injury model^38^), and amyloid-sensitized Th1 cells^39–41^ were the amyloid plaques/Aβ on immunohistochemistry and transcriptional signatures of major brain cells and brain parenchyma^32,33,35^ or CP infiltrating cells^25,42^. As systemically injected cells and immunoglobulins were not examined for their whereabouts or biodistribution, direct CNS effect or systemic actions on immune systems were always the alternative to explain the probable novel immunomodulatory therapy, which inevitably led to the insufficient understanding of the target cells and areas and thus inconsistent results among the reporting investigators.

Cellular transcriptome analysis from the buoyant single cell/nucleus RNA sequencing (scRNAseq/snRNAseq) allowed easy and fast clustering of cell types^43–46^. Recently available single spot spatial transcriptome (ST) analysis using solid phase on slides such as Visium®^47–50^, HDST^51^, slideSeqV2^52^, Seq-Scope^53^, or stereo-Seq^54–56^ were found to yield quicker and robust regional segmentation as well as reliable cell clustering. Regionally and cell-type specific characterization of one or more sections of the mouse brain based on this reliable anatomical segmentation allowed comparison of mostly the basal states between groups or even in others the task-related active states by calcium two-photon imaging and scRNAseq of visual cortex^57,58^. We can now investigate the therapy effect on the brain cells (and infiltrating or rare cell of the brain) of segmented brain areas using spatial transcriptomic brain imaging to look into the questions whether a probable immunomodulatory therapy yield their effect on major brain cells of each region of the brain after systemic administration of those drugs^30^. Biodistribution study after systemic injection can inform whether the immune cells or antibodies enter and directly interact with brain cells, however, if we don’t see the immediate presence of cells (usually none) and antibodies (less than 1% of injected dose) we used to assume they would influence the brain cells and pathologic processes. Transcriptomic changes owing to proper novel immunomodulatory therapy will enable us to zoom in the probable target cells/genes which would have caused or at least be associated with the expected behavioral effects of these therapy^30^. This is especially helpful at the beginning of pursuit of barely-probable new drugs for the investigators to be confident that they are in the right direction to modify and optimize the new therapy-candidates. Transcriptomic changes of a region or regions and a cell type or cells in a group are expected to explain the behavioral results of the mouse model. Or least not the last, the transcriptomic changes might predate the behavioral improvements. In both cases, we expect that transcriptomic analysis results would excel, in sensitivity and target-cell specificity^47,54^, over histological results of immunohistochemistry. Also, single-spot ST yield tissue-globally searchable data which can later be reanalyzed repeatedly when the marker gene combination^46,50, 59–62^ comes to be available.

To do this, we needed to advance the single-spot RNA sequencing and their analyses using the customized method to derive cell type/state-specific distribution of the Visium sections. Paying attention to the proper dissociation of cell types/states using the optimum/minimum number of genes and also to cell-cell interaction (CCI) and cell-cell communication (CCC)^63^, marker gene combination should be established with existing public database of scRNAseq/snRNAseq^46,49,59, 64–68^ and data of one’s own^30^. For both tasks, ready-to-adopt methods are available by Creative Commons regulations by the previous investigations.

Stromal and parenchymal cells of various organs including brain are now known to show common and specific characteristics of cell identity and their ontological characteristics, among which the best known are microglia and perivascular macrophages or resident macrophages^67,68^. Immune cells of monocyte-derived and resident macrophages have distinctive transcriptomic signatures, respectively, which predict their immune roles and tissue integrity-preserving roles per characteristic signatures^69^. It was also the case with T cells, where resident memory T_RM_ gut for intestines, effector memory T_EM/EMRA_ for blood, liver, and BM, mixed T_RM/EM_ for various organs and BM yield their own characteristic transcriptomic signatures which determined their differentiation of T cells in every tissue of interest dictating their respective functional roles^70,71^. Unfortunately, both of these recent cross-tissue, resident immune cell studies^69,71^ did not include brain, which mandates one’s own analysis.

In this investigation, we performed segmentation of brain regions on coronal/sagittal sections per dozens of animals using readily available methods and characterized the common pathologic transcriptomic signatures of 7-month-old 5xFAD mice. Immunomodulatory drugs were tried in these mice to confirm behavior improvement. ^99m^Tc-hexamethylpropyleneamineoxime (HMPAO)-labeled cell-tracking imaging^72^ ruled out the immediate infiltration of NK cells in the brain. Spatial transcriptomic changes of mice were examined after anti-CD4 immunoglobulin administration and expanded NK cell supplements treatment with the dose-schedule which proved to improve Y-maze alternation behavior impairment at this period of age in 5xFAD mice. Transcriptomic changes were dissected across areas and cell types/states using publicly-available methods and databases, and the analytical pipelines organized as an application named STquantool. We found that regional/areal gene-set-defined types/states specific cells showed characteristic differences after each trial-treatment in a genetic model of AD, 5xFAD mice on spatial transcriptomic Visium analysis. Combined brain major cells of neurons, astrocytes, microglia, and oligodendrocytes with their associated types and states and brain resident/infiltrating rare immune cells per each region were explored for their distinctive transcriptomic changes among mice groups to yield their probable association with behavior improvements.

## Results

### Spatial transcriptomic characterization of gene-set-defined type/state-specific brain major cells in 7-month-old wild type and 5xFAD mice

In total 63 mice of either wild type or 5xFAD, 35 coronal sections and 28 sagittal sections were put into the analysis. A spatial barcode was given for every spot, the unit tissue domain of spatial transcriptomics, and at least 50,000 reads were obtained from each of the 4,992 spots in a capture region. The brain tissues were covered by an average of 3,000 spots across all samples. Using the count matrices computed from Space Ranger as inputs to the reciprocal principal component analysis (RPCA) supplied by Seurat version 4.0 (Seurat 4.0 https://satijalab.org/seurat/), the multiple brain tissues were segmented based on their transcriptome patterns^73^. Optimizing the parameters such as the resolution of spatial clusters and others, we could yield the segmented spatial cluster images in every case including those with various treatments and manipulation. The treatments and manipulations are listed in **Supplementary Figure 1A** and **Supplementary Table 1**. This included anti-CD4 antibody treatment and NK cell supplement treatment groups. Others are 3-month-old 5xFAD, cervical lymphatic ligation, P301L model with or without amyloid/tau-rich lysate injection, and fingolimod hydrochloride (FTY720) injection with or without lipopolysaccharide (LPS) pre-treatment.

The difference between 10 ST data from wild type and 11 from 5xFAD animals was compared for each of 14 spatial clusters using 7 to 8 coronal and 3 sagittal sections, respectively. In the spatial clustering process based on RPCA, transcriptomes of wild type animals are used as pivots and the spots from diseased animals was recursively mapped to the PCA space of the wild type reference^74^. All the individual sections were visualized and verified for their authenticity of designating the areas according to already-known anatomical correlates (**Supplementary Figure 1B-D**). Only the askew sagittal section in a few mice missed dorsal striatum and instead supplied septal lobe in more median position, but the spatial clustering correctly showed the pair of dorsal/ventral striatum in some and septal/ventral striatum in others. The spots from each cluster and group were represented by a uniform manifold approximation and projection (UMAP) plot, a dimensionality reduction method for visualization, and the clusters were well separated in terms of gene expression (**Supplementary Figure 1E, F**).

Using the developed platform, STquantool, regional/areal transcriptomes representing major brain cell types/states were compared by averaging the major cell scores in wild type and 5xFAD mice (**Supplementary Figure 2**). The regional cell abundance measured by cell scores based on the curated marker genes was compared between groups and using Wilcoxon rank-sum test as a post hoc analysis. The regional cell scores were also calculated in 32 subtypes of neurons^75^, several types of astrocytes^76^, microglia, and oligodendrocytes^66^. Besides, the reactive state-specific changes and marker genes of astrocytes and microglia were compared between wild type and 5xFAD animals for 14 brain regions.

Brain major cells were classified to the types of neurons, astrocytes, microglia and oligodendrocytes and their allied cell types (**Supplementary Figure 3A**). Neurons were classified according to the reports of Hodge et al.^75^ of Allen Institute with Aeverman et al.’s^59^ a random forest hierarchical clustering method (NSForest) to define optimal marker gene combination for neuron subtypes which were verified by the reports of BICCN and another group’s approach^77–81^. Astrocytes were classified according to the types of white matter-associated and gray matter-associated astrocytes^76^ once and again region-specific astrocytes for cortex/hippocampus (telencephalon), thalamus, and others (diencephalon)^49,82^. Reactive astrocytes and their marker gene combinations were determined by the suggestions of Escartin et al.^83^ and other investigators (Habib et al.^84^, Ioannou et al.^85^, Chamling et al.^86^ etc.). Oligodendrocytes and their allied cell types followed the initial report by Marques et al.^66^ and also were verified by other investigators’ suggestions^85,86^. Microglia were classified according to their states but not types considering their homeostatic and reactive (microglia with neurodegeneration: MgND^87^, disease-associated microglia: DAM^88^, lipid-droplet associated microglia: LDAM^89^, etc.) states^68,90,91^ and thus no subtypes of microglia were assumed. Instead, the aging-related effect on its own or associated with amyloid pathology was examined to show amyloid pathology excluding the confounding effect of aging^92^.

Neurons were defined and named according to the recent reports of Allen institute^75,77–79^ and the report by BICCN^93^. Using the data by Xiemrakis et al.^64^ for defining cell types, the hierarchical clustering method suggested by Hodge et al.^75^ and NSForest (version 2.0) by Aeverman et al.^59^, neurons were typed, subtyped to 20 GABAergic neurons and 12 glutamatergic neurons. Their pattern of expression was displayed for wild type and 5xFAD mice in **Figure 1A**. The difference between wild type and 5xFAD mice for coronal and sagittal sections of 9 areas of interest was quantitatively analyzed (**Supplementary Figure 3B)**. Subtypes of GABAergic and glutamatergic neurons showed unique patterns along the wild type and 5xFAD mice. Expression patterns of the GABAergic somatostatin (Sst) subtypes in the amygdala differed between wild type and 5xFAD mice (**Figure 1B** **and Supplementary Figure 3C**). Increased expression in the amygdala was prominent in the 5xFAD compared to the wild type mice. Notably, the individual genes (*Sst, Nr2f2, Tac1*, and *Moxd1*) tended to express higher in the amygdala of 5xFAD mice among the gene combinations (**Supplementary Figure 4**).

**Figure 1.**
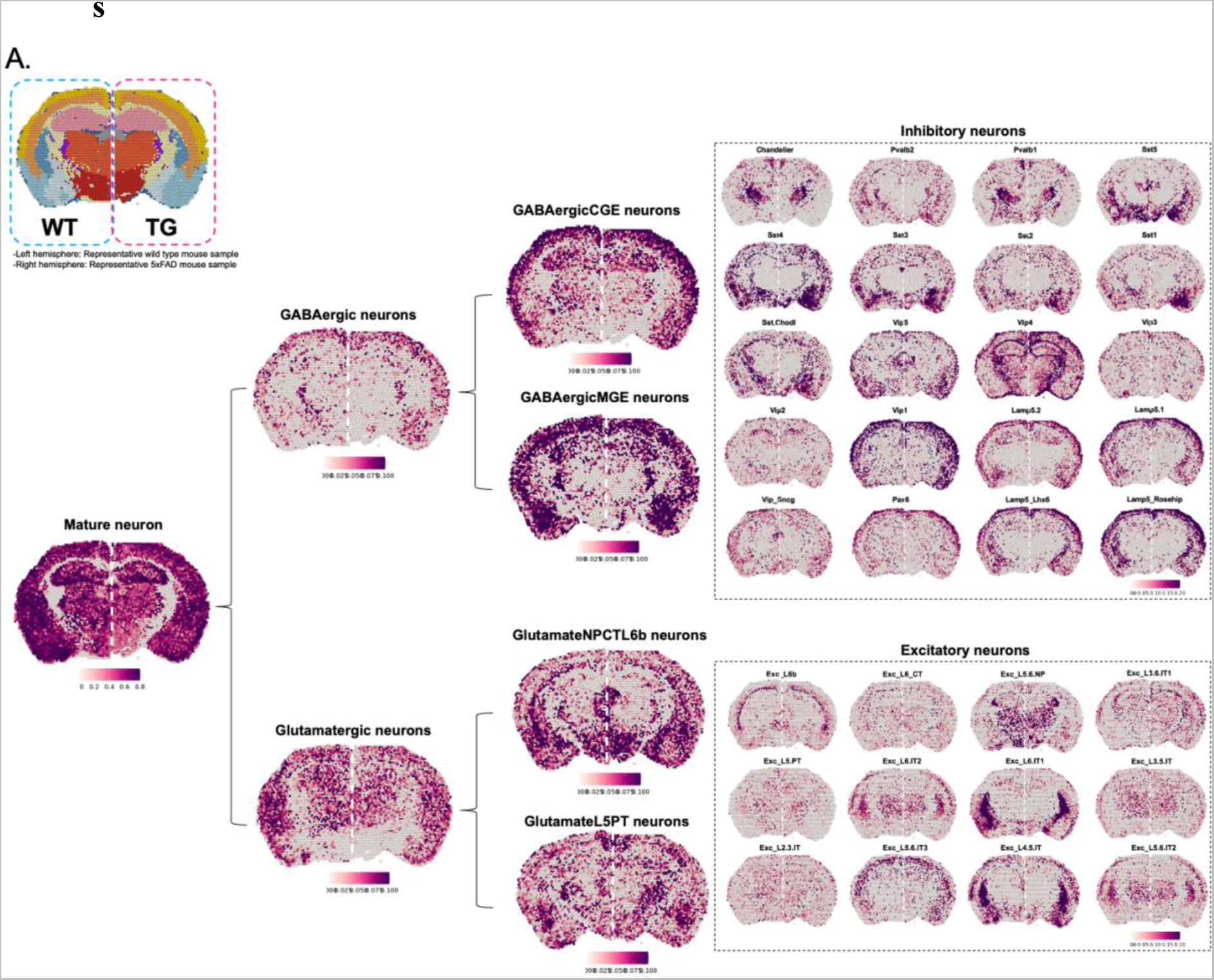
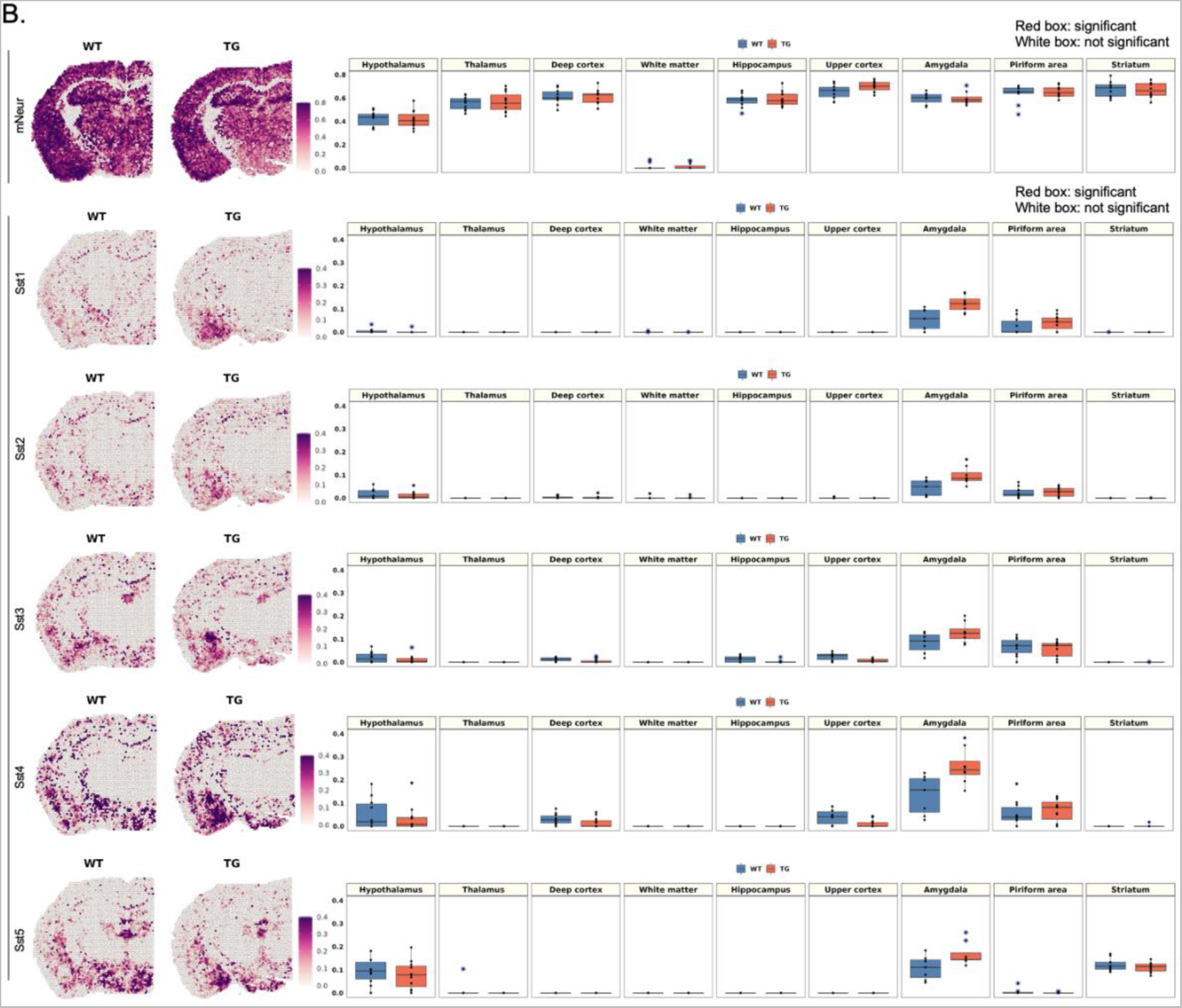
Brain region-specific expression patterns of the signatures of neuronal subclasses and spatial changes of neurons in 5xFAD mice compared to wild type mice. (A) Spatial patterns of diverse neuronal subclasses signatures. The left side of the hemisphere is representative slide of wild type mouse, and the right side is 5xFAD mouse. Each cell type showed a distinct region-specific expression. First, mature neurons were subdivided into GABAergic and glutamatergic neurons, and then the cells were further divided into subclasses to show the regional distribution of subclasses of inhibitory and excitatory neurons. (B) Spatial pattern of the neuronal signatures (mature neurons, Sst1, Sst2, Sst3, Sst4, and Sst5; left). Representative images of each group were selected among 10 spatial transcriptome data of wild type mice and 11 of 5xFAD mice. Spatial patterns of Sst-subclass of inhibitory neuronal signatures of the 5xFAD mice were the most remarkably different compared to those of wild type mice. The spatial distribution of Sst subclass neurons were similar between wild type and 5xFAD mice and the expression or Sst subclasses were higher in 5xFAD exclusively in the amygdala. The boxplot revealed the average module score of Sst-subclass inhibitory neurons and expression tended to be higher in the amygdala in 5xFAD mice, especially for the Sst4-subclass. Each dot represents a mouse in each group. (mNeur: mature neurons; Sst: somatostatin; WT: wild type; TG: 5xFAD mice; GABAergicCGE: caudal germinal eminence; GABAergicMGE: medial germinal eminence; GlutamateNPCTL6b: near projection, corticothalamic, and layer 6b; GlutamateL5PT: layer 5 and pyramidal tract)

The difference between wild type and 5xFAD mice was differential according to the definition (by gene combination to define reactivity) in reactive astrocytes and reactive microglia in their density and distribution (**Figure 2** **and Supplementary Figure 5A**). Reactive astrocytes and reactive microglia shared gene signatures and were supposed to collaborate to do the job of waste disposal in situ and out of the brain while promoting the interstitial fluid space (ISF) to perivascular/CSF space to meningeal lymphatics. Astrocytes were classified to cortical layers of deep and upper layer-specific or telencephalon- and diencephalon-origin according to Bayraktar et al.^82^ and to Kleshchevnikov et al.^49^. This classification did not disclose the difference between wild type and 5xFAD mouse (**Supplementary Figure 5A)**. However, another two types of white matter-associated and gray matter-associated astrocytes according to Werkman et al.^76^ yielded differential differences in density and distribution between wild type and 5xFAD mice. The white matter-associated astrocytes were significantly increased in the white matter and other gray matter regions in the 5xFAD mice, but no differences were observed in the gray matter-associated astrocyte signatures (**Figure 2A**). In addition, reactive astrocytes defined various ways^83–86,94^ which showed increase in density in white matter and neighboring gray matter areas (cortex and thalamus) in 5xFAD mice. Their distribution of reactive states was characterized to be diffuse but were prominent around the white matter on coronal and sagittal sections in 5xFAD at 7 months compared with wild type. Aging astrocytes showed significance but small differences between wild type and 5xFAD mice in the white matter, deep cortex, thalamus, and striatum (**Figure 2A**). Further analysis with individual transcriptomes used as markers for each state-specific astrocytes revealed the following findings. The expression of individual transcriptomes defining white matter-associated and reactive astrocytes showed similar patterns between wild type and 5xFAD, but the dominant individual transcriptomes were different (**Supplementary Figure 6A, B**). In the gene combination of white matter-associated astrocytes, *Lyz2, C1qa, Ctss, C1qb,* and *C1qc* were the top five genes with significant differences. In reactive astrocytes, *Gfap, Serpina3n, Vim*, and *C1qb* showed dramatic increases in 5xFAD compared to wild type mice.

**Figure 2.**
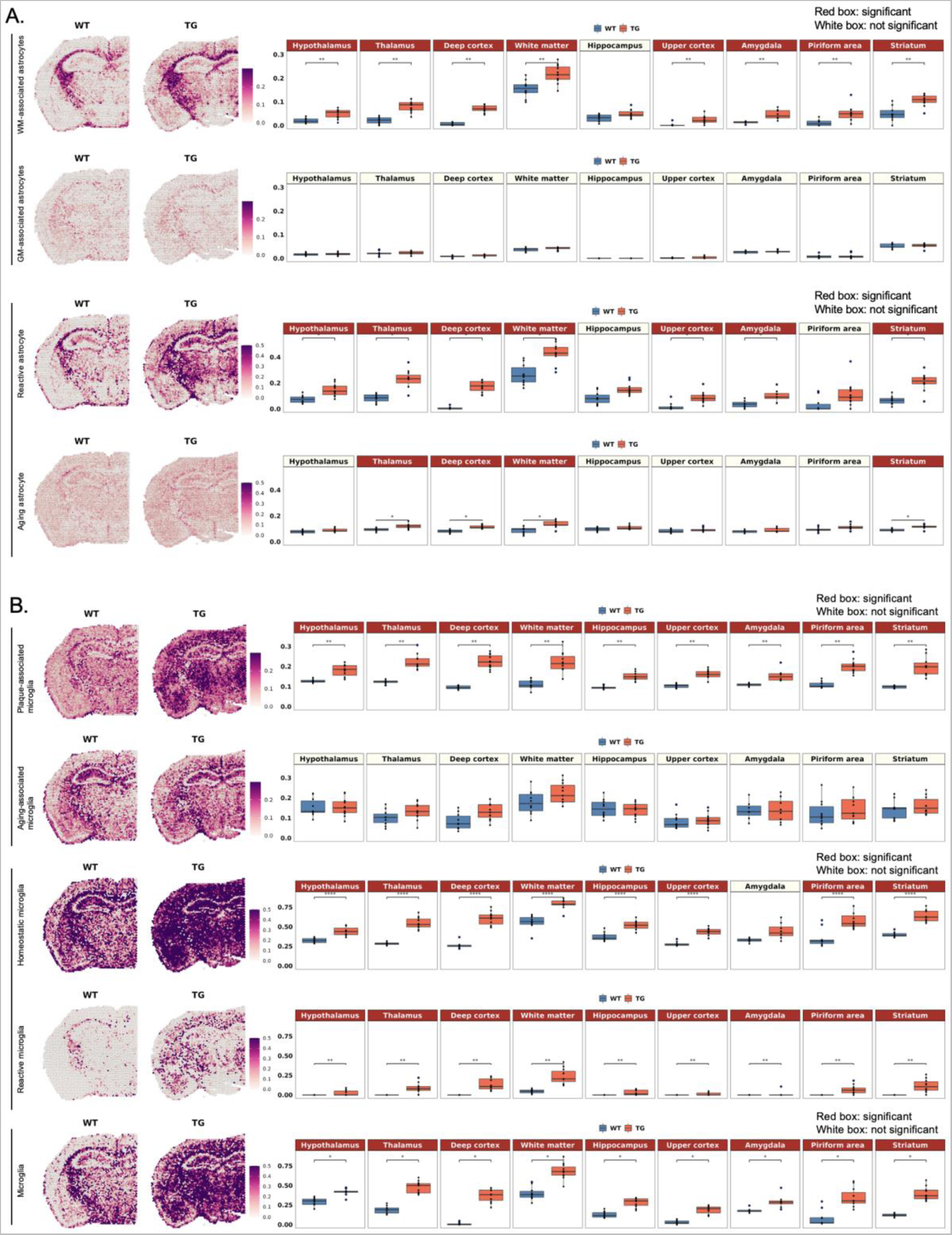
Spatial changes in the distribution of the region- or state-specific signatures of microglia and astrocytes in 5xFAD compared to wild type mice. (A) Spatial pattern of the region-specific signatures (white matter-associated and gray matter-associated astrocytes) and the state-specific signatures (reactive astrocyte and aging astrocyte; left). Representative images of each group were selected among 10 spatial transcriptome data from wild type mice and 11 from 5xFAD mice. Boxplot showing average module scores (right). Each dot represents a mouse in each group. The average module score of white matter-associated astrocytes was significantly increased in the white matter and other gray matter regions in the 5xFAD compared to wild type mice, but no differences were observed in gray matter-associated astrocyte signatures. Moreover, the average module score of reactive astrocytes showed a similar expression pattern to that of white matter-associated astrocytes, while significant but smaller differences were observed in white matter and several areas in the aging astrocyte signatures. (B) Spatial pattern of the state-specific signatures (plaque-associated, aging-associated, homeostatic, reactive, and pan-microglia). The average module score of plaque-associated microglia showed a significant increase in the 5xFAD compared to wild type mice, whereas aging-associated microglia showed no difference. Interestingly, both homeostatic and reactive microglia signatures showed a dramatic increase in the average module score in 5xFAD mice. Microglia in general (representing all the states-specific and non-specific signatures) showed increased expression in all the regions without showing any regionally distinctiveness. Bonferroni-adj. *p-value < 0.05, **p-value < 0.01, ****p-value < 0.0001. (WM: white matter; GM: gray matter; WT: wild type; TG: 5xFAD mice)

Microglia, classified to states of homeostatic ones and reactive one^88^ and aging-related and plaque-related ones^92^, showed increased density in wide areas for homeostatic state microglia and reactive microglia (disease-associated microglia by Keren-Shaul et al.^88^) and plaque-related (and aging-non-related but plaque-related) reactive microglia. Interestingly, both homeostatic and reactive microglia showed a dramatic increase throughout the regions in 5xFAD compared to wild type mice (**Figure 2B**). The plaque-associated microglia also showed a significant increase in 5xFAD mice, but aging-associated microglia showed no difference (**Figure 2B**). Of note, microglia signatures did not show differences by brain regions. Plaque-associated and reactive microglia shared a similar set of genes (**Supplementary Figure 6C, D**). Particularly, *Cst7, Spp1, Ccl6*, and *Axl* showed remarkable increases in 5xFAD compared to wild type mice in both microglial signatures. However, in homeostatic microglia signatures, other genes, such as *Hexb, Cst3, Cx3cr1, Tmem119*, and *P2ry12,* showed dramatic increases in 5xFAD mice.

Oligodendrocytes and their lineage cells classified by Marques et al.^66^ which comprise mature oligodendrocytes, myelin forming oligodendrocytes, newly formed oligodendrocytes, committed oligodendrocyte precursors (COP) and oligodendrocyte precursor cells (OPC), showed distinct distribution along the areas, mainly identified in the white matter and faintly in the thalamus and lateral hypothalamus (**Supplementary Figure 5B**). A significant difference of newly formed oligodendrocytes in the deep cortex and thalamus was observed between wild type and 5xFAD mice, but the expression was so low and the difference was also small. The classification according to Chamling et al.^86^, consisting of oligodendrocytes, OPC and cycling progenitor also showed similar characteristic distribution. The oligodendrocyte signatures showed relatively little differences between wild type and 5xFAD mice.

Astrocytes and microglia, specifically white matter-associated astrocytes, reactive astrocytes, plaque-associated microglia, homeostatic, and reactive microglia, tended to increase exclusively in the white matter in 3-month-old 5xFAD compared to the age-matched wild type mice. This meant that the changes with the signatures started at an earlier age and was around white matter, reflecting the similar result in our previous report^95^ (**Supplementary Figure 7**).

Lastly, differentially expressed genes (DEGs) were explored between the groups using the Model-based Analysis of Single-cell Transcriptomics (MAST) model^96^ to find the regional difference at the gene level. Of note, we sorted the extracted genes according to the abundance of genes and log-fold change for the single spot-based transcriptomic analysis. The spatial expression of individual DEGs in 5xFAD mice compared to wild type was visually assessed by STquantool (**Supplementary Figure 8 and Supplementary Table 2**). Venn diagrams of the significantly different transcripts per region were finally drawn and the exclusive genes were visualized as dot plots to examine the differences between wild type and 5xFAD. In 5xFAD mice, both the white matter and gray matter regions showed the significant increase of gliogenesis- and glial cell activation-related genes.

### Spatial transcriptomic characterization of rare brain-resident or infiltrating cells in 7-month-old wild type and 5xFAD mice

Spatial transcriptomic characterization of rare brain cells poses problems of finding the proper unique set of gene combinations for determining these rare cells residing among the confounding major cells. Unlike major cells whose distribution are already known, rare cells are rare in number and do not have any presumed distribution. The information of the propensity (rarity) of these cells are either derived from the scRNAseq study using the dissociated samples from various areas of the brain, even collected from a number of animals, or from the zoomed small areas observed on histochemistry. Abundance studies of rare immune cells in the brain reported that T cells were 4/mm^3^ while neurons were 90,000/mm^3^ and microglia 6,500/mm^3^ ^97–99^. Other cells such as B cells, monocytes, infiltrating macrophages, dendritic cells (DCs) either conventional or plasmacytoid, or neutrophils were counted and reported for the brain tissue as a whole because all these studies were from scRNAseq analysis using floated dissociated brain cells.

In contrast to the difficulty of obtaining information from previous studies disregarding the heterogeneity distribution of rare immune cells in the brain regions, the solid-phase spot RNA sequencing enabled the exact localization of the cells of interest, which was first documented in Ståhl et al.^47^. On this method and now available Visium ones, a spot has its own count (log1p of count ratio) not contaminated by any false positive derived from mixing samples and the detection limit is not affected by major cells of higher density. The only problem is to discern the rare cells from others with an appropriated gene (transcriptome) combination to exactly sort out only the specific marker transcriptomes. Selecting the possible key gene sets defining rare cells with highest specificity is influenced by the choice of the input data, which are composed of participating cells^50^. For example, T or B lymphoid cells, quite unique with their highest propensity of ribosomal protein genes such as *Rpl* or *Rps*, are characterized, by any cell-type annotation methods, to yield the candidate marker gene combinations. However, as other major brain cells are also equipped with these protein-producing genes expressed in sufficient amount to pretend to look like rare brain cells so as to confound the presence/density of rare lymphoid cells in any areas of the brain. Also, since the rare immune cells are commonly investigated by combining cell sorting strategies with scRNAseq, the rare cell markers acquired from the subpopulation single-cell dataset may overlap with the major cell type markers. This caused serious overestimation which was disclosed immediately on visual assessment. This was also the case despite the use of the recent data available by Schafflick et al.^68^ and NSForest by Aeverman et al.^59^. We adopted visual curation for excluding the frankly absurd transcriptomes as marker gene candidates, and finally sorted out the rare cells with optimal marker gene combinations thereof to compare wild type and 5xFAD mice (**Figure 3** **and Supplementary Figure 9**).

**Figure 3.**
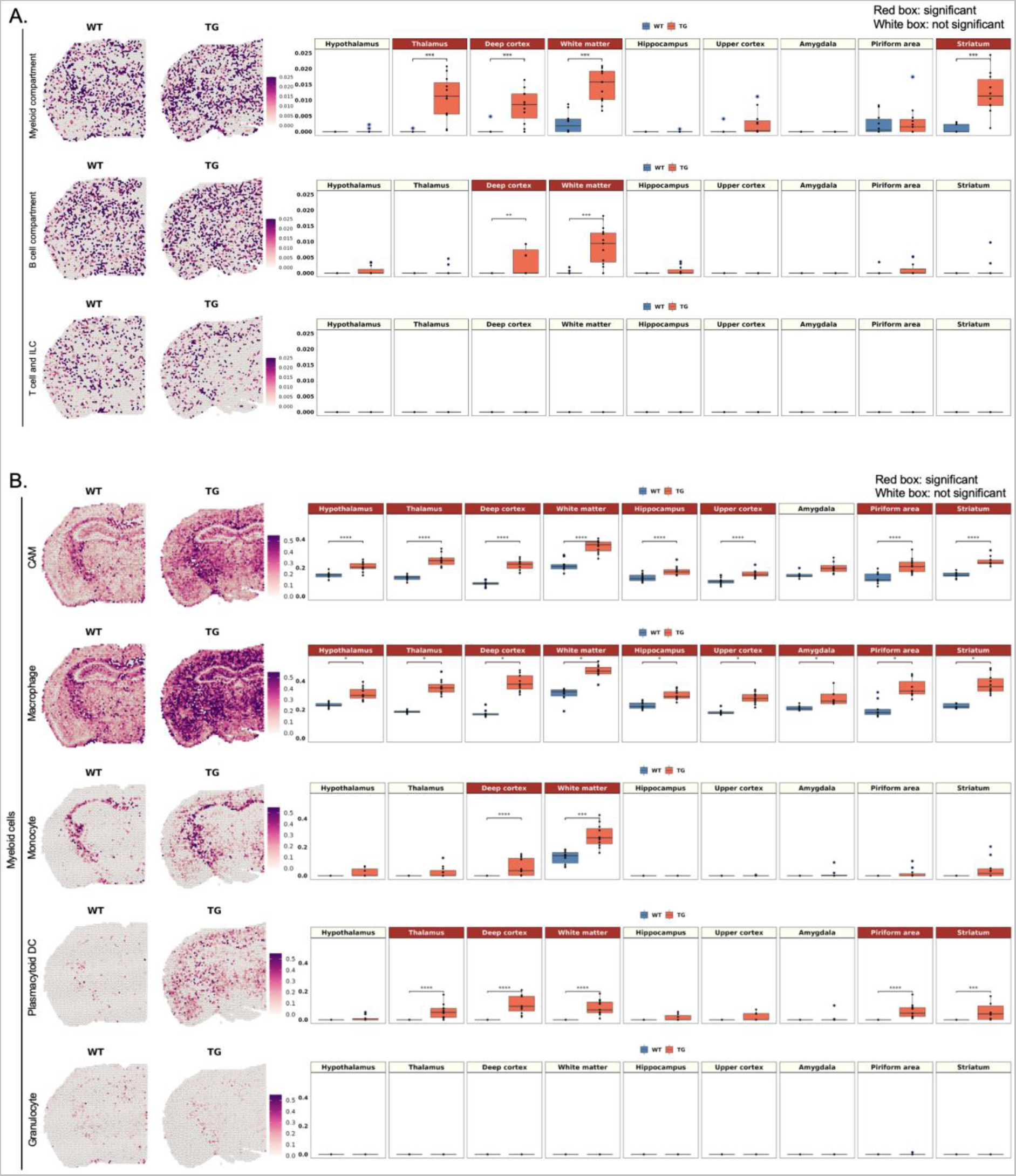
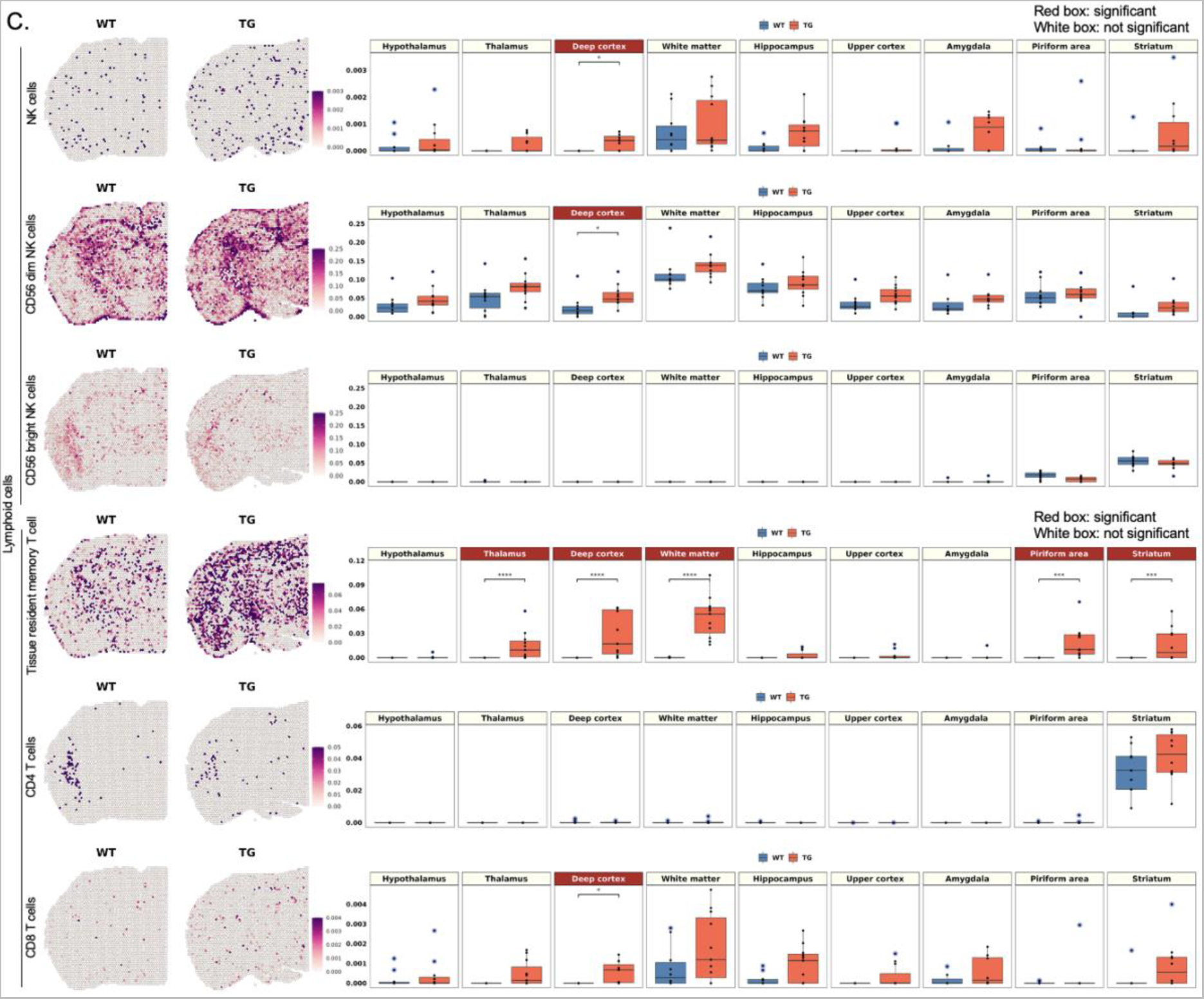
Spatial changes in the distribution of myeloid and lymphoid cell signatures in 5xFAD compared to wild type mice. (A) Spatial pattern of the signatures of myeloid, B cell, and T cell and ILC compartment according to the marker gene combination reported from Dominguez Conde et al.^71^ (left) and the boxplot showing the average module scores (right). Each dot represents a mouse in each group. The average module score of myeloid compartment showed a significant increase in the white matter and the gray matter regions adjacent to the white matter, including the thalamus, deep cortex, and striatum. B cell compartment signatures showed increase in deep cortex and white matter. In contrast, T cell/ILC compartment signatures were low without difference between wild type and 5xFAD. (B) Spatial pattern of the subtype signatures of myeloid cells, including CAM, macrophage, monocyte, plasmacytoid DC, and granulocyte according to marker gene combination from Schafflick et al.^68^. Notably, CAM and macrophage signatures showed the most pronounced increase in 5xFAD compared to wild type mice in most of the regions. Monocytes and plasmacytoid DC increased expression in deep cortex and white matter, and plasmacytoid DC further in thalamus, pyriform area and striatum. (C) Spatial pattern of the subtype signatures of lymphoid cells, including NK and T cells according to marker gene combination from Xiemrakis et al.^64^. In the case of NK cell signature, a significant increase was observed in the deep cortex of 5xFAD mice, which is associated with increase of CD56 dim NK cells. Among T cell signatures, tissue resident memory T cell signatures was higher in 5xFAD mice in deep cortex, white matter, thalamus, pyriform area and striatum. CD4 signature was explicit in the striatum both in wild type and 5xFAD mice but the expression was too low to show quantitative difference between wild type and 5xFAD. Bonferroni-adj. *p-value < 0.05, ***p-value < 0.001, ****p-value < 0.0001. (WT: wild type; TG: 5xFAD mice; ILC: innate lymphoid cells; CAM: CNS-associated macrophage; DC: dendritic cells; NK: natural killer)

As immune resident cells were classified in three ways 1) using novel data by Eraslan et al.^69^ (**Supplementary Figure 9A**) for tissue-specific monocyte-derived macrophages and data by Dominguez Conde et al.^71^ (**Figure 3A**) for tissue-resident T cells, 2) using the data by Schafflick et al.^68^ (**Figure 3B**) and markers refined using NSForest 2.0 and 3) using the data by Xiemrakis et al.^64^ and refined using NSForest 2.0 by Aeverman et al.^59^ (**Figure 3C**). The first two data^69,71^ were derived by using various tissues except brain while the other two reports^64,68^ were by using brain tissues.

Defining maker gene combination was more intricate for these rare immune resident/infiltrating cells as they are defined by surface markers in the report of Eraslan et al.^69^, in other organs/tissues than brain, or by transcriptome signatures suited for each study. Though the data were from the body tissues, not brain, in the first approaches, as the tissue stromal cells are included in DEG analysis and assuming stromal cells might be more similar between tissues including brain, transcriptomes of major parenchymal/stromal cells co-expressing with rare immune cells were to be correctly excluded. We chose NSForest to help exclusion of confounding stromal tissues. Monocyte-derived macrophages, among two types of which by Eraslan et al.^69^, one for immune function (MHCII+) and the other (LYVE1+) for using vascular integrity and repulsion of infiltrating immune cells. For the immune function, specifically for brain, conventional DCs with MHCII+ cells were found to be effective for antigen presentation adaptive immune cells (T cells and B cells) though border-associated macrophages or microglia were not^100^. We asked whether this surface marker-defined characterization can be translated to mouse amyloidosis (5xFAD) using the signature of MHCII+ related immune-functioning macrophages and LYVE1+ related integrity-charged macrophages by Eraslan et al.^69^ (**Supplementary Figure 9A**). Integrity-charged macrophages were not different among the groups of 7-month-old wild type and 5xFAD mice. Spot signatures of immune macrophages were more abundant in 5xFAD mice compared with wild type mice in the white matter.

Signature gene combinations used for cross-tissue immune cell analysis by Dominguez Conde et al.^71^ for T cells and innate lymphoid cells (T/ILC), B cell compartment and myeloid compartments revealed no difference between wild type and 5xFAD mice for T/ILC but significant difference among these mice for myeloid compartment cells in the white matter and the gray matter regions adjacent to the white matter, including the thalamus, deep cortex, and striatum and for B cells in the deep cortex and white matter (**Figure 3A**).

This result came from the following stepped analysis including curation procedure. At first step of curation, individual transcriptomes belonging to the three compartments by Dominguez Conde et al.^71^ were examined visually for their distribution/intensity and several transcriptomes which were already reported in the literatures as signatures for major brain cells and their reactive states, were removed, which excluded the background effects of abundant brain cells, eventually to yield marker gene combinations for three compartments and their cell types. Having removed 1) *Cx3cr1* and *Tyrobp* (microglia) from T/ILC, 2) *Ighm* (Scheurer et al.^101^ for neurons), *C1qa* (microglia) from B cell compartment, 3) *Trem2* (microglia), *Clu* (astrocyte), *Selenop* (microglia, astrocytes, oligodendrocytes), *Igf1* (reactive microglia and reactive astrocytes), as well as *C1qa* and *Cx3cr1* from myeloid compartment, scores of T/ILC still did not disclose difference between wild-type and 5xFAD mice, and scores of B cell or myeloid compartments revealed yet the slight but significant increase in 5xFAD compared with wild type mice. Individual variations within 5xFAD mice could be recognized on visual assessment too. For individual genes for T/ILC, localization was prominent for *Cd4* (striatum) and little difference regardless of abundance (*Slc4a4, Spry2, Ncam1, Pcdh*9 are abundant), no difference was observed except for *Pdcd1* (smaller cell fraction in various T cells including Trm/em_CD8 according to Dominguez Conde et al.^71^), which was slightly increased in 5xFAD. For B cell compartment, the difference between mouse groups, if any, were presumed to be due to *Itgax* and *Fcrls*, both of which were related with aging-associated B cells and *Fcrls* further with memory B cells and plasma cells/plasma blasts. For myeloid compartment, the difference of scores among mouse groups was contributed by *Tyrobp, Lyz2, Fcer1g, C1qc,* and *Apoe*, all of which are related with various types of tissue-specific macrophages and classical/nonclassical monocytes (**Supplementary Figure 10**).

The second one by Schafflick et al.’s data^68^ was tested for either the marker gene combination recommended by Schafflick et al. according to their supplementary table (LFCR>0.5) for 12 border cell leukocytes (including microglia) or the marker gene combination curated by NSForest upon their data. Schafflick’s own data yielded obviously too high intensity for CD4 and CD8 T cells among 12 border-associated leukocytes, such as B1, B2, CD4 T, CD8 T and NK cells, microglia, CNS-associated macrophages (CAM), macrophages, monocytes, myeloid DC (mDC), plasmacytoid DC (pDC) and granulocytes. When we surveyed the constituents of tentative marker transcriptomes for these inappropriate signature, ribosomal genes (many isoforms of *Rpl* and *Rps*) were the faulty annotation of CD4 T and CD8 T cells. This mis-annotation is assumed to be caused originally from the fact that the parenchymal and border leukocytes were included after their selection with CD45 positivity, meaning that they could not exclude the differential expression of these cells from the major brain cells including stromal cells. Individual transcriptome per spots were easily observed to disclose that if we chose highest LFCR with adjusted p values for determining marker genes, *Ighm* for b1 cell (also found in cortex not related with B cells)^101^, *H3f3b* (histone protein also nonspecific for brain) for b1 cells, or *Stmn1* for b2 cells (rather brain-wide expression), *Dut* (enzyme for nucleotide and ubiquitous including brain cells) for B cells and many similar examples (**Supplementary Figure 9B**). Though DEG analysis depends upon the input data composition, we tried NSForest to Scahfflick’s data and obtained better marker gene combination. This Schafflick/NSForest yielded improved intensity matching considering the prevalence of cells population in the brain parenchyma except for b2 cells (still too dense due to *Tuba1b* (tubulin related)) and CD4 T cells (depending heavily upon one transcriptome *Trbc2* (T cell receptor beta constant 2 but expressed also in cortex)), respectively. The other 10 cells’ signature looked to represent the cells intensity/distribution correctly, however, nonspecific and dense *Apoe* for CAM, dense *Cst3* for macrophages, *Mal* (full name of Myelin And Lymphocyte Protein implies its localization both in lymphocyte and myelin of neurons) for monocytes, *Tyrobp* (in association with *Trem2*, well-known marker for microglia) for both monocytes and pDC. Upon the application of NSForest, 6 to 10 maker genes were obtained and zero to three genes should be adjusted (kept or removed meaning curated by operators’ consensus). Application of curated gene combination to our ST samples revealed that CAM and macrophages showed the most pronounced increase in 5xFAD compared to wild type mice globally throughout the regions (**Figure 3B**). Also, pDCs showed increases in the white matter and some gray matter regions. However, transcriptome density of DCs were considered inappropriate as it yielded much higher intensity along the entire brain, considering that DCs occupy only 0.14% of myeloid/lymphoid cells of the brain and border including microglia (0.8% among myeloid/lymphoid cells excluding microglia)^68^.

Among lymphoid cell signatures, an increase in the tissue resident memory T cell was inferred in 5xFAD compared to wild type mice (**Figure 3C** **and Supplementary Figure 10**). However, it is necessary to consider technical limitation of spot-based transcriptomic analysis for evaluating rare brain cell signatures. It is still unclear which genes specifically define rare cells, while gene combinations may overlap with major brain cells on ST brain imaging.

Using single gene as a marker will be better and convenient to designate a rare cell type. This was possible to designate infiltrating macrophage derived from circulating monocytes originating from BM (**Supplementary Figure 11**). CD11c surface marker and its gene *Itgax* was the marker for this cell. Resident T cells was suggested to be CD73 positive and its gene *Nt5e* by Fang et al^102^. CD56^bright^ and its gene *Ncam1* was considered to be circulating and immature NK cells marker but express high also in neurons^103,104^. Perivascular macrophage caused a great problem to be distinguished from microglia, finally *Lyve1* was the discriminator of pvMϕ and microglia (*Sall1*)^105^. Similarly, for brain major cells, *Trem2* and *Tyrobp* was suggested to be conjoint marker for microglia, *Cspg* and *Olig2* were supposed to represent OPC, not any other cell types. Homeostatic microglia could have been defined by *Sall1*, however, transgenic mouse study^105^ found not *Sall1* but *Hexb* was the better marker for authentic microglia. The importance of *Aif1* (IBA1) as activated microglia and of *Gfap* as activated astrocyte was disclosed to be non-specific or at least subtype-specific marker, respectively. Once a marker was defined well for designating rare cell type well discriminated from major brain cells including microglia and perivascular spaces (pv) macrophages (and submeningeal macrophages), then that marker in a spot will disclose the fact that the gene signature of that spot might be from the rare cell of interest or the fact that the signature is never from the rare cell if no signal were observed. Multiplication of spot scores for a gene and cell-designated marker gene combination could do this job. Genes widely expressing over all the cell types but with specific isoforms could also do the job successfully and *Prdx* (for peroxiredoxin) was one of the examples (*Prdx6* and *Prdx2* for astrocytes, *Prdx4* for microglia and *Prdx1* for oligodendrocytes) (**Supplementary Figure 12**).

### Improvement of behaviors with much variation by immunomodulatory therapy of anti-CD4 antibody and NK supplements in 5xFAD AD mouse model

With preliminary behavioral study to prove the effect of aducanumab, pretreatment of anti-CD4 antibody caused larger degree of changes in alternation scores in the control animals (meaning higher improvement in the group of animals of anti-CD4 antibody treatment^29,30^. Three batches of several animals with anti-CD4 antibody treatment reproduced the previous group-wise behavioral improvement with the similar much variation (67.7% ± 18.4%) at the 7-month of age of 5xFAD model mice (**Figure 4**). We assumed that anti-CD4 antibody treatment modulated systemic adaptive immune system as transgenic insertion of five types of mutated human APP/PS1 genes would have caused immune disturbance due to their presence in the mouse chromosome, meaning the presence of human mutated genes would have resulted in the brain-immune interaction dysfunction as well as in the plaque-prone amyloid burden in animals. Spatial transcriptomic analysis was considered to reveal the eventual response of brain cells, either major or rare resident and infiltrating immune cells, if any.

**Figure 4.**
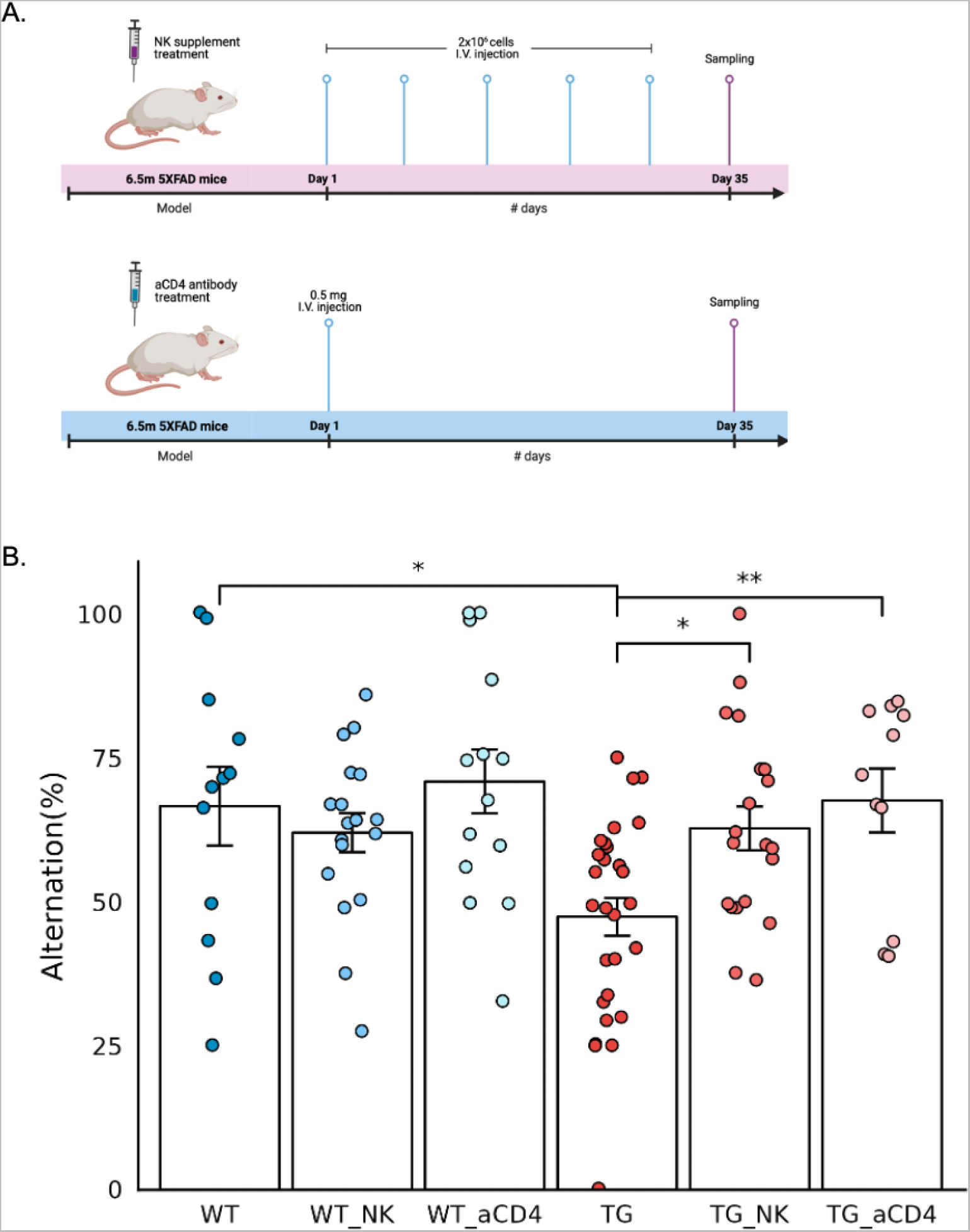
Improved behavior after intravenous administration of NK cell supplement and anti-CD4 antibody in 5xFAD mice. (A) Timeline of the experiments for intravenous NK cells supplements (upper) and anti-CD4 antibody (lower) administration in 6.5-month-old wild type and 5xFAD mice. NK cells (2×10^6^ cells/injection) were administrated once a week, a total of five times, and anti-CD4 antibody (0.5 mg/injection) was administrated once as a single injection. After a month, behavior analysis was performed, and brain tissue samples were obtained for spatial transcriptomic brain imaging analysis. (B) Behavior function of exploring new environments was examined using the Y-maze test and expressed as alternating percentages. Each dot represents a mouse in each group. The alternation rate was decreased in 5xFAD compared to the wild type mice with much variation at this age of mice, wild type and 5xFAD. And the alternation percentage score of 5xFAD mice increased significantly after injection of NK cells supplements and anti-CD4 antibody treatments than 5xFAD mice without treatments. Wild type mice also showed variation, however, their alternation score were not different between no treatment and either treatment groups. Wilcoxon *p-value < 0.05, **p-value < 0.01. (aCD4: anti-CD4 antibody; WT: wild type; TG: 5xFAD mice; WT_NK: NK cell-treated wild type; WT_aCD4: anti-CD4 antibody-treated wild type; TG_NK: NK cell-treated 5xFAD; TG_aCD4: anti-CD4 antibody-treated 5xFAD)

With another preliminary study with APP/PS1 model mice with water maze with expanded NK cell supplements derived from spleen of wild-type BALB/c mice, anecdotal cases of behavioral improvement were observed (data not shown). Three batches of allogeneic NK cell supplements, as 5xFAD is B6 lineage, reproduced the behavioral improvement of alternation scores on Y-maze tests on average, however, with much variation (**Figure 4**). Much variation on both anti-CD4 antibody treatment study and NK supplements study reminded that 7-months of age in 5xFAD mice would have been going through their own course of aggravation of the pathological changes of Aβ oligomer insults and amyloid plaque burden resulting in the later pathological and behavioral nadir dysfunction around 12-months of age or later. Spatial transcriptomic analysis was considered to reveal the regional and cell-type specific changes of transcriptomes of brain major and rare cells corresponding to each individual mouse’s degree of behavioral dysfunction in the NK supplements-treated group, too.

### Regional cell-type/state specific transcriptome changes of 5xFAD compared with wild type mice after intravenous administration of NK cell supplements

Three mice with higher alternation score on Y-maze test were selected for each saline-treated and NK cell supplements-treated group (**Supplementary Figure 13**). Coronal/sagittal brain sections of these mice were put into the Visium analysis. Each group were paired to the same plates so that the batch effect of the read per slide would be minimized. Using 30,000 to 50,000 counts per mouse, we retrieved the count data which were normalized for their total count and log1p of the ratio data were used for further analysis. Spatial clustering allowed anatomical segmentation to yield 14 regions with almost similar size (**Supplementary Table 3**). Cell type and state-specific marker gene combination was also used to analyze the cell-specific and/or cell state-specific changes after NK cell supplements treatment. For 5xFAD case with NK cell supplements, one mouse with very-high behavior score among the group in a batch was chosen to make three (batches) representatives ‘behaviorally best’ and another mouse with very-low score among the group in a batch was chosen to make three representatives ‘behaviorally worst already at 7 months of age’. Thus, this was merely the looking into the transcriptomic changes according to the behavioral impairment of the 5xFAD mice at the age of 7 months. NK cell supplements contributed at least the widening the distribution of scores of behavioral impairments at this middle age.

GABAergic Sst subtype neurons showed significant decrease after NK supplements in the amygdala, which showed abnormally increased signature in 5xFAD (n=11) compared with wild type mice (n=10) (**Figure 5A**). Among the Sst neuronal signatures, *Sst, Tac1*, and *Nr2f2* showed dramatic decreases in the amygdala after NK cell supplements administration in 5xFAD mice, but no distinct differences in the other regions (**Supplementary Figure 14A, B**). No significant difference was found either in the mature neuron score or in any other subtypes of neurons than Sst neurons between 5xFAD mice without NK supplements and those with NK supplements treatment. Also, the NK cell signature tended to increase after administration of NK cell supplements exclusively in the white matter region of 5xFAD mice (**Supplementary Figure 15A**). The change in the module score level was not observed by the anti-CD4 antibody treatment, which was in contrast with the change by the NK cell supplement treatment (**Supplementary Figure 15B**). The signatures of astrocytes, microglia, and oligodendrocytes did not show any difference. Also, no difference of the brain rare cells, either resident or infiltrating, was observed (**Supplementary Figure 16A-D**). The biodistribution of ^99m^Tc-HMPAO-labeled NK cells was examined using SPECT/CT to determine how systemically injected NK cells caused changes in the brain (**Supplementary Figure 17**). Within 1 h after injection, the labeled NK cells were mainly taken up by the liver, and this radioactivity decreased gradually by 16 h. Of note, no definite brain uptake of the labeled NK cells was observed with the resolution of SPECT/CT images. Thus, NK cells may have caused changes of brain cells at the transcriptional levels by indirectly via cytokine or other secretory factors released by NK cells and/or inherent peripheral immune cells influenced by supplemented NK cells.

**Figure 5.**
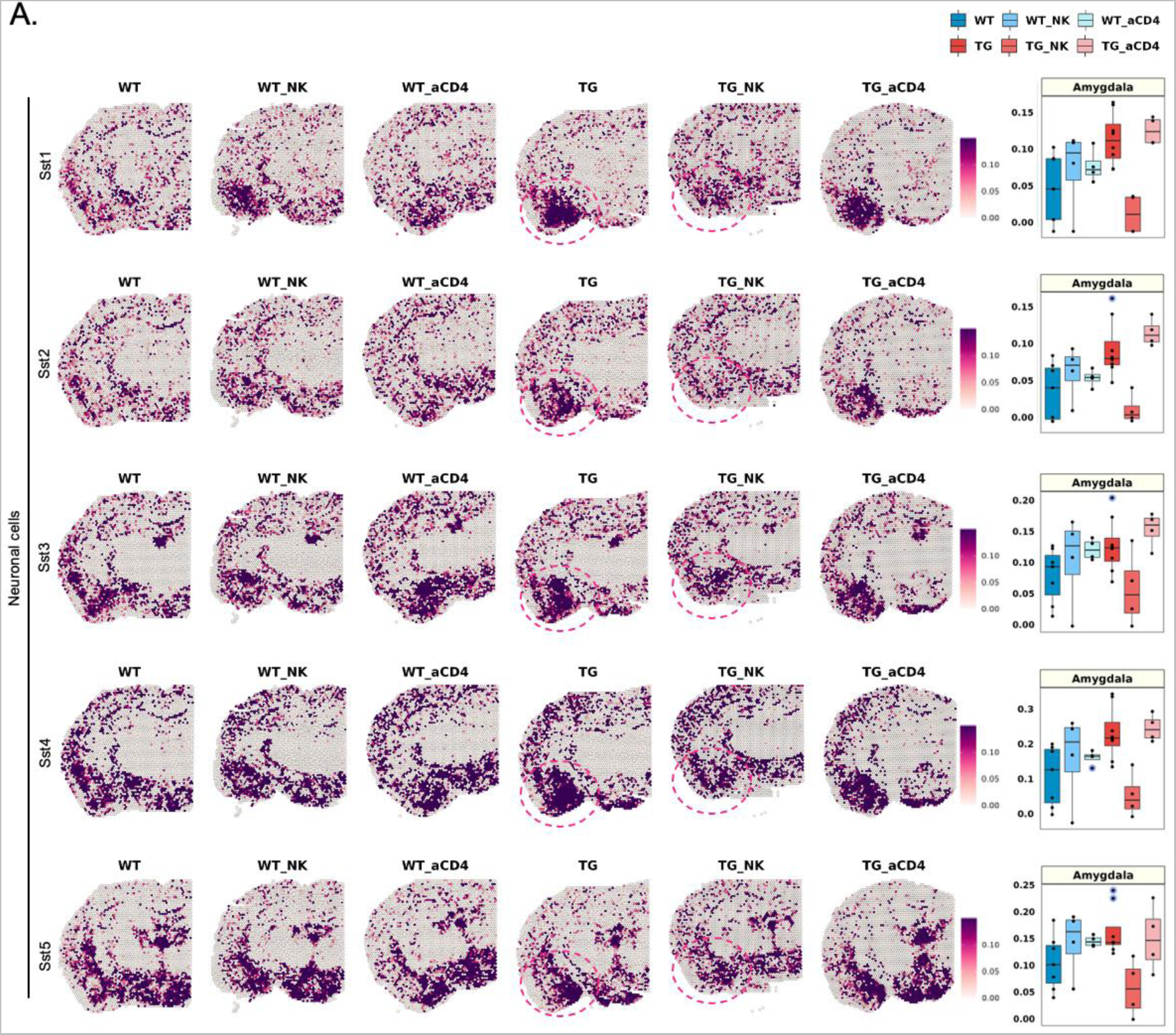
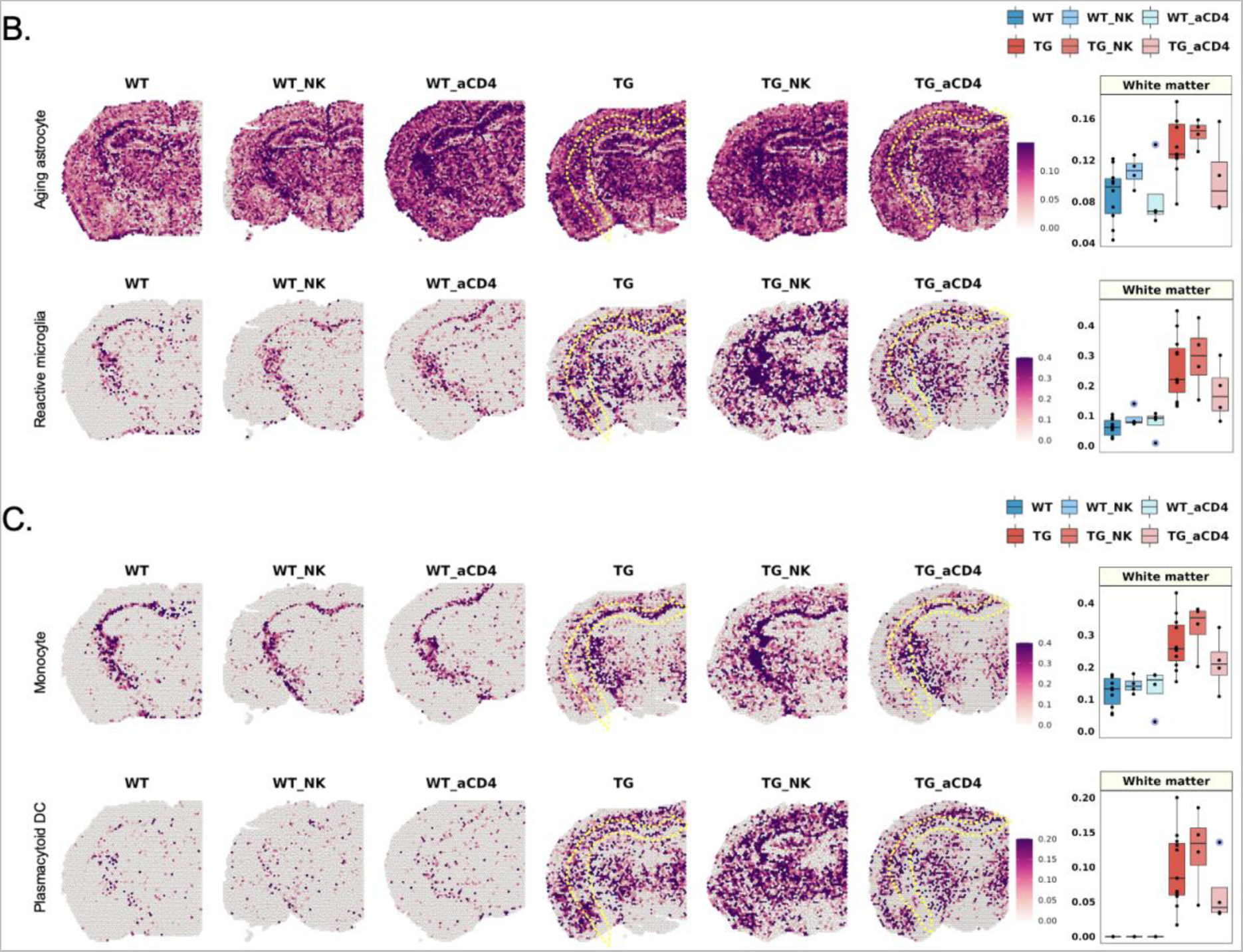
Brain region-specific transcriptome changes in cell signatures after NK cell supplement and anti-CD4 antibody treatment in 5xFAD mice. (A) Spatial pattern of the signatures of somatostatin (Sst)-inhibitory neuronal signatures (Sst1, Sst2, Sst3, Sst4, and Sst5; left) and the boxplot showing the average module scores in the amygdala (right). Each dot represents a mouse in each group. The average module scores of Sst neuronal subclasses tended to decrease specifically in the amygdala after administration of NK cell supplement in 5xFAD mice. Interestingly, the Sst neuronal signatures, which were increased in expression in the amygdala of 5xFAD mice, were decreased to the expression level in wild type mice by NK cell supplements. In contrast, the module score was not different between no treatment and anti-CD4 antibody treatment groups while there were difference between no treatment and NK cell supplement groups. (B) Spatial pattern of the signatures of state-specific glial cells (aging astrocyte and reactive microglia; left) and (C) immune cells (monocyte and plasmacytoid DC), and the boxplot showing the average module scores in the white matter (right). The expression of state-specific subtypes of glial cell and immune cell signatures, which showed a significant increase in 5xFAD mice compared to wild type mice, tended to slightly decrease in the white matter after anti-CD4 antibody treatment. Considering that NK cell supplement showed no appreciable differences in glial cell and immune cell signatures, anti-CD4 treatment effects on these cell subtypes of state-specifics looked real. Expression, however, did not decrease to the expression observed in wild type mice. In summary, NK cell supplement and anti-CD4 antibody treatment affected different state/type-cell signatures and brain regions, respectively. (aCD4: anti-CD4 antibody; WT: wild type; TG: 5xFAD mice; WT_NK: NK cell-treated wild type; WT_aCD4: anti-CD4 antibody-treated wild type; TG_NK: NK cell-treated 5xFAD; TG_aCD4: anti-CD4 antibody-treated 5xFAD; Sst: Somatostatin; DC: dendritic cells; NK: natural killer)

### Regional cell-type/state specific transcriptome changes of 5xFAD model mice compared with wild type after intravenous anti-CD4 antibody treatment

Three mice in anti-CD4 antibody treatment group were selected and their coronal brain section were plated on the slide of Visium. The frame of sample distribution on the quadrants of each Visium slide was the same as above for NK cell supplements treatment. Further analysis of transcriptomes per spots and spatial clustering and designation of transcriptomes to the around 3,000 spots were the same, too.

Among major brain cells, state-specific glial cells, such as aging astrocytes and reactive microglia, which showed a significant increase in 5xFAD compared to wild type mice, showed a slight decrease exclusively in the white matter after administration of anti-CD4 antibody in 5xFAD mice (**Figure 5B**). In the gene combination of aging astrocytes, the expression of *Lgmn, Gsn, Mt1, Fcrls,* and *Hexb* was noticeably decreased in the white matter after anti-CD4 antibody administration in 5xFAD mice, and expression decreased slightly in the other regions (**Supplementary Figure 14C, D**). In the reactive microglial signature, *Cst7* showed a decreased pattern throughout the region, while other genes, such as *Spp1, Cd9*, and *Axl*, showed decreased patterns only in the white matter (**Supplementary Figure 14C, D**). Interestingly, the CD4 T cell signature tended to decrease slightly in the deep and upper cortex (striatum) after anti-CD4 antibody treatment (**Supplementary Figure 15B**). However, mature neuronal signatures showed typical and very similar patterns of distribution between 9 major brain regions and within each region regardless wild type, 5xFAD, wild type with anti-CD4 antibody treatment and 5xFAD with anti-CD4 antibody treatment (**Supplementary Figure 16A**). No significant differences in other types of glial cells, including white matter-associated astrocytes, reactive astrocytes, plaque-associated microglia, homeostatic microglia, and oligodendrocytes, were found meaning all the 10 wild type mice and all the 11 5xFAD mice could not be differentiated between no treatment and anti-CD4 antibody treatment (**Supplementary Figure 16B, C)**. The difference between wild type and 5xFAD sustained while not revealing any effect of the anti-CD4 antibody treatment.

Brain rare cells, resident or infiltrating, made distinction between no treatment group and anti-CD4 antibody treatment group. The expression level of monocyte and pDC, which showed a significant increase in 5xFAD mice compared to wild type mice, tended to slightly decrease only in the white matter after anti-CD4 antibody treatment (**Figure 5C**). In both monocyte and pDC signatures, the expression of *Tyrobp* was dramatically reduced throughout the region, whereas *S100a4, S100a10,* and *Mal* in the monocyte signature showed decreased expression patterns exclusively in the white matter after anti-CD4 antibody treatment. Reduction in the expression of individual genes by anti-CD4 antibody was mainly identified in the white matter region (**Supplementary Figure 14C, D**).

While NK cell supplement showed no appreciable differences in immune cell signatures, the fact that anti-CD4 antibody treatment showed effects on subtypes of immune cells should be noteworthy. However, the expression level was not decreased yet to the expression level observed in wild type mice.

### Methods to scrutinize spatial, cell-type, cell-state specific changes upon the platform of setting norms and characterization of abnormality of a test mouse

As spatial distribution is critical for characterization of a new mouse specimen for their status of normalcy, pathology, and response to any therapy. The specimen can be on every section, but with on or a limited number of coronal sections (per monkey in a report of Chen et al.^55^) or sagittal sections. To acquire representative information regarding mouse groups, we combined multi-individual mouse sections to yield the apparently correct spatial segmentation. Each region was then prepared to present their norms for various scores for cell types, cell states and response to the tentative immunomodulatory drugs. We tried to establish methods to reveal the regional cell-type/state specific norms and their probable changes by drug intervention. Setting-up norms for normal controls in wild types, age of interest, and influence of drug treatments and determination of an individual mouse of their disease states (5xFAD of certain age of diverse behavior/Aβ abnormality, P301L of no behavioral/pathological abnormality) and the therapy effect by the treatments (anti-CD4 antibody or NK cell treatments).

Cell types should have been annotated to the then-best knowledge of the scientific community based on the resource reports in the literature up to the date of this investigation run by trials-and-choice; for example, for neurons and neuronal subtypes, Hodge et al’s report^75^ was adopted as is or after NS Forest^59^ to define 20 GABAergic neurons and 12 glutamatergic neurons. Available data were downloaded from specific sites or supplementary tables of each report. Thus, for example, Scng-VIP neuron subtypes by more recent report by Bugeon et al.^57^ were ignored but later can be reanalyzed with the current Visium data, by specifying their markers along the regions segmented using RPCA of Seurat 4.0. For astrocytes and microglia, especially, reactive state signature was surveyed by scrutinizing the counts of transcriptomics upon each spot according to the reports by Keren-Shaul et al.^88^, Friedman et al.^90^, Grubman et al.^92^ and others. Co-expression of the same transcriptome by astrocytes and microglia such as *Apoe, Gfap, Tspo* and others were removed from the tentative marker gene combination. The same procedure was done for astrocytes and oligodendrocytes, or microglia and oligodendrocytes. Oligodendrocytes and their lineage cells did not have ‘reactive’ transcriptome signature. Signature transcriptomes between reactive and homeostatic microglia (and also astrocytes) was also surveyed for their conjoint expression between both states. Homeostatic transcriptomes were designated to exclude the signature of reactive transcriptomes and vice versa, referring to the literature reports by Prinz et al.^106–112^, and Kim et al.^105^, so that *Sall1* and *Hexb* took the role of counting the number of microglia as each microglia express these genes constitutively while assuming that these transcriptomes did not increase in quantity per microglia when reactive^105,111^. As spatial transcriptomic imaging using Visium yielded linear (semi)quantitative (due to log1p transformation for further processing using Seurat 4.0)) metric, no fractional presentation was adopted to disclose that homeostatic microglia were increased in quantity of signature per spot in 5xFAD mice compared with wild type or P301L mice.

Quantification was performed for 9 regions (hypothalamus, thalamus, deep cortex, white matter, upper cortex, hippocampus, amygdala, piriform area, and striatum) for major cell types including neuron subtypes, reactive and homeostatic glial cells, astrocyte subtypes, oligodendrocyte lineage cell subtypes and finally rare resident/infiltrating immune cells. For regions, for example, white matter-associated astrocytes or white matter-localized microglia were quantified and correlated with white matter-localized oligodendrocyte lineage subtypes. Dense and thus intense quantity of mature oligodendrocytes could be compared among mice groups. On the contrary, diffuse and sparsely distributed rare immune cells, three compartments by Dominguez Conde et al.^71^, four types by Xiemerakis/NSForest^59,64^, and 12 types by Schafflick et al.^68^. Tissue resident macrophages by Dogra et al.^113^ and Eraslan et al.^69^ were also quantified for its intensity to yield the difference for each region among mice groups. Wilcoxon rank sum test was used to determine the significance between pair of groups (uncorrected p value or corrected by three for paired groups comparison). Beyond group comparison, an anomaly detection procedure (or confirmation of normalcy meaning no difference found on any regional, cell-type, cell-state specific signature by density per region) was performed for each mouse.

Once a region and cell-type and its state were found, we performed DEG analysis to find the transcriptomes of interest in each brain region. Then, the association of discovered genes with biological pathways was examined using an over-representation test based on evidence-driven databases. It was to determine of the significance of the found transcriptome designating their functional role in pathology (amyloid pathology or tauopathy, glial cell dysfunction, etc.), physiology (aging) and their participation in the response to the tentative immunomodulatory therapy. Considering the diversity of behavior improvement after anti-CD4 antibody and NK cell supplements treatments, and also the un-paired nature of Visium study, we could only detect the treatment effect (and non-effect) of transcriptome signature of regional cell-types/states upon treatment per individual. Transcriptomes of marker gene combination which we used were all checked for their individual transcriptome to elucidate key transcriptomes for specifying type/state characteristics or therapy effects. We also tried to assess their individual contribution for this specification to find one, two or more distinctive transcriptomes to predict their presence in each spot. This means that curation by operator in addition to the readymade Wilcoxon, logistic, or NSForest methods was used at least as two steps, first to choose a seemingly optimal combination ruling out cross-expressing, background, or confounding genes and then at last to find the succinct combination of transcriptomes for cell-type/state annotation or if any, the sole transcriptome (Supplementary Figure 2). The above pipelines for dissecting cell-type and cell-state specific regional transcriptomic changes can be readily implemented with our in-house STquantool application which facilitates visualization of spatial gene expression and enables quantification across multiple transcriptomic datasets.

## Discussion

In this investigation, we tried to use ST for their superiority over scRNAseq/snRNAseq to localize the specific transcriptomic signature of cell-type or cell-state to almost 5 thousand spots, among which 3,000 or more spots harbored the brain tissue, either coronal or sagittal sections. Before going further to use this transcriptomic signature to disclose the effects of novel but unproven neuromodulatory treatments, we trimmed how to use this Visium-based ST imaging to elucidate regional, cell-type/state specific changes. How to choose an or two optimal transcriptomic marker combinations among so many possible combinations was adjusted to yield the best contrast between cell types/subtypes using the literature resources and our inhouse method of curation. Simple and easy method to sort out the candidate transcriptomes was set up to assure that we did the job to find the best cell type/state annotation methods, for either abundant brain cells or rare immune cells. Separation of 4 or more major brain cells and their subtypes with transcriptome combinations or definition of rare immune cells for their exact propensity and distribution/location were the challenge and obstacle. Stahl et al.’s^47^ suggestion of counting the transcriptomes per spots using the original ST Visium methods, Tirosh et al.’s^42^ approach to generate cell signature scores based on the curated marker genes, and comparing them between mouse groups with genuine or sham treatments worked well for this endeavor. We overcame the problem of high dependence of this endeavor on the choice of tentative marker gene combinations varying upon the diverse input data derived from the preliminary DEG studies using single-cell data of brain tissues^67,68,106^ or even other tissues than brain tissues. Assessing the sophisticated use of the public database and scrutinizing the individual transcriptomes visually by us the operators (neuroimaging experts) were essential. Curation by operators would look heuristic at best and be sure to subject to operator arbitrariness, however, eventually were the key step to enhance the authenticity of the observation of large number of cells (2 to 10 per spot) admixed in spots and also a dozen specimen from individually different but syngeneic mice. From the neuroimaging perspective, integrated single spot imaging (100μ x 100μ x 10μ) containing average 5 cells (2 to10) in each unit domain did not have significant batch/individual variation effect to confound further analysis as we observe dozens/hundreds of spots at the same time and the batch effect was corrected during the sample integration. With this visual investigation, we soon came to be confident that spatial transcriptomic brain spot imaging with visual assessment and its quantitative analysis using the framework of voxel (spot) imaging of mouse/human brains was suitable for the effect evaluation of certain drugs/treatments for disease-course modification in dementia mouse models.

Transcriptomic signature of brain cells could clearly segment every section of mice, regardless of disease status under no treatment or probably effective treatment, taking advantage of 22,000 or more transcriptomes per cells so as to identify the cell-type/state with thousands of variable transcriptomes. Unlike functional neuroimaging such as functional magnetic resonance imaging (fMRI) or positron emission tomography (PET), which needs co-registration and segmentation considering individual variation for further analysis, the segmentation of neuroanatomical entities on Visium could be performed without any more assumption, except the one that functional regional entities could be determined by transcriptomes belonging to spots and their conglomerated spots making explicit functional regions. An eccentric case of regionally remote but similar transcriptome composition was observed, needing to say that cortical amygdala and subcortical septal lobe were categorized as the same cluster on sagittal section, but excluding this exception, all the other spatial clusters were within the expected anatomical border definition (https://connectivity.brain-map.org/3d-viewer?v=1&types=IMAGEPLANE&IMAGEPLANE=imageplanes). Thus, spatial or regional representation of characteristic changes related with pathology and treatment response could then be described and quantified. Finding marker gene combination to define the spots to belong to certain functional region of interest, then, became the problem to be solved by finding the optimal or best combination, which would be appropriate and succinct.

Determination of best annotation of neurons and other major brain cells was initially dependent upon the previous reports mainly derived from non-spatial scRNAseq/snRNAseq analysis^63,73,79,114^. On these previous studies, spatially expected designation of cells was suggested as success of cellular clustering, raising concern that there was no gold standard information regarding their true location, nevertheless, this floated cells’ clustering allocate the cells to their probable location and thus proposed to be correct. ST brain imaging obviated this concern. In ST imaging, however, there still remains two major problematic ambiguity for spatial clustering and cell type/state identification per spot. The first one was spatially agnostic annotation by transcriptome signature, which was tried to solve by sampling regions of brain such as posterior isocortex, hippocampus (or hippocampal formation), striatum, thalamus and hypothalamus etc. in the reports of Saunders et al.^65^ or Chai et al.^115^. This problem was easily solved by ST imaging using Visium of 3 to 5 thousand of spots which allows capturing transcriptomic changes across the broad region of the brain. Now the imaging with the resolution of 100μ x 100μ on 2D is available allowing easy segmentation, whose difference from fMRI/PET is that the huge multiplexing capability of ST brain imaging allows almost infinite times of reanalysis using combinatorics. The second one was cell/state identification per spot was done by using transcriptomic signature of the marker gene combination determined by previous DEG studies using detached and sometimes surface-marker-sorted brain cells. When scRNAseq/snRNAseq was used for detached brain cells to determine the effect of drug/treatment on those brain cells, lack of spatial localization was the major hurdle blocking the understanding of the role of any treatment. In situ hybridization of immunohistochemistry complemented transcriptomic signatures of global/regional brain in this job, but without reassuring results to explain the therapy effect. ST brain imaging solved this problem. As is shown in this study, ST brain imaging is equipped with the expression profile per spot for the entire genes of the individual cells localized on each spot, the data could be analyzed in an unsupervised fashion without any assumption or in appropriate cases by using a priori knowledge derived from the literature resources of scRNAseq/snRNAseq. Considering the challenges and difficulties to overcome these problems, we streamlined the use of visual reading by expert operators called as curation. As many steps as practical curation could have been performed, however if any, then minimal as possible, initially to exclude non-specific and cross-expressing transcriptomes between major cells, and finally to exclude cross-tissue, stromal cell-dependent and confounding background signatures. Curation should have been better to be based on individual transcriptomic features of any types of cells for their association with disease states or drugs/treatments responsiveness.

To tackle these problems, we asked how we could use the individual mouse ST brain imaging data to determine the disease states, which are variable as much even in syngeneic animals and also the variable treatment responses affecting major and rare brain cells. Taking advantage of the automatic segmentation results for groups of individual mouse specimen, irrespective of section planes and stereotaxic coordinates, we tried to individualize the transcriptomic feature of each individual specimen compared with the norms we constructed. Comparing regional, cell type/state specific transcriptomic signature using visual and quantitative decision of an individual mouse with the norms was done. This analysis method added the individuation-based interpretation of animals for their behavior correlates. We could obtain and reproduce the wide variety of behavior metric, which is in this study, the alternation score on Y-maze, and this Y-maze alternation scores of 7-month-old wild type mice ranged widely as well as of 5xFAD mice, but those of 8.5-month-old wild type mice converged with smaller variation to the lower values meaning commonly poorer performance at this age even in wild type mice. After anti-CD4 antibody treatment, variation sustained with slight improvement of their average scores (**Figure 4B**). After NK cell supplements treatment, variation also sustained with slight improvement of their average scores too (**Figure 4B**). We assumed that these behavior variabilities are the keystone for proving the feasibility of a tentative novel immunomodulatory treatments and that we would find that the mouse behavior scores concord with the transcriptomic signature^57^. NK cell-treated 5xFAD mice with higher Y-maze alternation scores definitely showed that their amygdala GABAergic Sst neuronal subtypes decreased in intensity (**Figure 5A**). This decrease (or increase, if any) did not prove the efficacy of NK cell supplements treatment on 5xFAD mice, but definitely disclosed that transcriptomics of the neuronal subtype of that region were correlated with the degree of behavior impairment. More importantly, this meant that so many other neuronal subtypes, other homeostatic or reactive glial cells and their subtypes, did not show any change in intensity over all the regions examined on these sections despite the improved behavior score. Anti-CD4 antibody treatment recapitulated only a slight decrease of specific immune cell signatures in the white matter, but beyond this finding, no other discovery of the drug-responsive transcriptomic changes on every region, on any cell-types and cell states was found. This was even on individual interpretations both visually and quantitatively for each mouse (**Figure 5B** **and Supplementary Figure 15**). We could say that anti-CD4 immunoglobulins did not affect transcriptomic signatures of brain major cells (on this single coronal section), and such was also the case with rare immune cells. Due to the lack of Y-maze score measures of the anti-CD4 antibody-treated wild type and 5xFAD mice, behavior correlation could not be reported here.

The interpretation of rare immune cell signatures for the localization of immune cells presented different challenge from the major brain cells. The first one was to remove the background effects by major brain cells. Homeostatic and reactive microglia and their co-expressed transcriptomes between microglia and infiltrating monocytes were the major challenge but were easy to remove, and astrocytes and oligodendrocytes followed as the reactive glial cells expressed the same/similar transcriptomes. Double-checking the unique transcriptomes and their combinations were tried with the data by Ximerakis et al.^64^ and Schafflick et al.^68^ based on brain tissue studies. The study by Schafflick et al.^68^ used the cells sorted by FACS for CD45 (gene *Ptprc*) positivity and thus could not allow us to remove the co-expressed transcriptomes of ribosomal protein transcriptomes for lymphoid cells, which if removed, would have enabled correct classification of myeloid and lymphoid cells among major brain cells in terms of intensity and distribution. Nevertheless, visual/manual curation by surveying individual transcriptomes helped to remove absurdly intense and unrealistically distributed transcriptomic signature. When we used only the data of Schafflick et al.,^68^ we could not correct the inappropriate signature for B and T cells compartments even after NSForest application to their data. The data came to look realistic after we adopted cross-tissue data and visual curation upon the two reports by Eraslan et al.^69^ and Dominguez Conde et al.^71^. DEG data with arbitrary threshold of 2.0 higher or −2.0 lower of log fold change ratio (LFCR) for MHC+ infiltrating immune macrophage (Mϕ) or LYVE+ infiltrating integrity Mϕ produced 200 or more or 100 or more transcriptomes, respectively. We needed to remove, upon visual curation, 30% or 20% of transcriptomes to annotate the infiltrating monocyte-derived Mϕ. Infiltrating Mϕ and border-associated Mϕ^67,68^ should have been differentiated but not possible due to the lack of clear understanding between two cell types in the literature and sparsity of the cells of both types. Tissue-resident and effector memory cells were traced with the transcriptomic signature by Dominguez Conde et al.^71^. As these authors included variety of tissues (unfortunately brain not included) and stromal tissue-specificity were considered as possible confounders in common for every tissue and thus as expected, they yielded the signature for three compartments of T/ILC, B cell and myeloid compartments. Of course, the types/subtypes of classically well-known immune cells belonging to these three compartments represented well the rare immune cells which would have originated from the bone marrow. We could find the difference of the intensity and distribution of three compartments in the brain sections between 5xFAD mice and wild type mice (**Figure 3**). Drug/treatment effects should have been disclosed with this comparison, but we just say that further investigation is warranted with larger number of mice to avoid noise confounder to hide or spuriously render probable false negative/positive results regarding the effect or no effect of any tentative immunomodulatory treatments (**Supplementary Table 5 and 6**).

Ultimate objective of using ST brain imaging with its visual and quantitative analysis is to designate convincingly the target cells with regional localization, either major or rare, either brain parenchymal/stromal or rare immune cells, either resident or infiltrating immune cells and their states of homeostatic/reactive, and target genes with significant contribution to pathologic changes of cells/regional tissues and their response to the effective or ineffective treatments. More important was that we could be sure that the unfound cells and transcriptomes were innocent, meaning that they were not affected by the test-trial of a novel immunomodulatory therapy. For neuroimmune interaction of disease process or by disease modifying drugs, we now know that skull BM communicates with dural sinus and peri-sinus regions, dural lymphatics as well as across ABC and CSF and thus perivascular spaces and ISF, or totally different and unique route via capillary endothelium and stroma of choroid plexus, and choroid epithelium despite its tight junctions as well as brain blood vessels’ microvascular endothelium despite its tight junctions. Once immune cells from three compartments of T/ILC, B cells and the myeloid are infiltrating to the brain parenchyma, dynamically changing along the aging or disease process (in 5xFAD or P301L), they can respond to systemic immunomodulatory drugs treatment directly or at least indirectly. The ratio of T cells, for example, average 4 /mm3, to neurons (90,000/mm3) or microglia (6,500/mm3) suggest that few immune cells could change the response of major brain cells in their significant way of transcriptomic signature. How the signals are transferred and/or translated from systemic administered anti-CD4 immunoglobulins or NK cell supplements are to be investigated further. In this study, the study scheme and analysis methods were proposed to be applied to use ST brain imaging for investigating the impact of a novel tentatively disease-modifying treatments upon neurodegenerative diseases, to elucidate whether regional brain cell-type/state specific changes in the entire transcriptomes per spot/region/cells of the brain or immune system would respond or not. Comprehensiveness and resolution would be bettered much with the more novel technology^54,55^ which will be available soon in many institutions like Visium methods^47^.

## Methods

### AD models in different ages

Three-month- and 7.5-month-old male 5xFAD mice (Tg6799; on a C57/BL6-SJL background) containing five FAD mutations in human APP (the Swedish mutation, K670N/M671L; the Florida mutation, I716V; and the London mutation, V717I) and PS1 (M146L/L286V) and wild type mice were used for spatial transcriptomic brain imaging data. Six- and seventeen-month-old male tau P301L mice (MAPT P301L mutations; on a FVB/N background) were used. Mice of all strains were raised in a laboratory cage with controlled temperature and humidity and on a 12 h light-dark cycle with no restriction of standard feeding and water drinking. All experimental protocols and animal usage were approved by the Institutional Animal Care and Use Committee at Seoul National University.

### Peripheral CD4 T cell blockade in 5xFAD AD model

Anti-CD4 antibody (0.5 mg/mouse; Bio X Cell) was intravenously injected to the 6.5-month-old 5xFAD and wild type mice according to the groups. The samples of different tissues were obtained after a month. Coronal sections of brain samples were used for spatial transcriptomic brain imaging and analyzed.

### Administration of NK cell supplement in 5xFAD AD model

NK cells were expanded for 7 days after the isolation of NK cells from BALB/c mouse spleen. NK cells (2 x 10^6^ cells/mouse in saline) were intravenously administrated once a week for a total of five times to the 6.5-month-old 5xFAD and wild type mice. The brain samples were obtained after five weeks and used for spatial transcriptome data.

### Spatial gene expression library construction

Brain hemispheres were prepared in frozen block using OCT compound (Sakura) and cryosectioned to 10 μm of coronal and sagittal sections. According to the manufacturer’s protocols using Visium Spatial Tissue Optimization slides (10X Genomics) permeabilization time was optimized to find the time of 12 minutes. The brain sections were methanol-fixed, hematoxylin and eosin (H&E)-stained and imaged on a TissueFAXS PLUS (TissueGenostics). The slides were merged into a picture of the whole brain using TissueFAXS imaging software. Then, the sections were permeabilized and processed to obtain cDNA Visium Spatial Gene Expression libraries according to the manufacturer’s protocol. To verify the size of PCR-enriched fragments, the template size distribution was checked using high sensitivity DNA assay (Agilent Technologies 2100 Bioanalyzer).

### Generation of count matrix

The libraries were sequenced using HiSeqXten (Illumina) with a read length of 28 bp for read 1 (Spatial Barcode and UMI), 10 bp index read (i7 index), 10 bp index read (i5 index), and 90 bp for read 2 (RNA read). Raw FASTQ data and H&E images were processed by the Space Ranger v1.1.0 (10X Genomics) pipeline for the gene expression analysis of Visium Spatial Gene Expression library data. Illumina base call files from the Illumina sequencing instrument were converted to FASTQ format for each sample using the ‘mkfastq’ command. Visium spatial expression libraries were analyzed with the ‘count’ command. Image alignment to predefined spots was performed using the fiducial alignment grid of the tissue image to determine the orientation and position of the input image. Sequencing reads were aligned to the mm10 reference genome (mm10-2020-A) using STAR (v2.5.1b) aligner. Gene expression profiling in each spot was performed with unique molecular identifier (UMI) and 10X barcode information.

### Spatial transcriptome data: Integration and spot clustering

The generated gene counts were normalized using ‘LogNormalize’ methods with the scale factor 10,000 in Seurat v.4.1.1. The top highly variable genes (n= 2,000) were then identified using the ‘variance stabilizing transformation (vst)’ method in Seurat. The log-normalized count matrix was scaled, and (reciprocal) principal component analysis (PCA) was performed for dimensionality reduction using the top highly variable genes. The number of RNA counts for each spot in the scaling process. An integration was performed for multiple spatial datasets before spot clustering. A set of anchors were discovered between the datasets using ‘reciprocal PCA’ and normal mice (wild type) were used as a reference during integration. The anchors were utilized to correct the count matrix in each spatial data. Then, the corrected counts were integrated and were scaled and PCA was performed. For spot clustering, a shared nearest neighbor (SNN) graph was constructed, and the graph-based clustering was performed based on the Louvain algorithm. The resolution was set to 0.15. The anatomical location of each cluster was visually identified by comparison with the Allen Mouse Brain Reference Atlas (https://mouse.brain-map.org/static/atlas). In all analyses, the count matrix normalized in the same way as above was used. For visualization, dimensionality reduction was performed using UMAP on the top 30 principal components. The spots with gene expression data were analyzed with the Seurat package (ver 4.1.1). For analysis of AD models in different ages, 63 spatial transcriptome datasets with 32,885 genes in common were integrated. All analyzes used data integrated and clustered by the above methods.

### Differential gene expression analysis

MAST^96^ in the Seurat function was used to perform differential gene expression analysis. Differentially expressed genes were extracted from the comparison of wild type and 5xFAD mice in each cluster. The cutoff of significantly different genes was false discovery rate (FDR)-adjusted p < 0.05 and log FC > 0.25.

### Gene ontology analysis

Gene ontology analyses were performed with clusterProfiler^116^, which supports statistical analysis and visualization of functional profiles for genes and gene clusters. The Benjamini-Hochberg (BH) adjusted p-value was indicated. The ‘Enrichplot’ and ‘igraph’ package were additionally used.

### Marker panel selection and curation

For gene combinations identified in other references, a panel of markers was constructed from individual gene curation. Genes that were not present in our spatial transcriptome data were excluded. Also, Necessary and Sufficient Forest (NS-Forest) version 2^59^ was applied to the publicly available single-cell datasets that contain the cell type annotation information. The genes were scored according to binary expression profiles in a specific cell type compared to other cell types. Then, based on the random forest algorithm, the minimum gene set which best describes the given cell type was searched. The curated gene sets are listed in **Supplementary Table 4**.

### Cellular signatures of reference data

The marker panel, a gene set that best represents a certain cell type, was determined based on a variety of reference papers. Then, the signature score of each cell type was computed on the spatial transcriptomic data by utilizing the AddModuleScore function in Seurat^44^ with default parameters. The score in each spot was spatially mapped to the tissue using the SpatialFeaturePlot function and the spatial distribution pattern was identified. The average of the signature scores was calculated in a certain region of interest and the values were compared between groups. Multiple comparison correction was performed with the Bonferroni method after Wilcoxon rank-sum test and the cutoff for the adjusted p-value was 0.05.

### Statistical analysis

For the spatial transcriptome data, plots in R were created either with the ggplot2 R package or Seurat modified by custom codes for data visualization. All p-values reported in this study were adjusted by FDR (for DE analysis using MAST) and Wilcoxon rank-sum test and Bonferroni method (all other analyses).

### Development of an application to visualize and quantify ST datasets

R shiny-based application named STquantool was developed to comprehensively analyze ST datasets to explore cell type and cell state-specific regional change in the wild-type, 5xFAD, and treatment mouse models. The application allows users to easily load and integrate the multiple ST datasets, and visualize the spatial expression of genes and cell type scores based on Seurat^74^ and shiny running on R (ver. 4.1.1). One of the key features of STquantool is that it facilitates the curation of cell-specific marker combinations by sorting out key genes based on NSForest^59^ algorithm and finalizing the markers by visually assessing the spatial expression patterns. As an adjunct, the cell type decomposition method, CellDART^46^ can be implemented to find the spatial distribution patterns of major cell types constituting the brain tissues. Moreover, the spatial patterns of the cell scores and cell fraction can be quantified and statistically analyzed with STquantool. Lastly, the gene level transcriptomic alterations between the mouse groups can be explored by performing the DEG analysis provided in the application. Then the functional implications of the selected genes can be represented by gene ontology (GO) and Kyoto Encyclopedia of Genes and Genomes (KEGG) terms^117–119^. The suggested platform was packaged and can be readily installed from the GitHub (https://github.com/bsungwoo/STquantool.git).

## Acknowledgments

We thank Joo Young Park for his technical assistance. The work was supported by grants from the National Research Foundation of Korea (NRF-2017M3C7A1048079, NRF-2020R1A2C2101069, NRF-2022R1A5A6000840).

## Funding

This work was supported by National Research Foundation of Korea (NRF) Grants funded by the Korean Government (MSIP) (No. 2017M3C7A1048079, No. 2020R1A2C2101069 and No. 2017R1A5A1015626).

## Author contributions

EJL: Methodology: Investigation: Visualization: Writing

SWB: Methodology: Investigation: Visualization

MS: Methodology: Investigation: Visualization

HC: Methodology: Investigation: Funding acquisition

YC: Methodology: Investigation

DWH: Methodology: Investigation

DSL: Conceptualization: Methodology: Investigation: Visualization: Funding acquisition: Project administration: Supervision: Writing – original draft: Writing – review & editing

## Competing interests

Authors declare that they have no competing interests.

## Data and materials availability

All data are available in the main text or the supplementary materials. All data, code, and materials used in the analysis are available through a standard material transfer agreement with Seoul National University to academic and nonprofit by contacting the authors.

## List of Supplementary Tables and Figures

**Supplementary Table 1.** Information on brain samples used for spatial transcriptome analysis (Excel).

**Supplementary Table 2.** List of differentially expressed genes identified between 7-month-old 5xFAD and age-matched wild type mice (Excel).

**Supplementary Table 3.** Information on the number of spots per sample in anatomical segmentations identified by spatial clustering (Excel).

**Supplementary Table 4.** List of reference-based gene combinations identified via a workflow-based curation process (Excel).

**Supplementary Table 5.** Summary of the spatiotemporal changes in the progression of amyloid pathology in 5xFAD mice.

**Supplementary Table 6.** Summary of the feasibility test of ST analysis for verifying the mode of action of NK cell supplement and anti-CD4 antibody treatment in 5xFAD mice.

**Supplementary Figure 1.** Spatial transcriptome-based cluster analysis in mouse brain samples of various types and conditions.

**Supplementary Figure 2.** Schematic workflow to identify differences between groups using spatial transcriptome brain imaging data.

**Supplementary Figure 3.** Identification of spatial differences in the distribution of neuronal subclasses between wild type and 5xFAD mice.

**Supplementary Figure 4.** Spatial patterns of individual genes involved in differences in somatostatin inhibitory neuronal signatures in wild type and 5xFAD mice.

**Supplementary Figure 5.** Spatial differences in the distribution of the region- or subtypes-specific signatures of glial cells between wild type and 5xFAD mice.

**Supplementary Figure 6.** Spatial patterns of individual genes involved in signature differences of microglia and astrocytes between wild type and 5xFAD mice.

**Supplementary Figure 7.** Spatial changes in the distribution of brain cell signatures in 3-month-old compared to 7-month-old 5xFAD mice.

**Supplementary Figure 8.** Identification of DEGs and related biological pathways in the 5xFAD compared to wild type mice.

**Supplementary Figure 9.** Spatial differences in the distribution of rare brain cell signatures between wild type and 5xFAD mice.

**Supplementary Figure 10.** Spatial patterns of individual genes involved in differences in type/subtype signatures of rare immune cells between wild type and 5xFAD mice.

**Supplementary Figure 11.** Spatial differences of individual genes specifying brain major or rare cell types between wild type and 5xFAD mice.

**Supplementary Figure 12.** Spatial distribution patterns of specific isoforms of peroxiredoxin representing different major brain cells.

**Supplementary Figure 13.** Display of batch-by-batch samples used for ST brain imaging analysis and their Y-maze results after intravenous aadministration of NK cell supplements in 5xFAD mice.

**Supplementary Figure 14.** Spatial expression pattern of individual genes involved in changed brain cell signatures after NK cell supplement or anti-CD4 antibody treatment in 5xFAD mice.

**Supplementary Figure 15.** Spatial differences in the distribution of NK and CD4 T cell signatures with or without NK cell supplement or anti-CD4 antibody treatment in 5xFAD mice.

**Supplementary Figure 16.** The distribution of brain cell signatures showing no changes in any region after NK cell supplement and anti-CD4 antibody treatment in 5xFAD mice.

**Supplementary Figure 17.** *In vivo* SPECT/CT images and biodistribution of ^99m^Tc-HMPAO-NK cells.

## Notes

### Competing Interest Statement

The authors have declared no competing interest.

