## Supplementary_Figures_Tables for "Spatial transcriptomic brain imaging reveals the effects of immunomodulation therapy upon specific regional brain cells in mouse dementia model"

**Supplementary Materials**

**Supplementary Tables 1-6**

**Supplementary Figures 1-17**

**Supplementary Table 1. Information on brain samples used for spatial transcriptome analysis (Excel).**

**Supplementary Table 2. List of differentially expressed genes identified between 7-month-old 5xFAD and age-matched wild type mice (Excel).**

**Supplementary Table 3. Information on the number of spots per sample in anatomical segmentations identified by spatial clustering (Excel).**

**Supplementary Table 4. List of reference-based gene combinations identified via a workflow-based curation process (Excel).**

**Supplementary Table 5. Summary of the spatiotemporal changes in the progression of amyloid pathology in 5xFAD mice.** Summary of evaluation of major and rare brain cell signatures in ST data of 5xFAD mice at 3 and 7 months of age. Brain region-specifically changed cell signatures were identified as the amyloid pathology progressed. (AD: Alzheimer’s disease; CAM: CNS-associated macrophage; DC: dendritic cells; NK: natural killer)

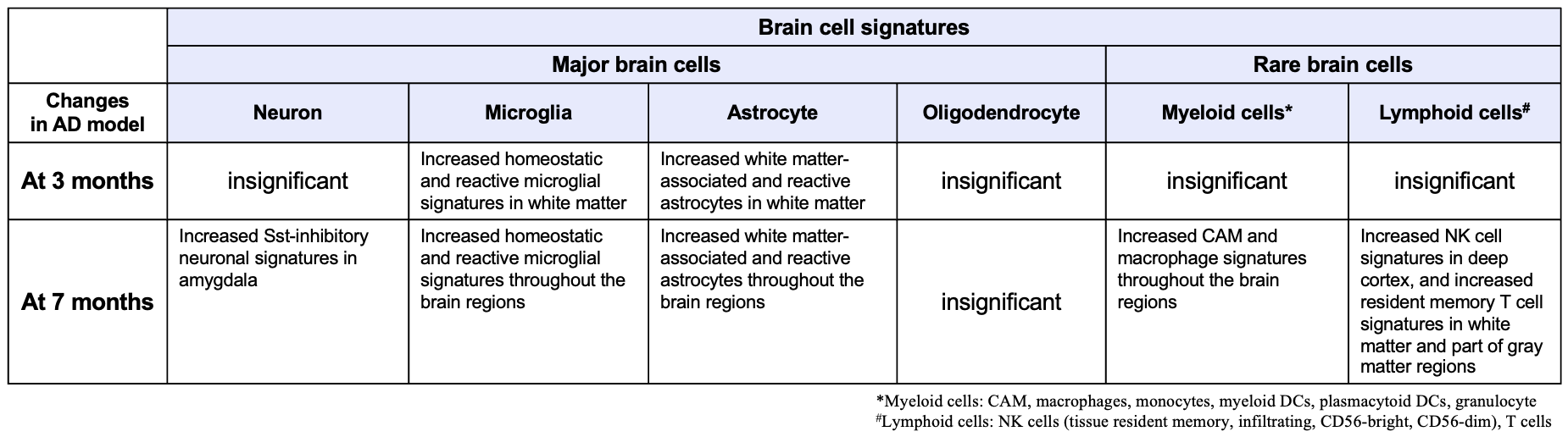

**Supplementary Table 6. Summary of the feasibility test of ST brain imaging analysis for verifying the mode of action of NK cell supplement and anti-CD4 antibody treatment in 5xFAD mice.**

Summary of evaluation of major and rare brain cell signatures in ST data of 5xFAD mice after administration of NK cell supplement and anti-CD4 antibody. Brain region-specific transcriptome changes in cell signatures were identified as the mode of action of therapeutic agents that is associated with behavior improvement. (**ST: spatial transcriptomics**; aCD4: anti-CD4 antibody; NK: natural killer; AD: Alzheimer’s disease; CAM: CNS-associated macrophage; DC: dendritic cells)

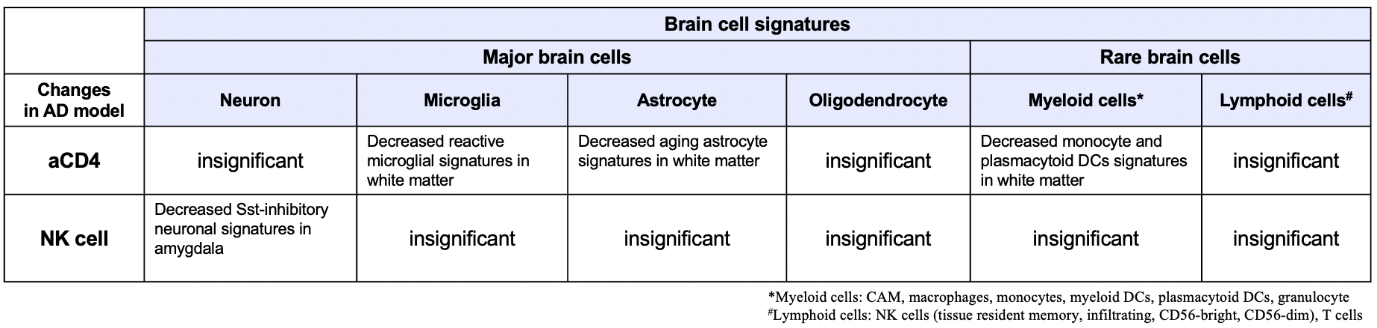

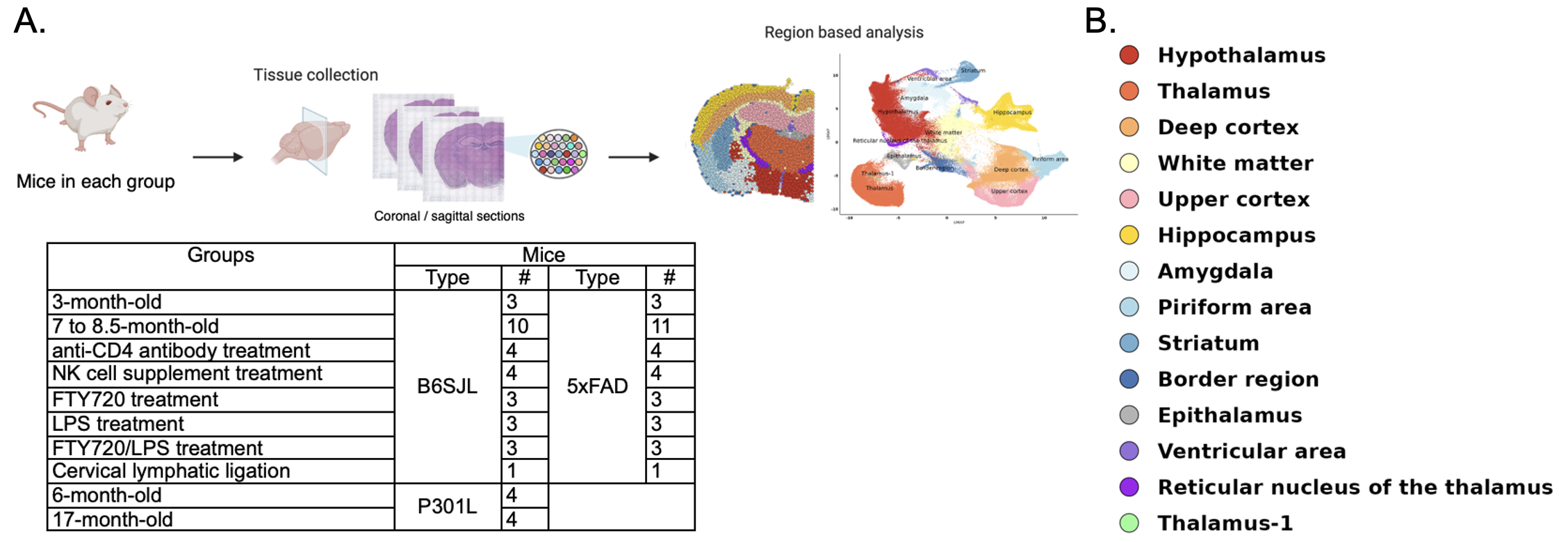

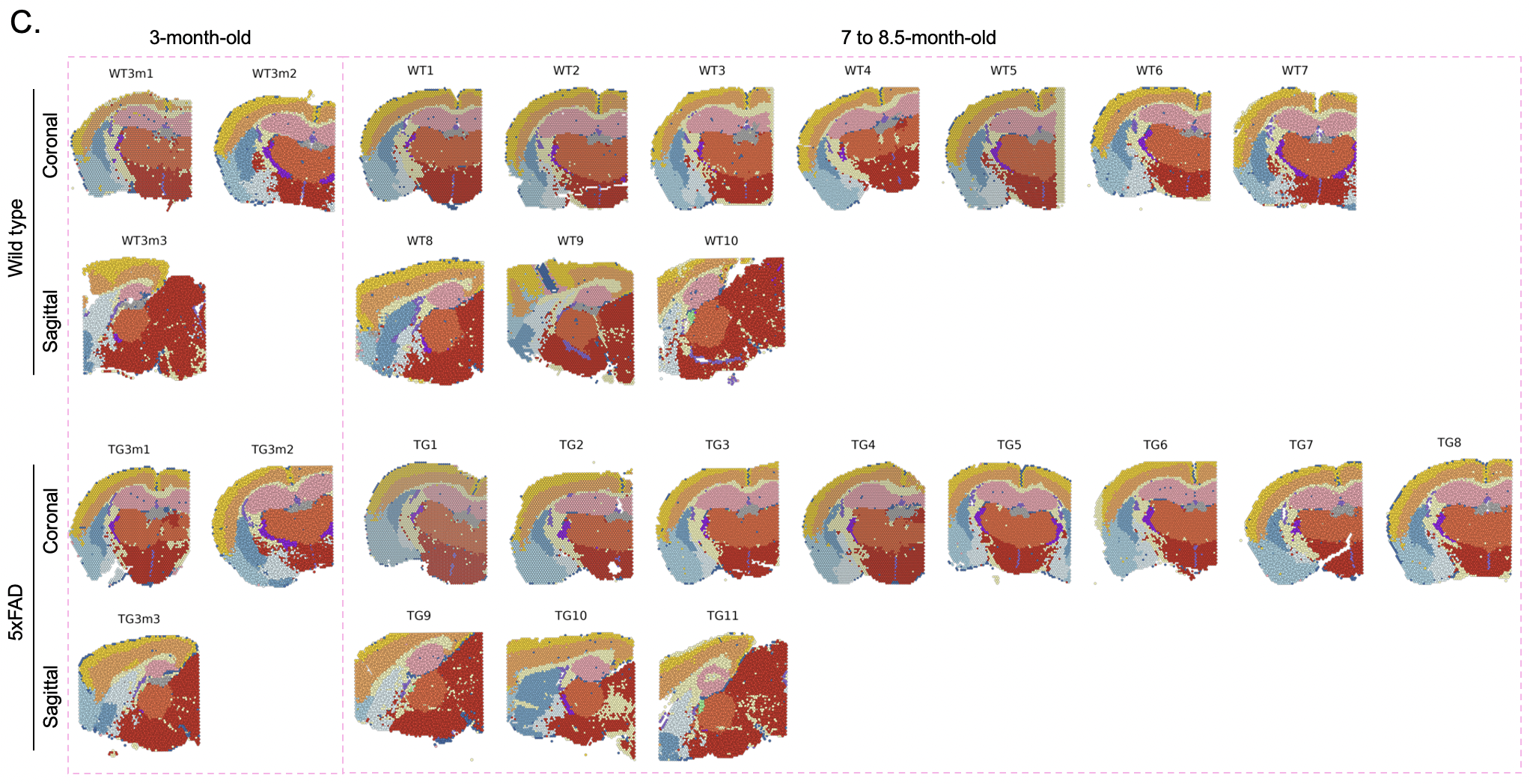

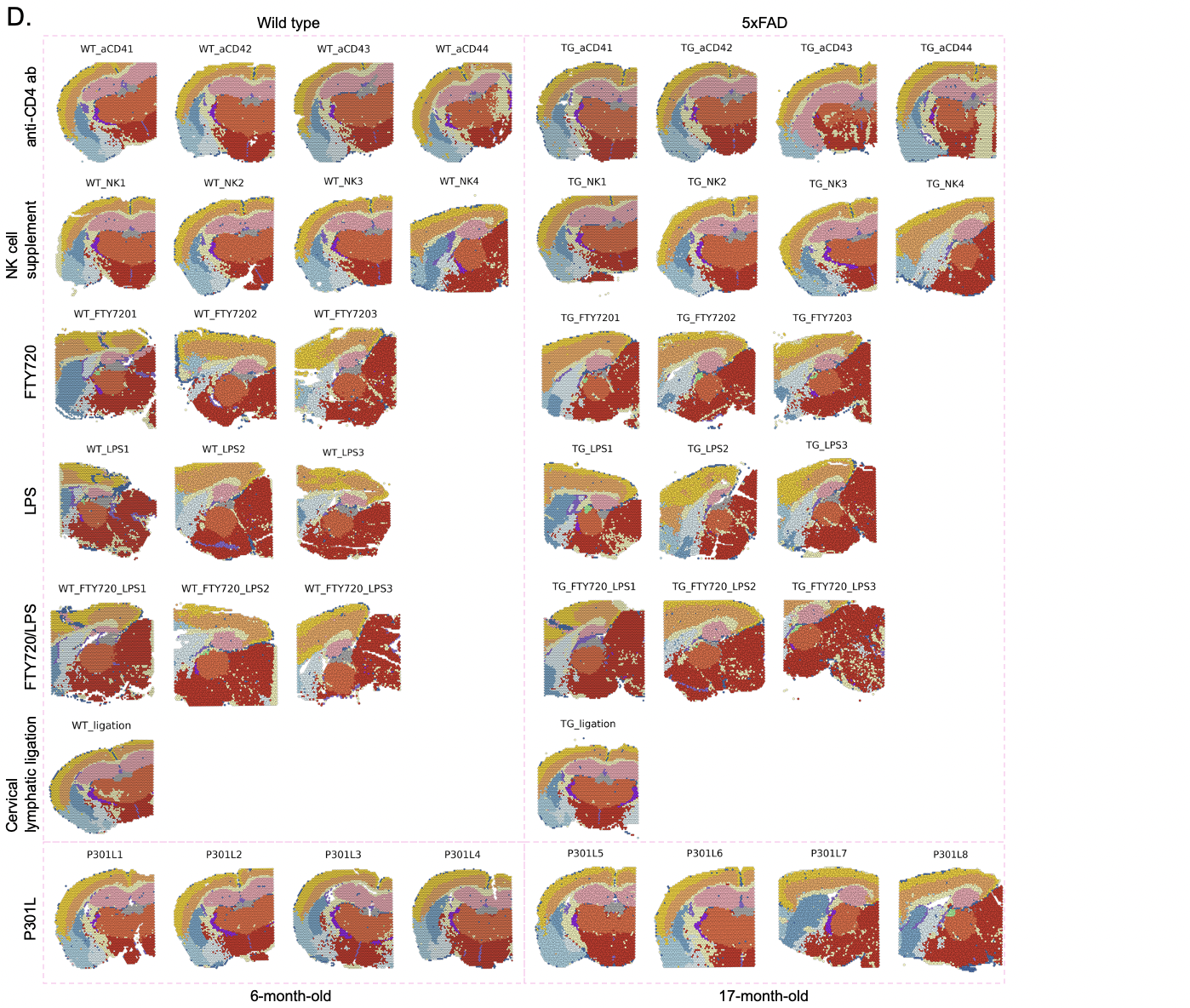

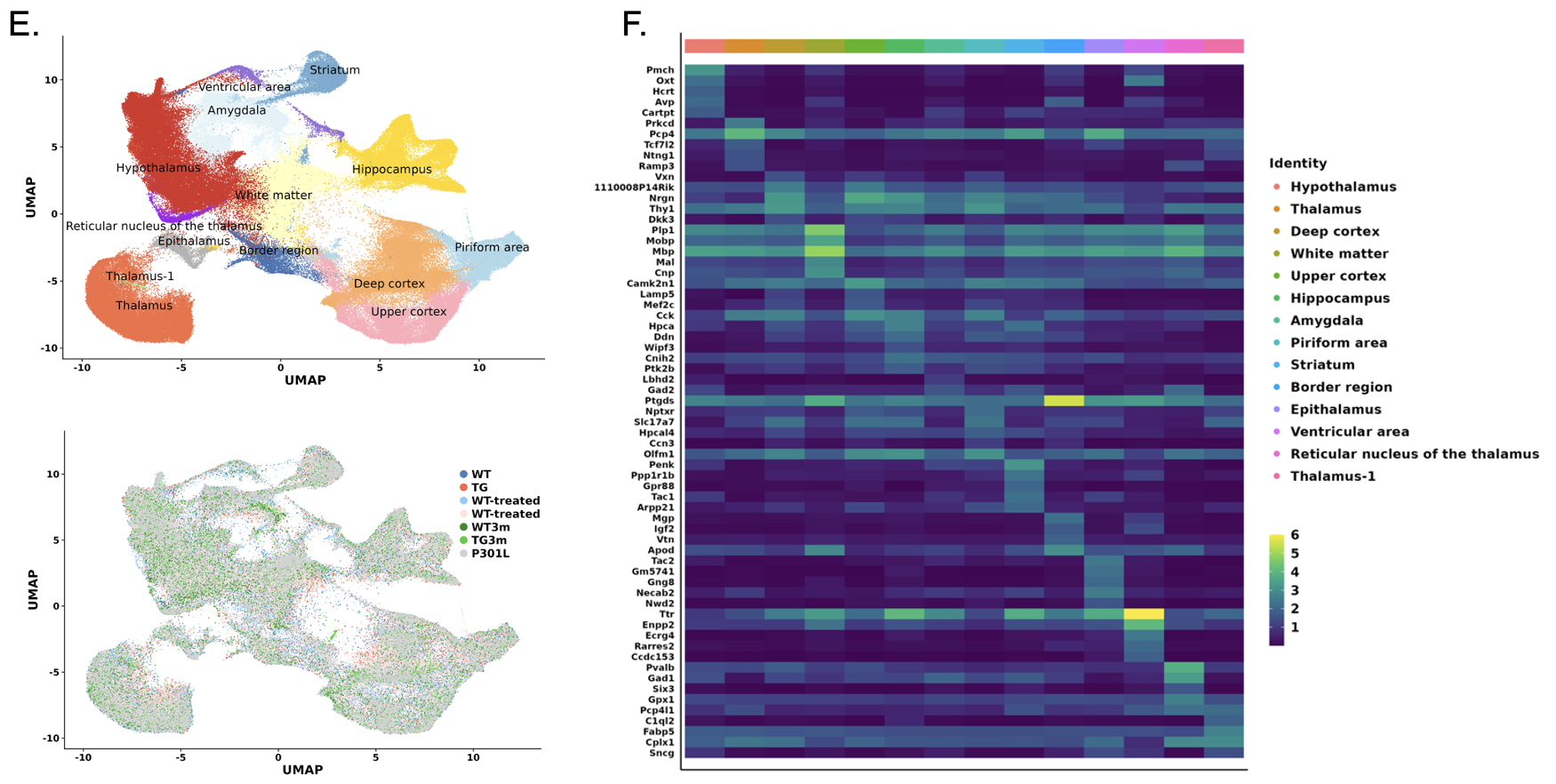

**Supplementary Figure 1. Spatial transcriptome-based cluster analysis in mouse brain samples of various types and conditions.**

(A) Schematic overview of the experimental workflow for spatial transcriptome data from mouse brain samples. The table lists the types and the number of mice used in each group. Coronal or sagittal sections were obtained from the brain tissues (one per each mouse) and region-based analysis was performed. (B) Fourteen clusters were identified by unsupervised clustering of a total of 71 mice in each group. The clusters were colored and annotated based on anatomical brain regions. (C) Spatial cluster images of wild type and 5xFAD mice aged 3 months and 7 to 8.5 months old. (Wild type/5xFAD-3-month-old: 2 coronal/1 sagittal sections; -7 to 8.5-month-old: 7-8 coronal/3 sagittal sections). (D) Spatial cluster images of wild type and 5xFAD mice belonging to the groups of various treatments and manipulations. (Wild type/5xFAD-anti-CD4 antibody: 4 coronal sections; -NK cell supplement: 3 coronal/1 sagittal sections; -FTY720: 3 sagittal sections; -LPS: 3 sagittal sections; -FTY720/LPS: 3 sagittal sections; -Cervical lymphatic ligation: 1 coronal section; P301L-6-month-old: 4 coronal sections; -17-month-old: 4 coronal sections). (E) UMAP of 261,815 spots. The clusters were colored and annotated based on the information obtained in C and D (upper). A UMAP plot was colored to represent each group (lower). The blue dots indicate 10 mice in the WT group, the red dots indicate 11 mice in the 5xFAD group, the sky-blue dots indicate 18 mice in the WT-treated group, and the pink dots indicate 18 mice in the 5xFAD-treated group, dark green dots indicate 3 mice in the 3-month-old WT group, green dots indicate 3 mice in the 3-month-old 5xFAD group, and gray dots indicate 8 mice in the P301L group. Each dot represents the transcriptomic data of each spot. (F) Heatmap showing the distribution of expression levels of marker genes across 14 brain region-based clusters. The top five genes were identified for each cluster. The clusters were well segmented to be compatible with anatomical brain structures through unsupervised clustering in every case including those having various treatments and manipulation. (WT: Wild type; TG: 5xFAD AD model; NK: Natural killer; LPS: lipopolysaccharide; FTY720: fingolimod hydrochloride)

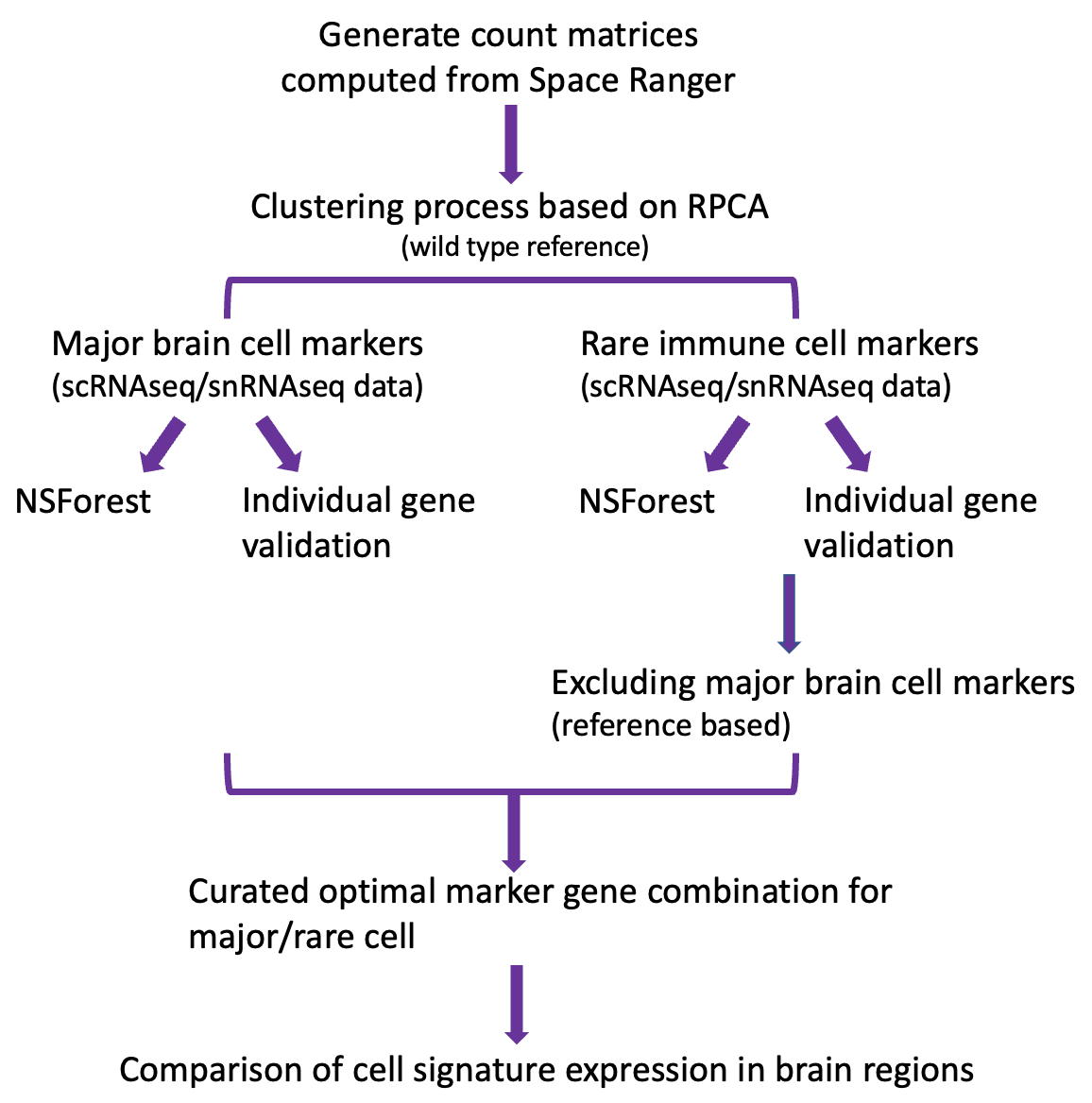

**Supplementary Figure 2. Schematic workflow to identify differences between groups using spatial transcriptome brain imaging data.**

Schematic diagram of workflow carried out in this study. Criteria are specified for each analysis. Gene that do not meet the criteria are excluded. For major brain cell markers, NSForest and individual gene validation methods were used, and for rare immune cell markers, a method of excluding reference-based major brain cell markers was additionally performed. Expression in each brain region was compared using the optimal marker gene combinations selected for major and rare cells. (RPCA: reciprocal principal component analysis; NSForest: Necessary Sufficient Forest; scRNAseq: single cell RNA sequencing; snRNAseq: single nucleus RNA sequencing)

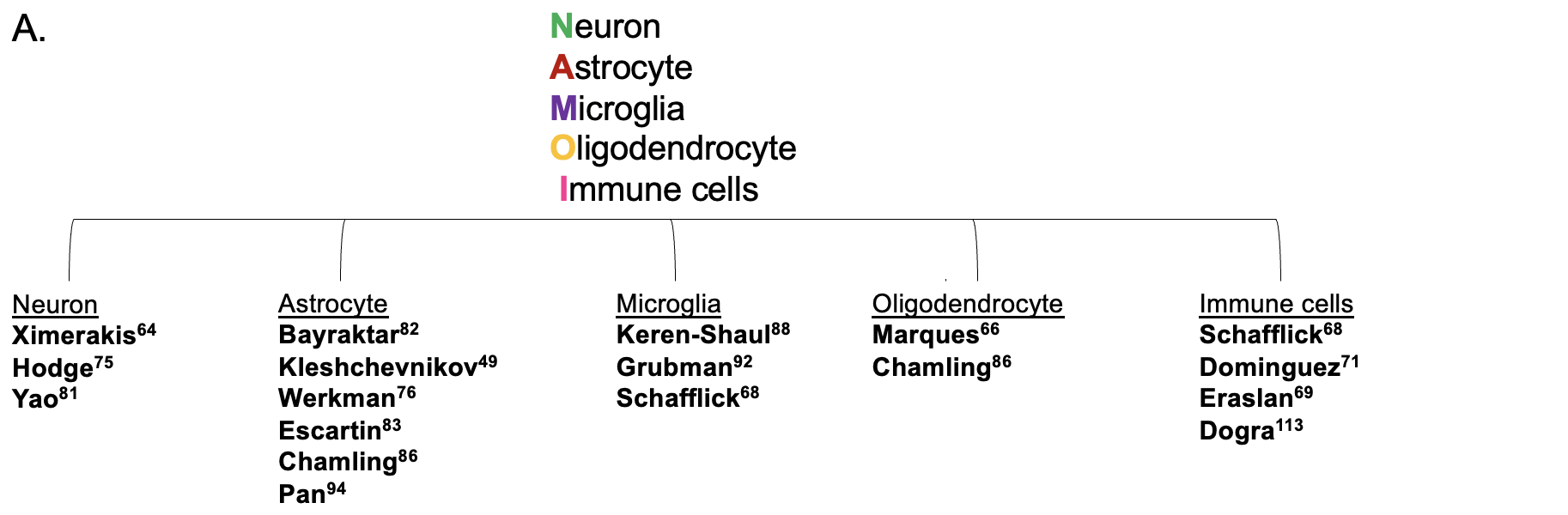

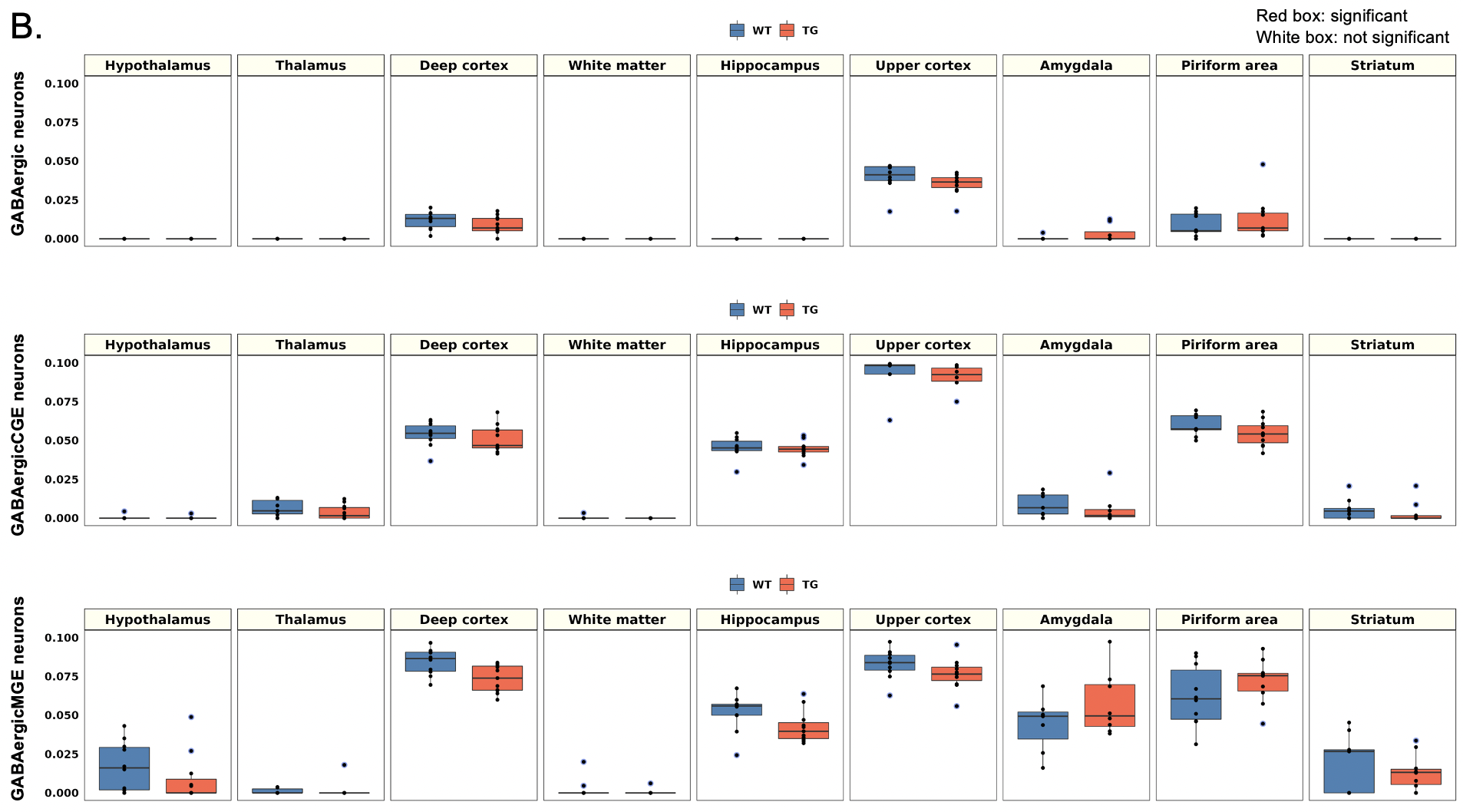

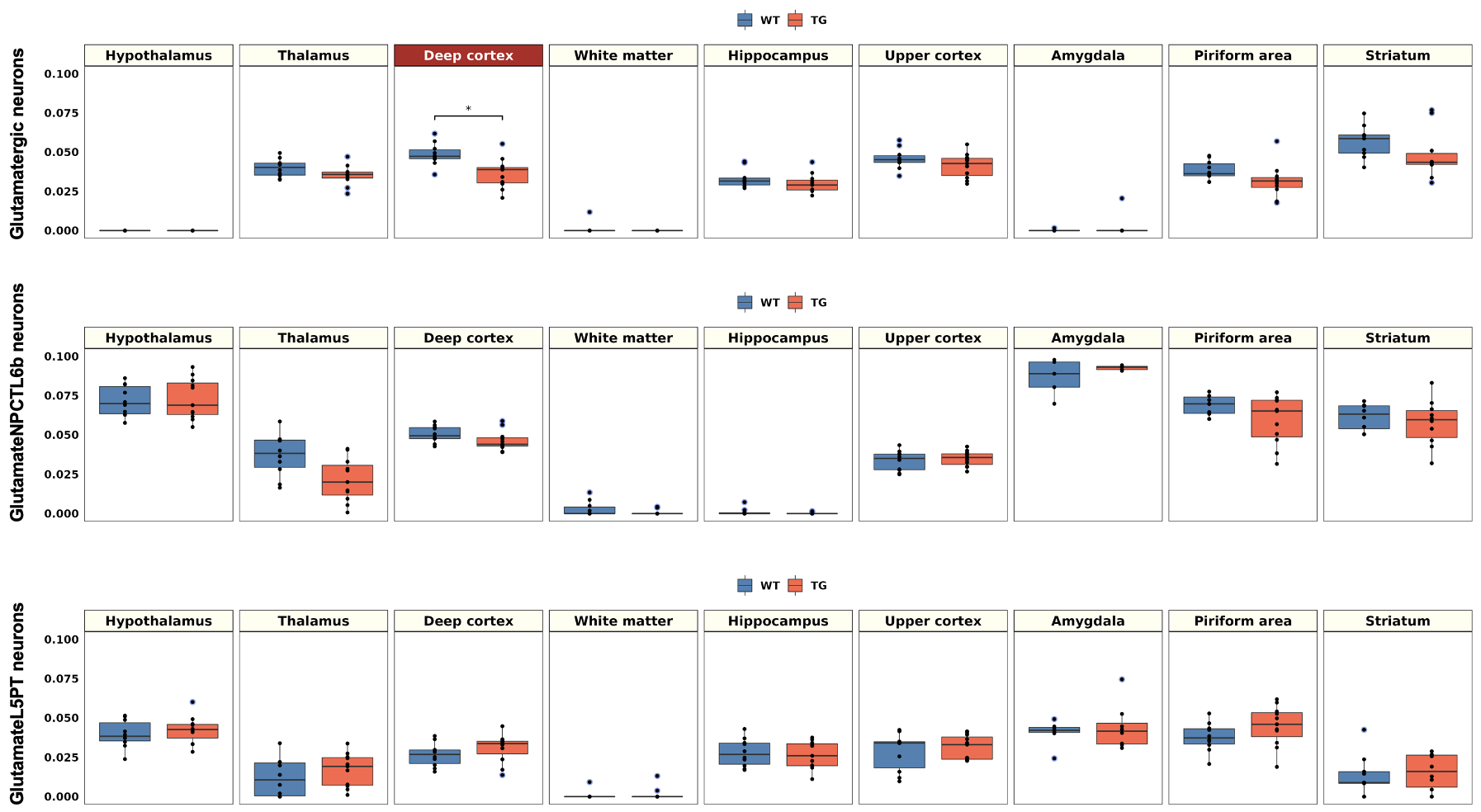

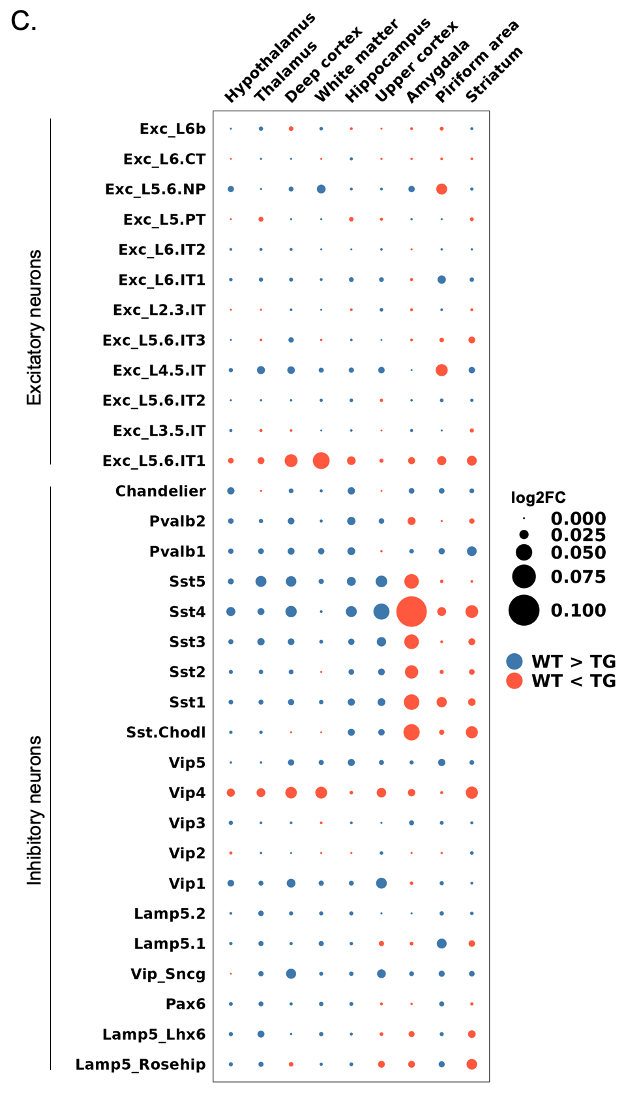

**Supplementary Figure 3.** **Identification of spatial differences in the distribution of neuronal subclasses between wild type and 5xFAD mice.**

(A) References selected for application to neuron, astrocyte, microglia, oligodendrocyte, and immune cell signatures. Detailed information on the references and used gene sets are listed in **Supplementary Table 3**. (B) Boxplot showing average module scores of GABAergic and glutamatergic neuronal signatures. Each dot represents a mouse in each group. The average module scores of the signatures showed no significant differences between wild type and 5xFAD mice, except for a significant decrease in glutamatergic neuronal signatures in the deep cortex of 5xFAD mice. (C) Dot plot showing the average expression differences of the subclasses of excitatory and inhibitory neurons in 9 different regions (hypothalamus, thalamus, deep cortex, white matter, hippocampus, upper cortex, amygdala, piriform area, and striatum). The average expression differences of each brain region between wild type and 5xFAD mice are indicated. For 5xFAD mice compared with wild type mice, the blue dot indicates a decrease, and the orange dot indicates an increase. The size of the dots is proportional to the average expression difference value. Most of the signatures of excitatory and inhibitory neuronal subclasses showed no significant difference between the two groups, but in the Sst subclasses, the expression increase in the amygdala was noticeable in the 5xFAD compare to the wild type mice. Bonferroni-adj. *p-value < 0.05. (Sst: somatostatin; WT: wild type; TG: 5xFAD mice)

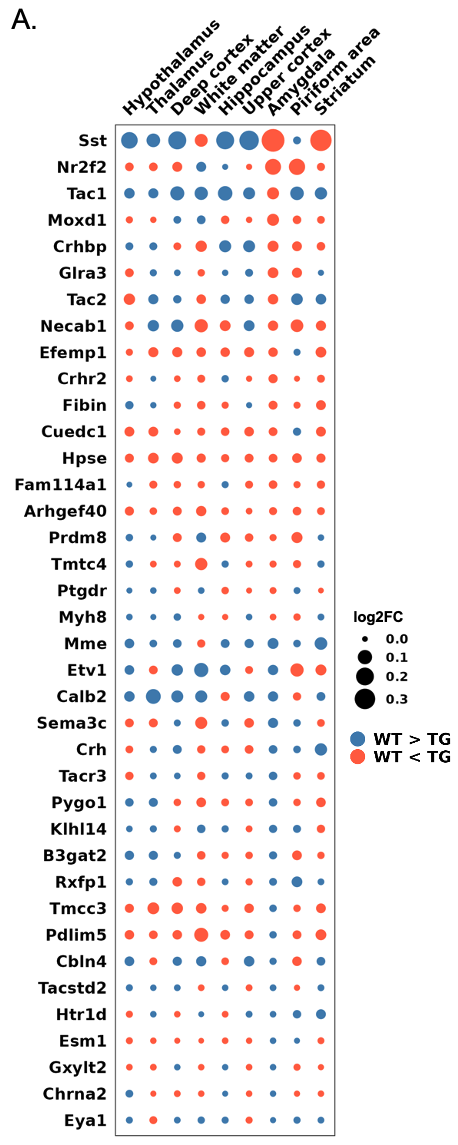

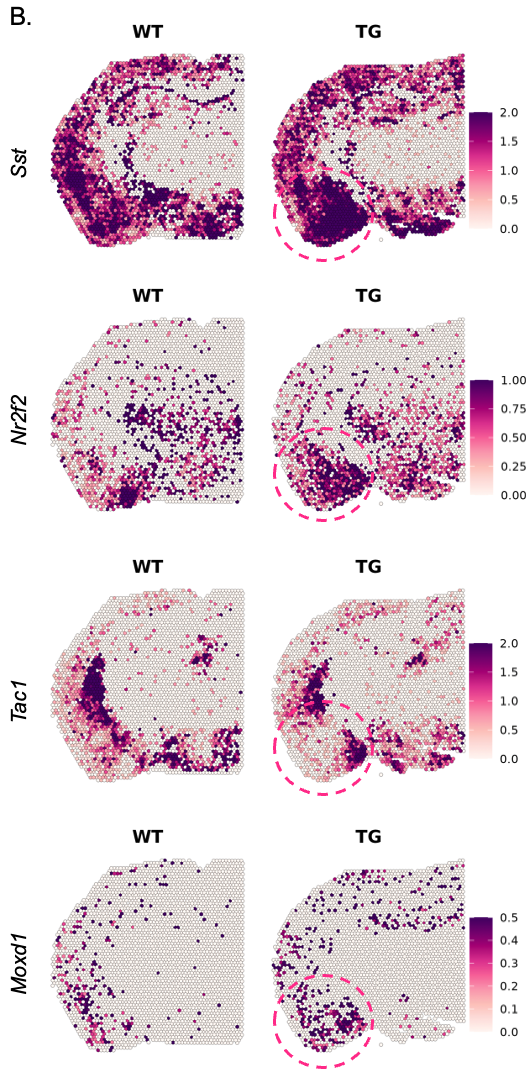

**Supplementary Figure 4. Spatial patterns of individual genes involved in differences in somatostatin inhibitory neuronal signatures in wild type and 5xFAD mice.**

(A) Dot plot showing the average expression differences of individual genes of somatostatin inhibitory neuronal signatures in 9 different regions (hypothalamus, thalamus, deep cortex, white matter, hippocampus, upper cortex, amygdala, piriform area, and striatum). The average expression differences of each brain region between wild type and 5xFAD mice are indicated. For 5xFAD mice compared with wild type mice, the blue dot indicates a decrease, and the orange dot indicates an increase. The size of the dots is proportional to the average expression difference value. (B) Spatial pattern of the selected individual genes showing at least 0.05 of the average expression difference values (*Sst, Nr2f2, Tac1,* and *Moxd1*). The genes showed a tendency to increase expression in the amygdala of 5xFAD compared to wild type mice. (Sst: somatostatin; WT: wild type; TG: 5xFAD mice)

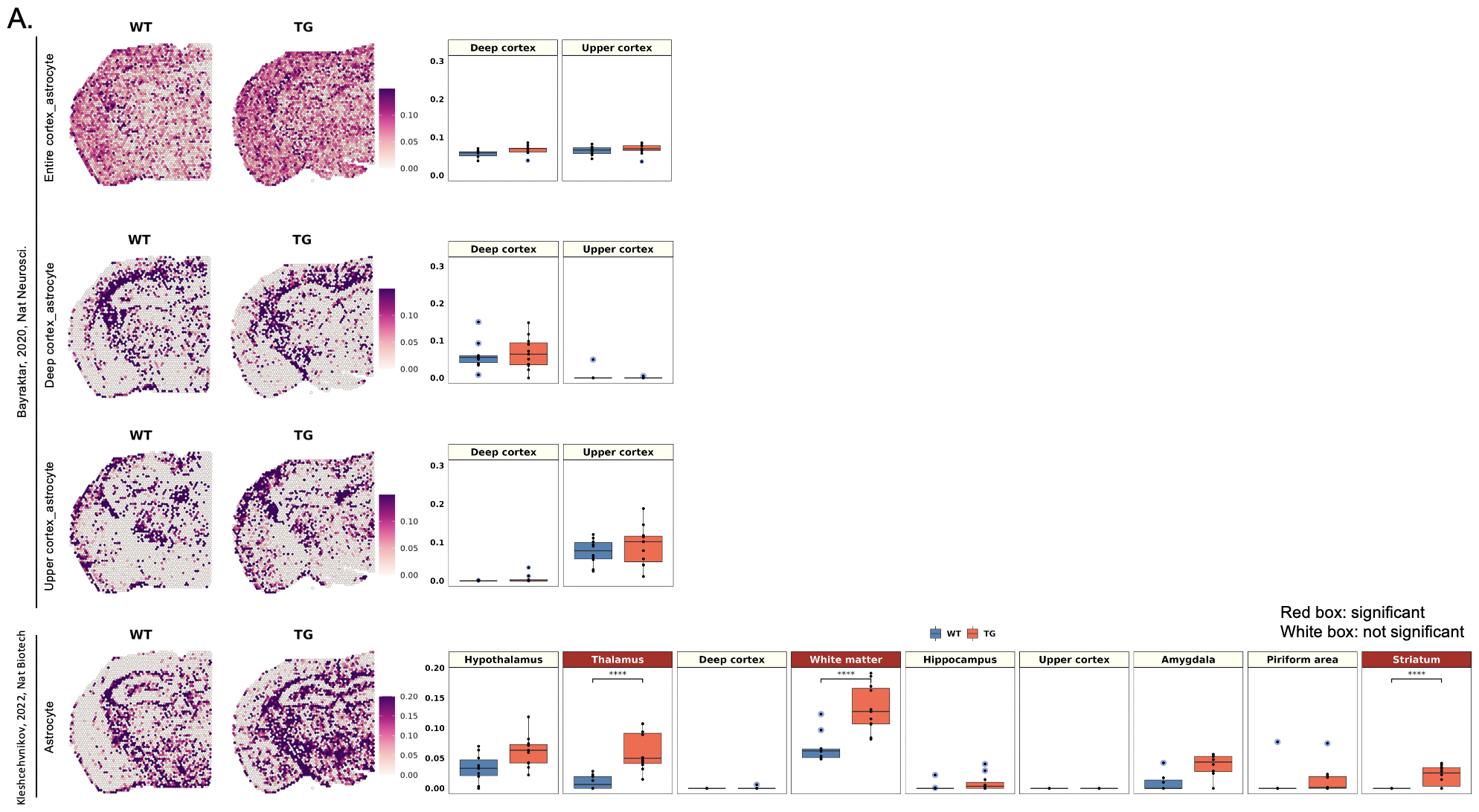

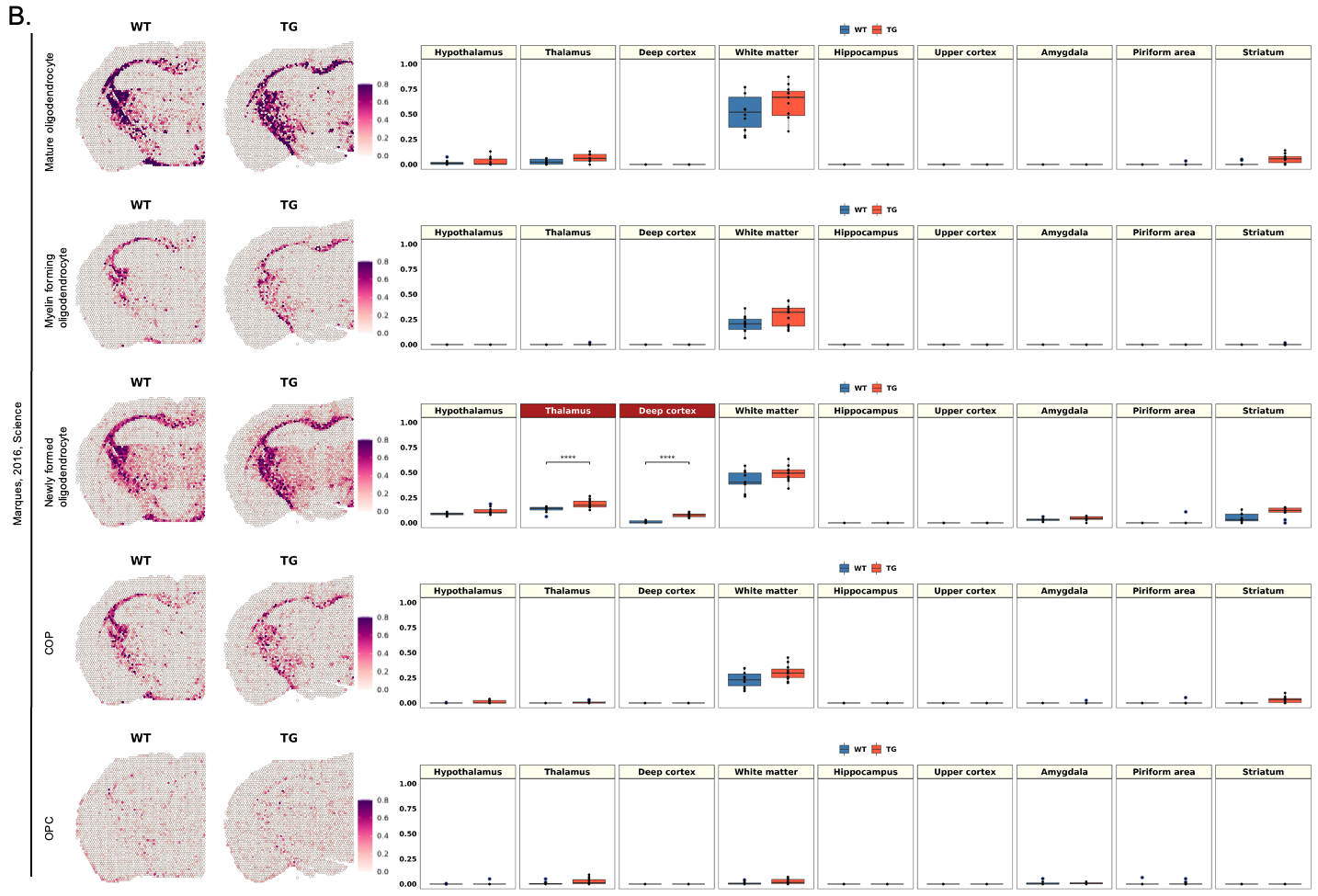

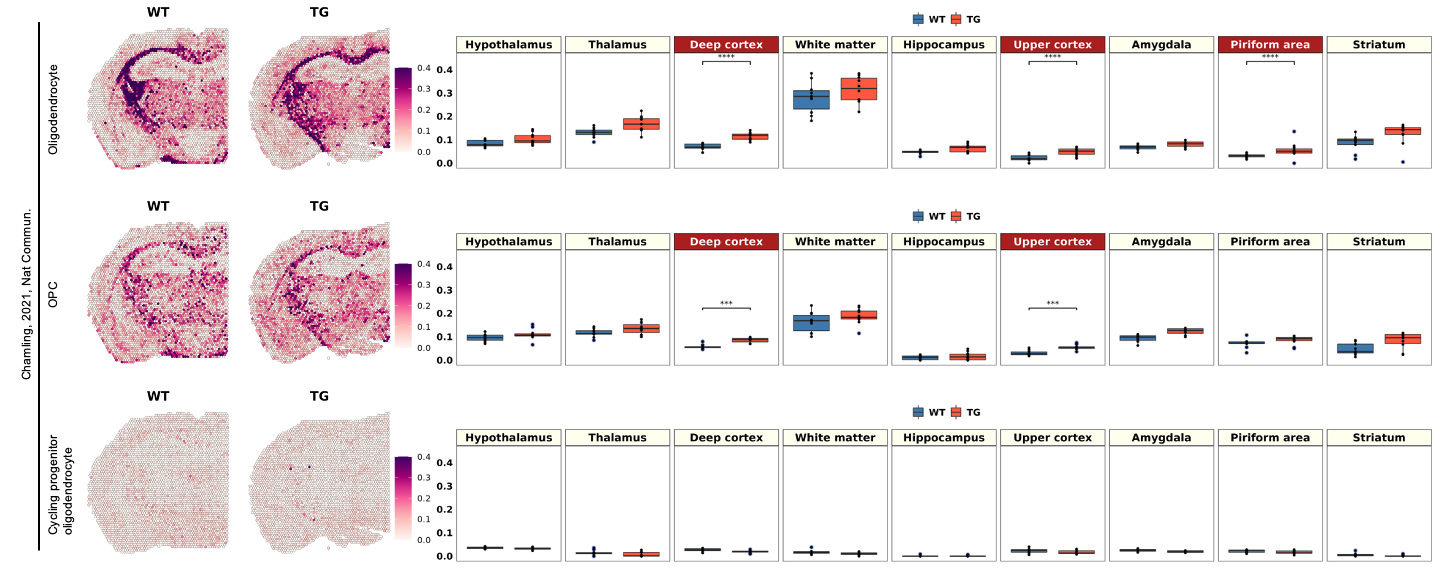

**Supplementary Figure 5.** **Spatial differences in the distribution of the region- or subtypes-specific signatures of glial cells between wild type and 5xFAD mice.**

(A) Spatial pattern of the region-specific signatures (entire cortex, deep cortex, and upper cortex astrocytes) and pan-astrocyte signatures (left). Representative images of each group were selected among 10 spatial transcriptome data from wild type mice and 11 from 5xFAD mice. Boxplot showing average module scores (right). Each dot represents a mouse in each group. The average module score of cortex region-associated astrocyte showed no significant differences between 5xFAD and wild type mice. In the expression of the pan-astrocyte signature, a significant increase in the white matter of 5xFAD mice was most noticeable, and significant differences were also observed in the thalamus and striatum regions. (B) Spatial pattern of different subtypes of oligodendrocyte signatures (mature, myelin forming, newly formed oligodendrocyte, COP, and OPC according to Marques et al.^64^; Oligodendrocyte, OPC, cycling progenitor oligodendrocyte according to Chamling et al.^84^). The average module score of diverse subtypes of oligodendrocyte showed no remarkable differences for any region in the 5xFAD compared to wild type mice. A significant but slight difference in deep cortex and thalamus for newly formed oligodendrocytes, and deep and upper cortex for oligodendrocyte and OPC by Chamling et al.^84^ piriform area for oligodendrocyte was small in magnitude. In summary, microglia and astrocyte showed significantly difference between 5xFAD compared to wild type mice, but oligodendrocyte relatively little changes. Bonferroni-adj. *p-value < 0.05, ***p-value < 0.001, ****p-value < 0.0001. (WT: wild type; TG: 5xFAD mice; OPC: oligodendrocyte precursor cells; COP: committed oligodendrocyte progenitors)

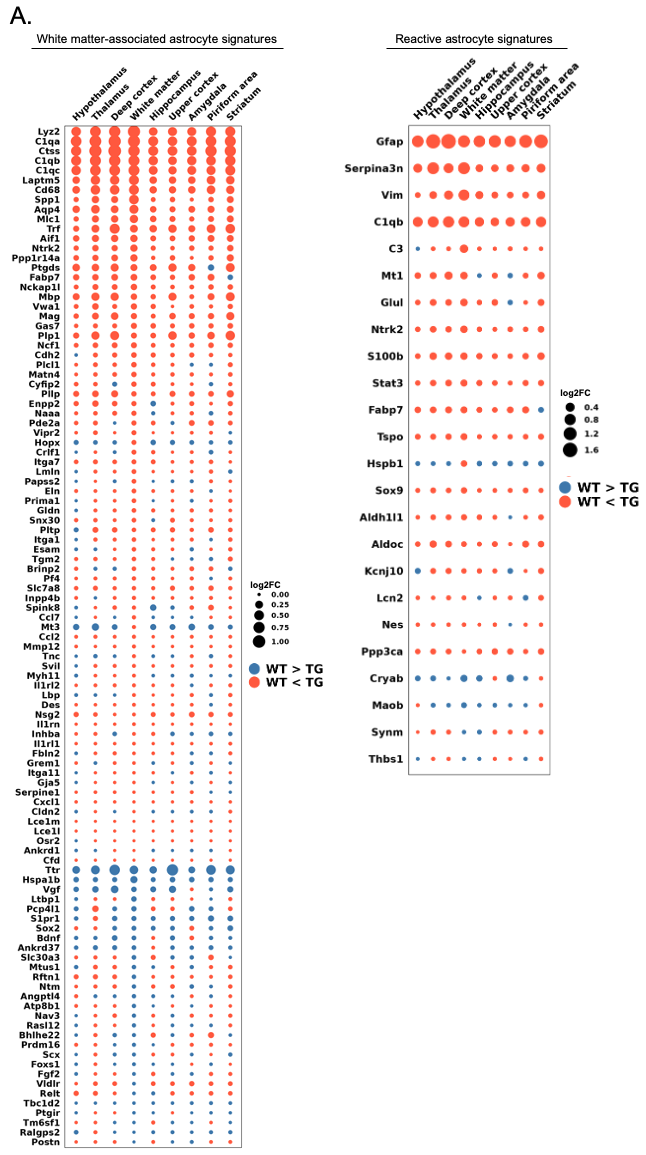

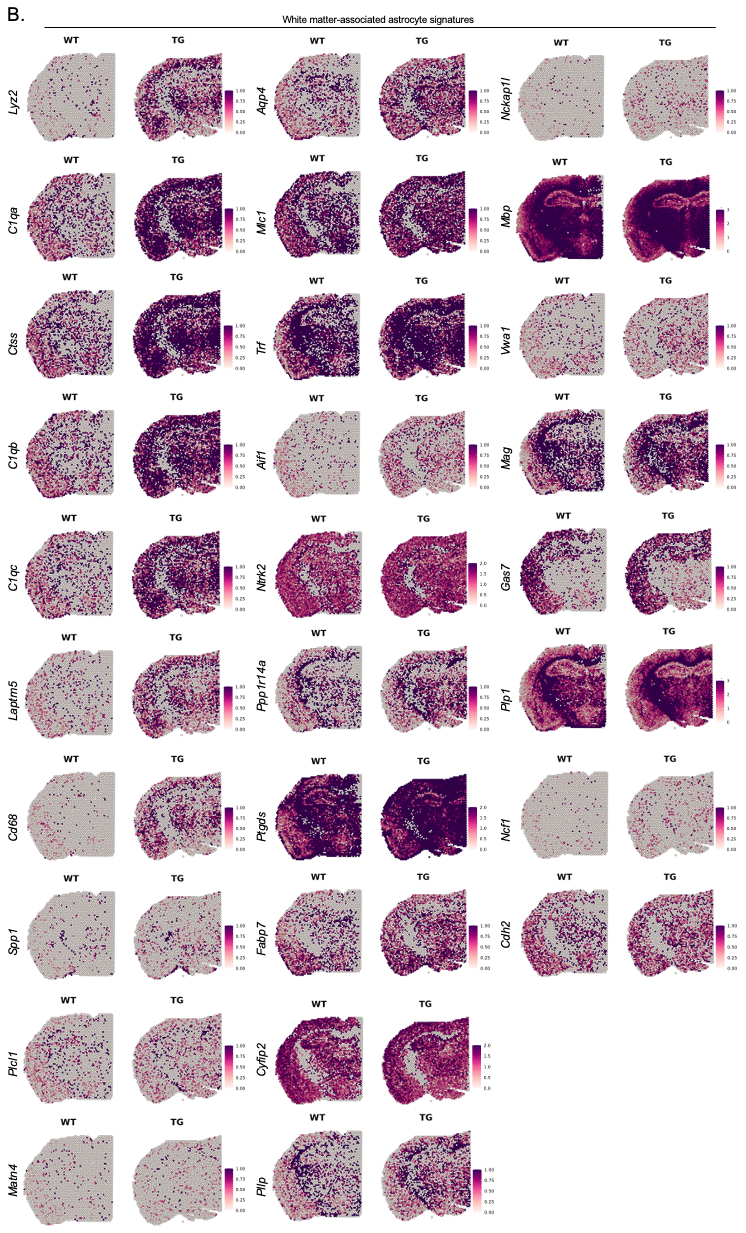

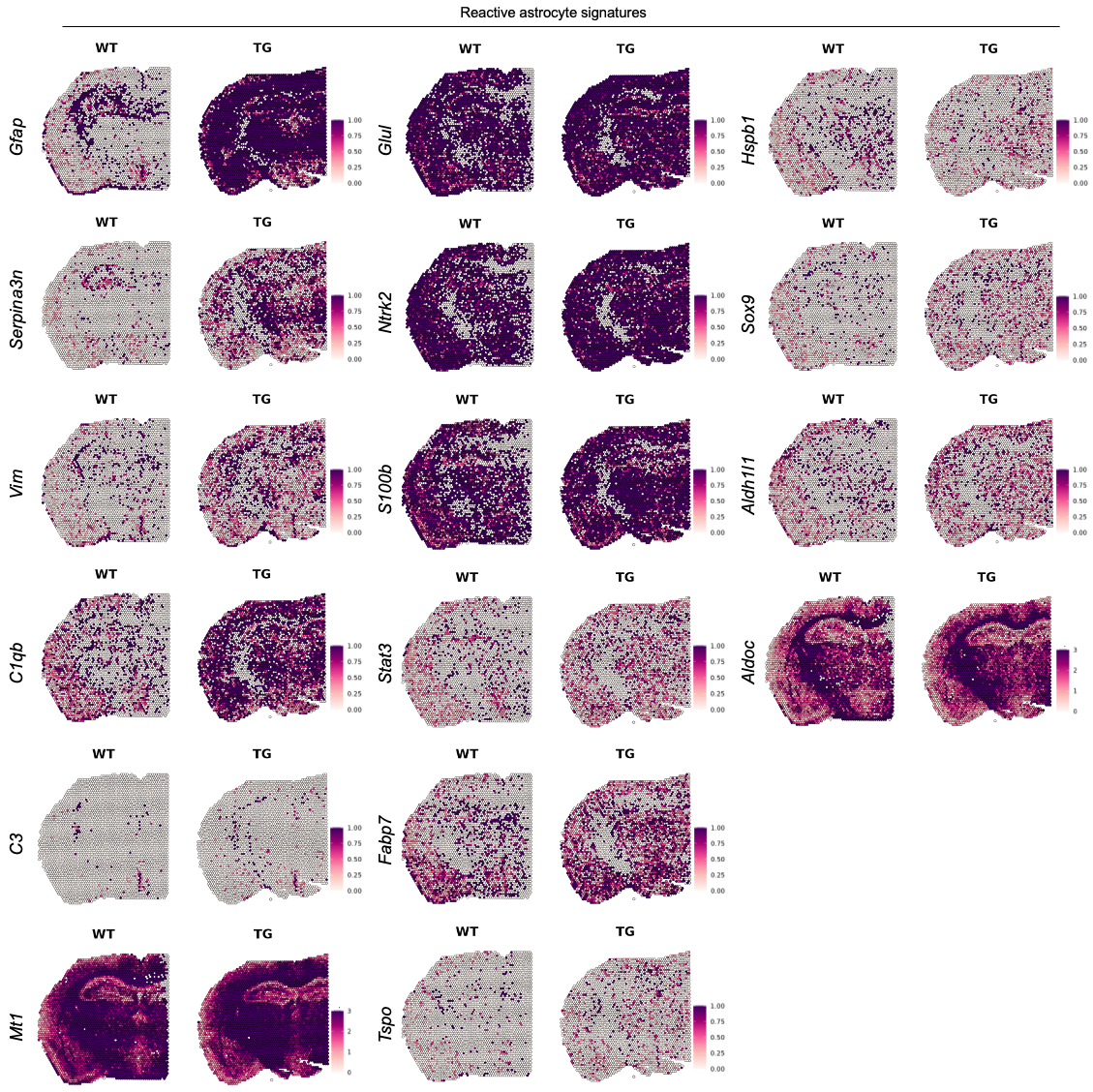

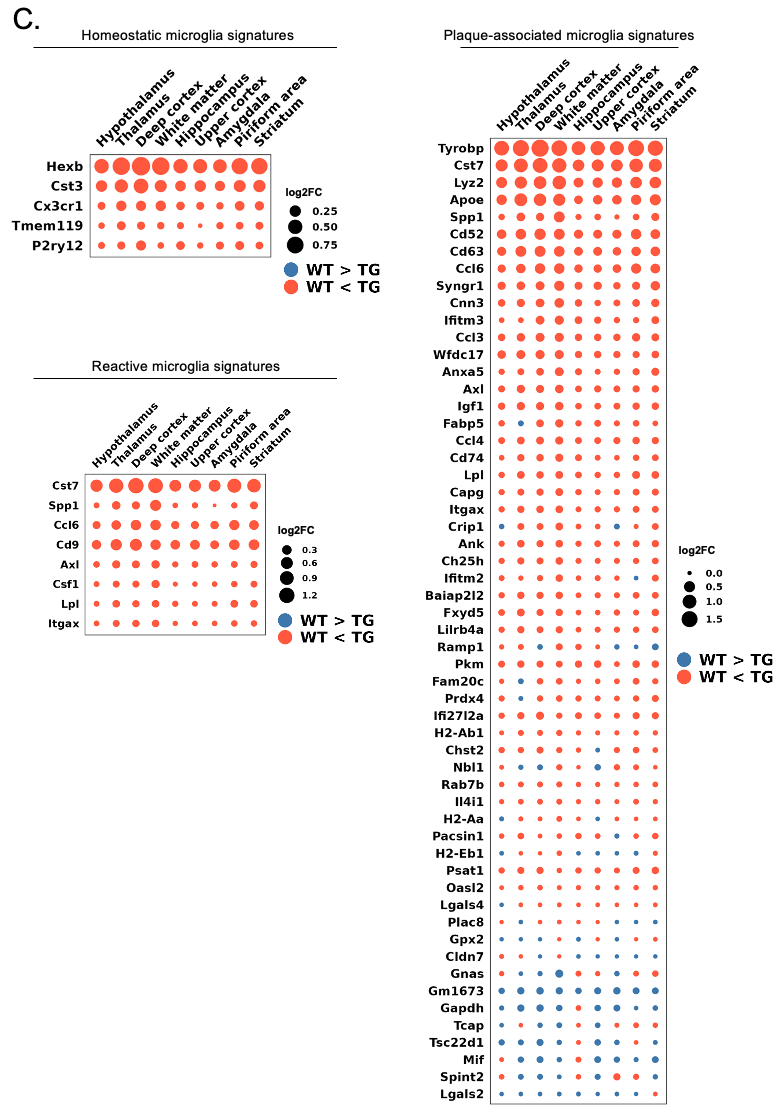

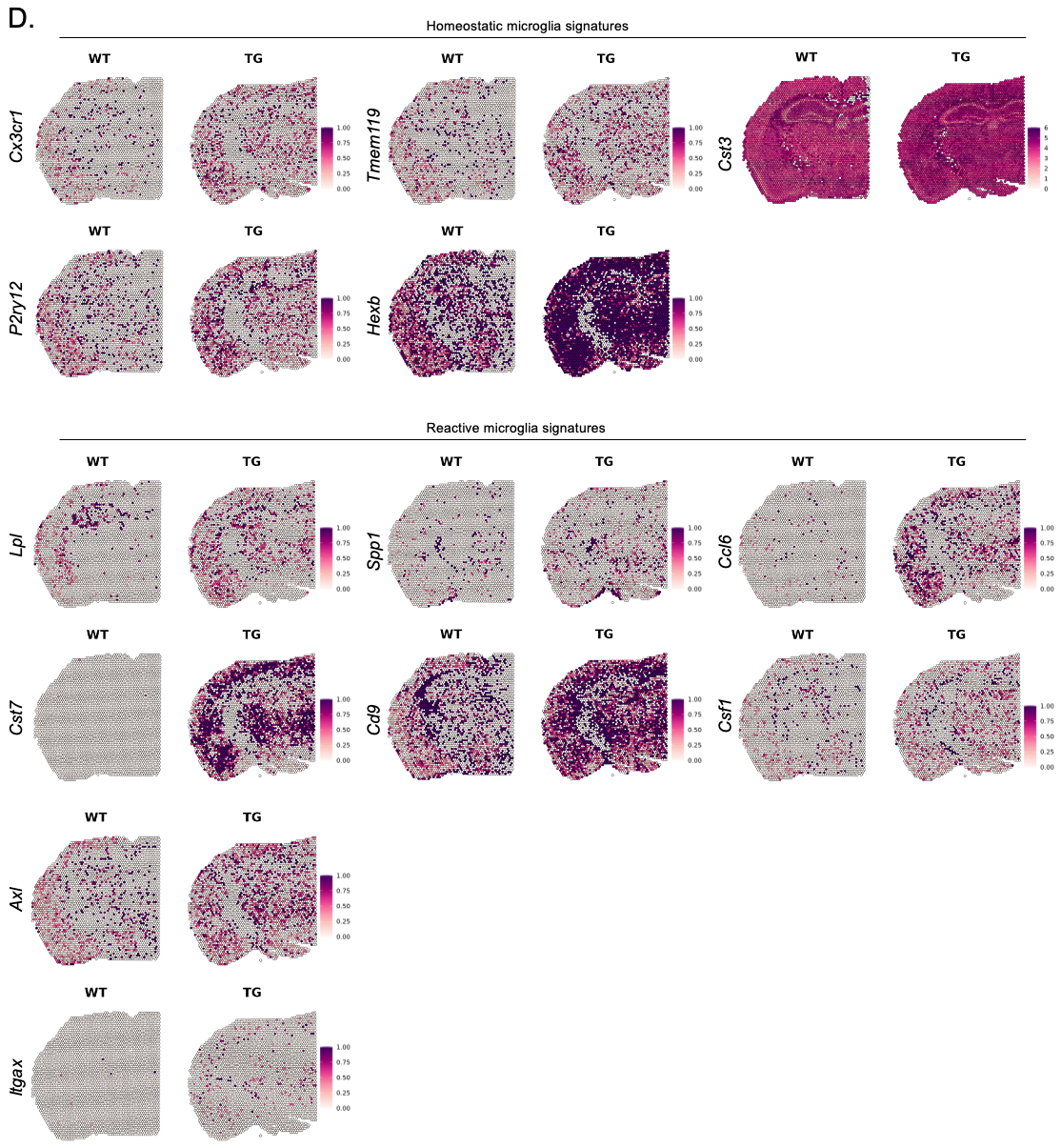

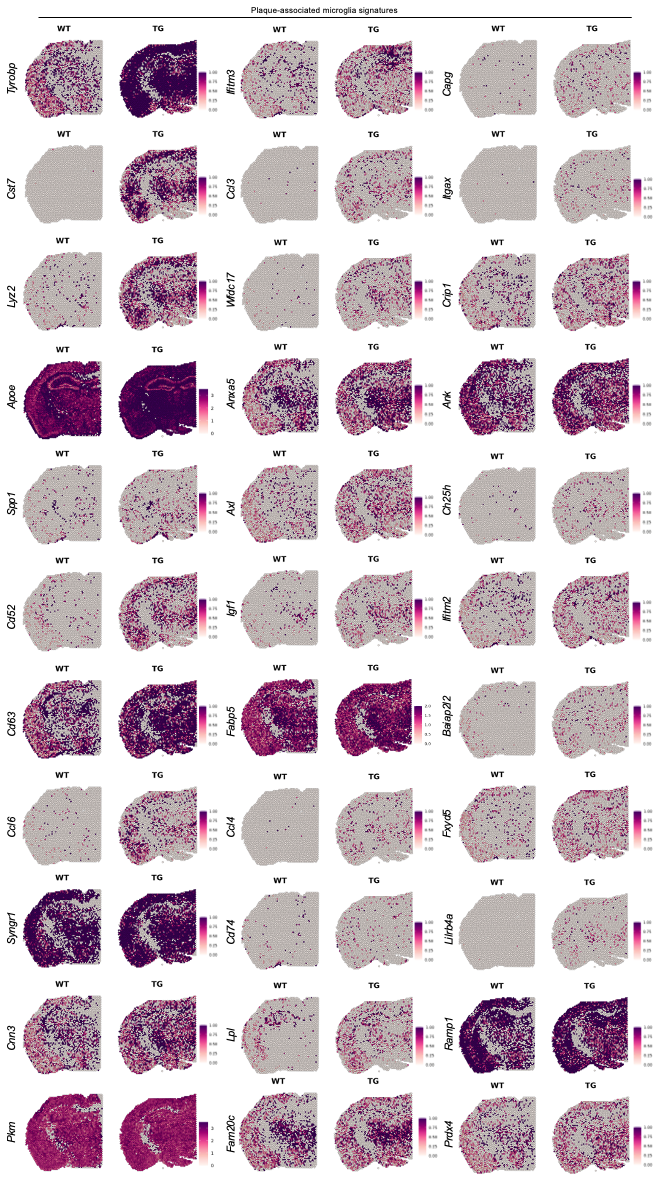

**Supplementary Figure 6. Spatial patterns of individual genes involved in signature differences of microglia and astrocytes between wild type and 5xFAD mice.**

(A) Dot plot showing the average expression differences of individual genes of region- or state-specific astrocyte signatures in 9 different regions (hypothalamus, thalamus, deep cortex, white matter, hippocampus, upper cortex, amygdala, piriform area, and striatum). The average expression differences of each brain region between wild type and 5xFAD mice are indicated. For 5xFAD mice compared with wild type mice, the blue dot indicates a decrease, and the orange dot indicates an increase. The size of the dots is proportional to the average expression difference value. (B) Spatial pattern of the selected individual genes showing at least 0.05 of the average expression difference values. (C) Dot plot showing the average expression differences of individual genes of state-specific microglial signatures. The average expression differences of each brain region between wild type and 5xFAD mice are indicated. (D) Spatial pattern of the selected individual genes showing at least 0.05 of the average expression difference values. The astrocyte and microglia-related genes tended to increase expression across all major brain regions in 5xFAD compared to wild type mice. (WT: wild type; TG: 5xFAD mice)

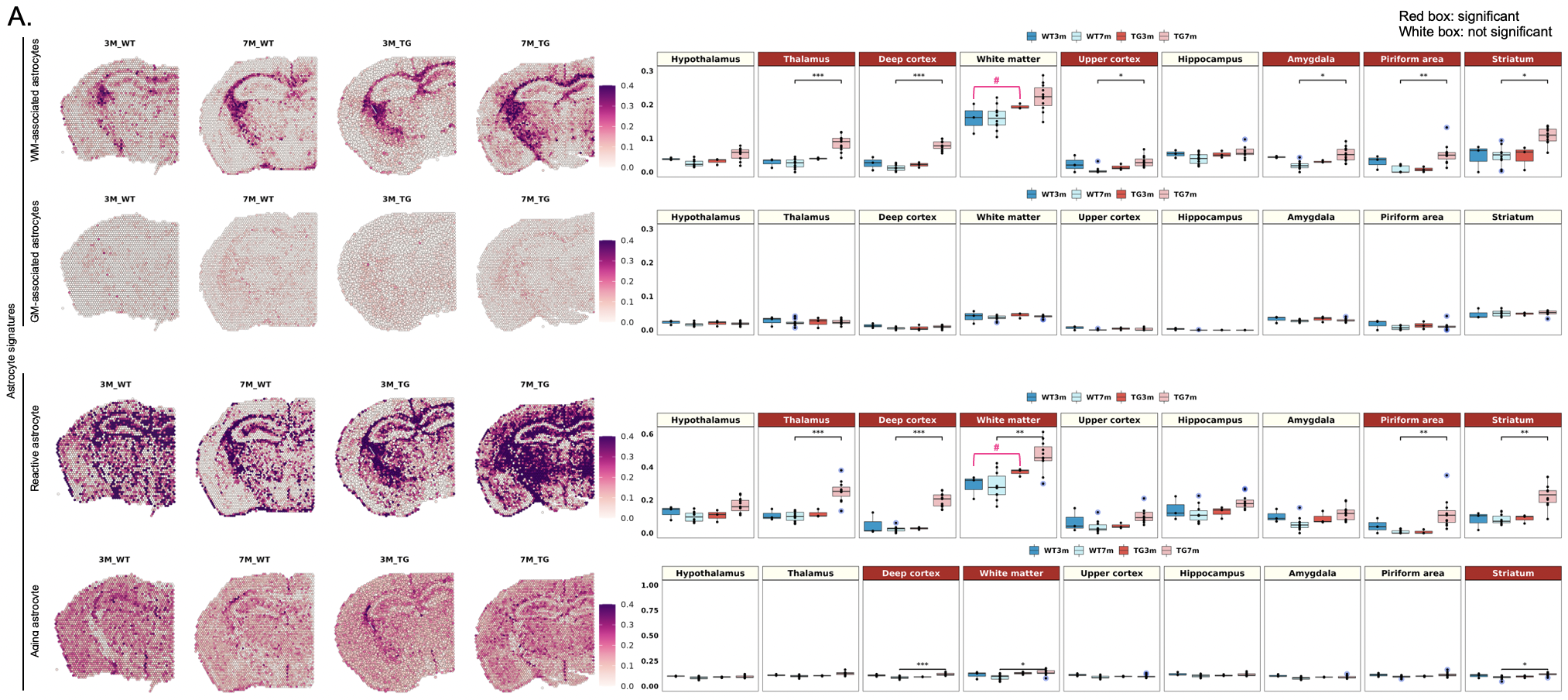

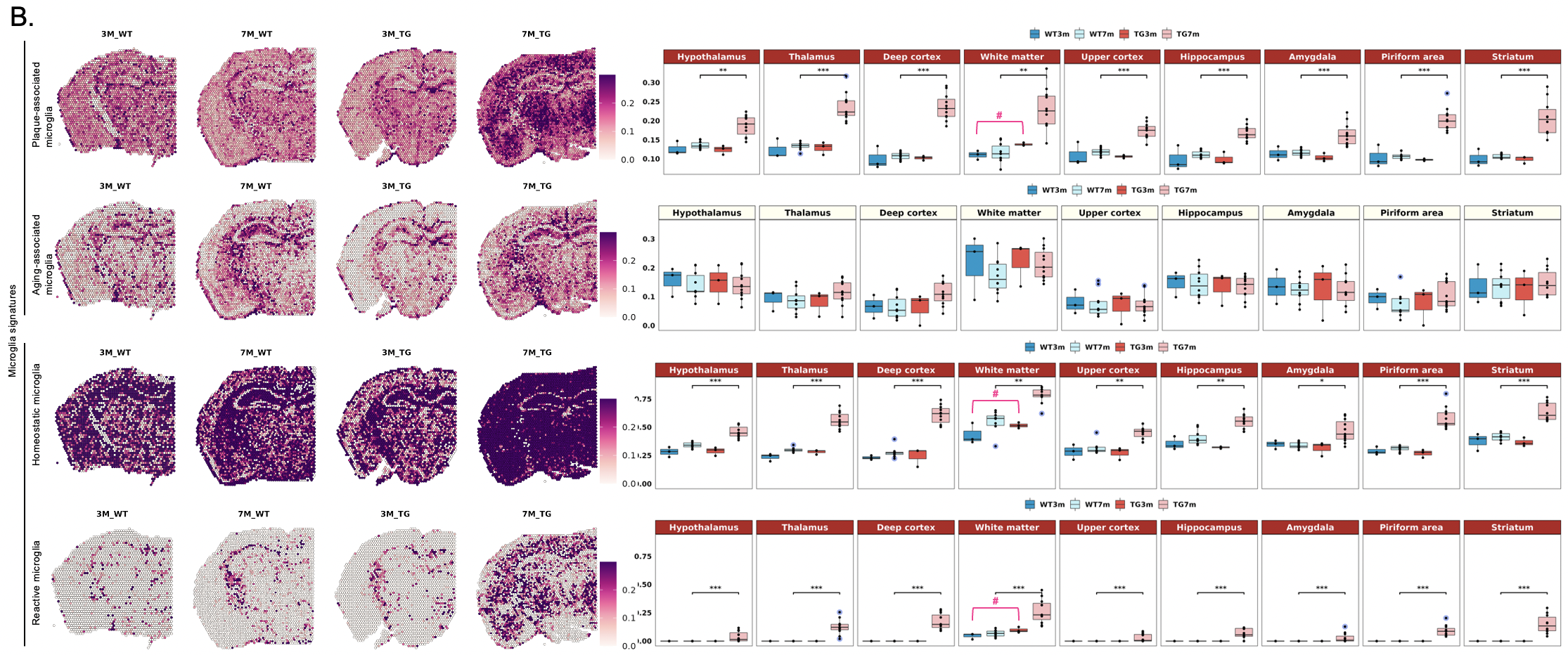

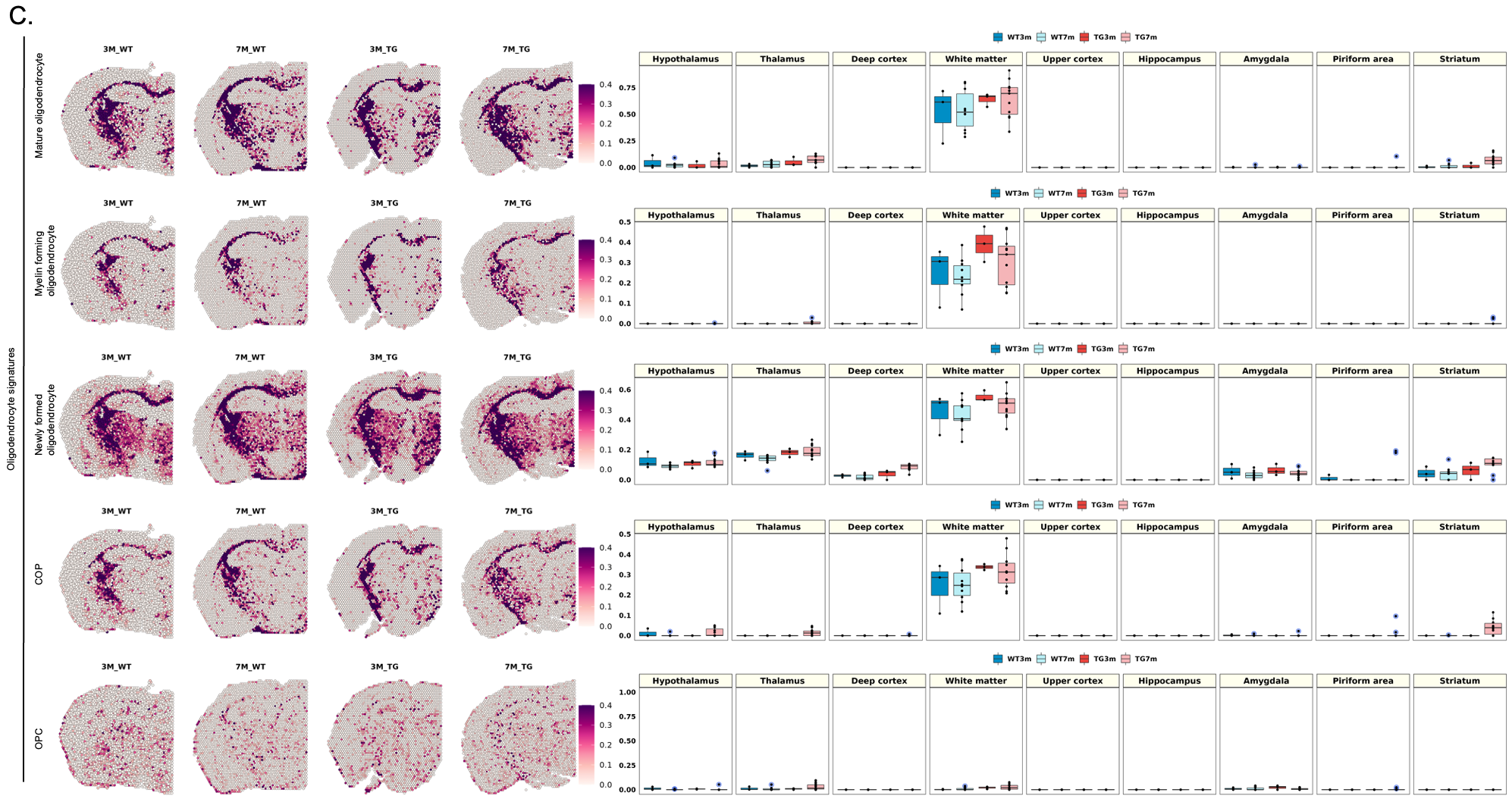

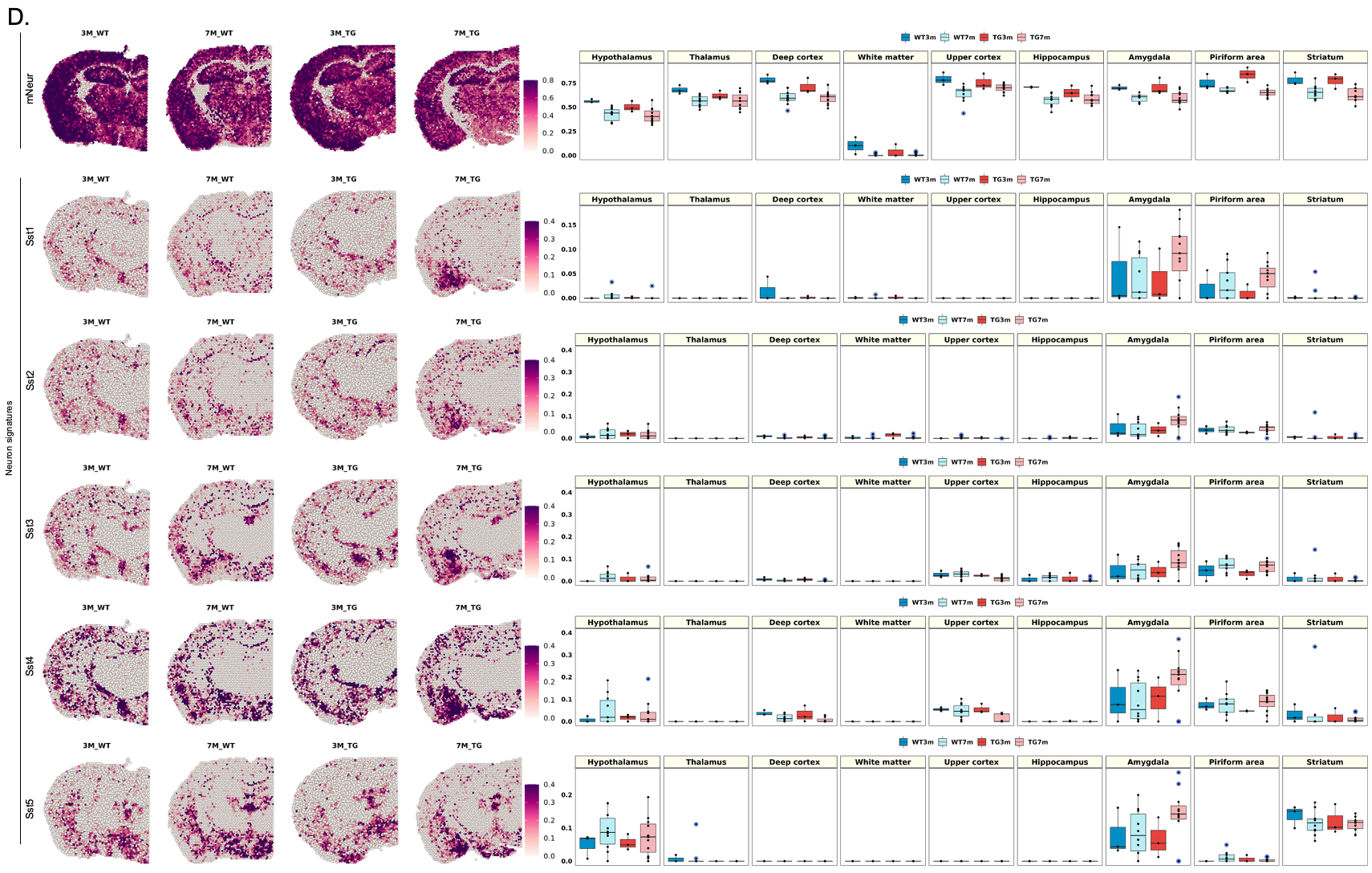

**Supplementary Figure 7. Spatial changes in the distribution of brain cell signatures in 3-month-old compared to 7-month-old 5xFAD mice.**

(A) Spatial pattern of the region-specific signatures (white matter-associated and gray matter-associated astrocytes) and the state-specific signatures (reactive astrocyte and aging astrocyte; left), (B) the state-specific signatures (plaque-associated, aging-associated, homeostatic, reactive, and pan-microglia), (C) the subtypes of oligodendrocyte signatures (-mature, myelin forming, newly formed oligodendrocyte, COP, and OPC according to Marques et al.^64^), and (D) the neuronal signatures (mature neurons, Sst1, Sst2, Sst3, Sst4, and Sst5; left). Boxplot showing average module scores (right). Each dot represents a mouse in each group. The average module score of white matter-associated astrocytes, reactive astrocytes, plaque-associated microglia, homeostatic, and reactive microglia tended to increase exclusively in the white matter in the 3-month-old 5xFAD compared to wild type mice. It is suggested that the earlier (3-month old) changes of microglia and astrocytes preceded in the white matter later (7-month old) progress in 5xFAD mice showing behavior abnormalities (and amyloid plaques). Other brain cell signatures did not show any differences between 3-month and 7-month of age in 5xFAD mice. Groups showing differences between 3-month-old wild type and 3-month-old 5xFAD mice, not significant but showing tendency, are marked with # in pink. Bonferroni-adj. *p-value < 0.05, **p-value < 0.01, ****p-value < 0.0001. (WM: white matter; GM: gray matter; WT: wild type; TG: 5xFAD mice; mNeur: mature neurons; Sst: somatostatin; OPC: oligodendrocyte precursor cells; COP: committed oligodendrocyte progenitors)

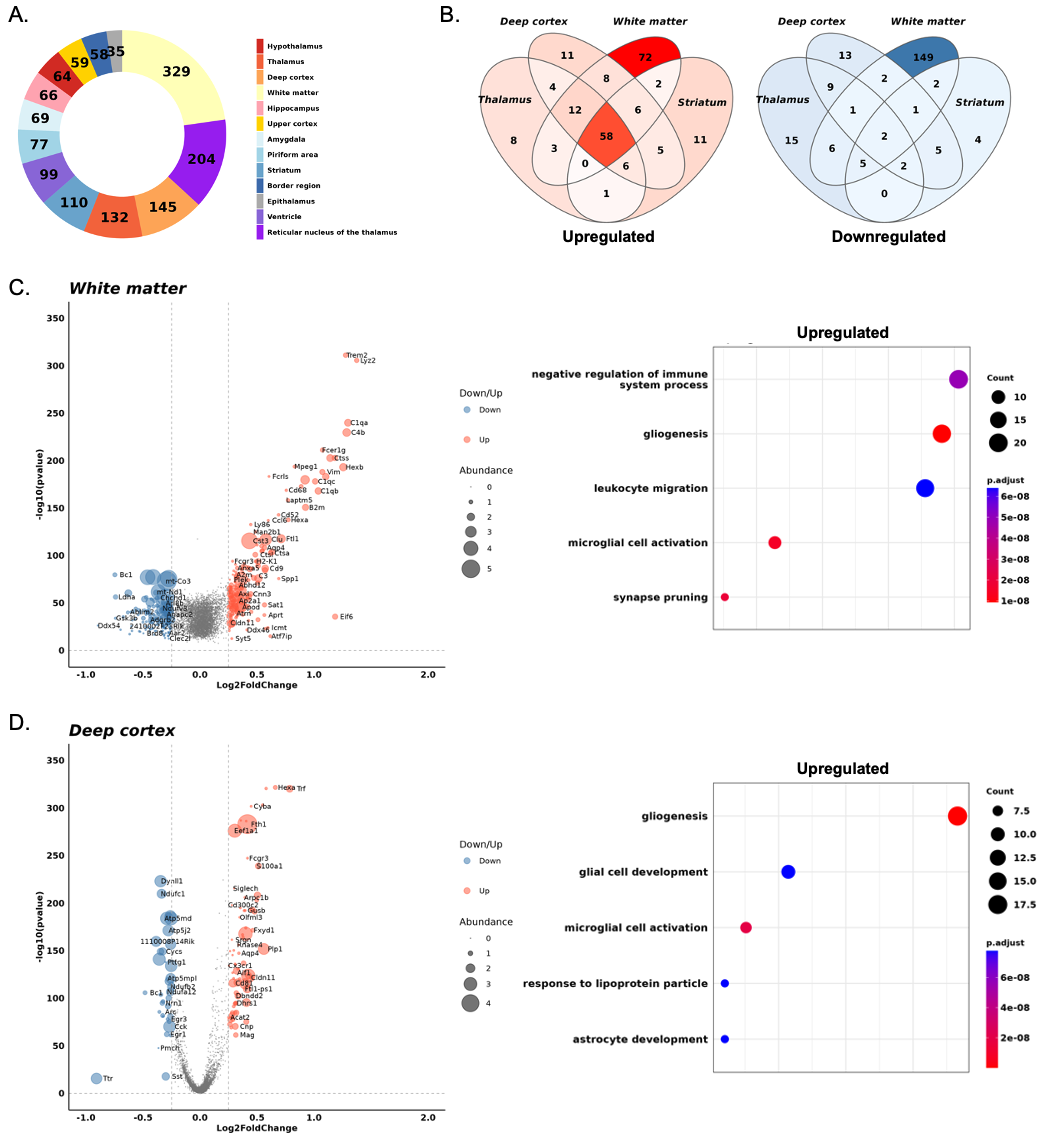

**Supplementary Figure 8. Identification of DEGs and related biological pathways in the 5xFAD compared to wild type mice.**

(A) Pie chart demonstrating the DEG proportion of 5xFAD compared to wild type mice in each brain region. (B) Venn diagram representing the upregulated (left; red) and downregulated (right; blue) DEGs in the 5xFAD compared to wild type mice. The regions with larger number of genes are expressed in vivid colors. (C) Volcano plot for the genes identified in the 5xFAD compared to wild type mice in the white matter and (D) deeper cortex. Genes in the colored dots are significantly (logFC threshold = 0.25) upregulated (red dots) and downregulated (blue dots). The dots colored dark gray represent the genes that are not significantly changed in the 5xFAD mice. The abundance of genes was indicated by the size of the dots using the average expression value of 5xFAD mice. The top five GO terms related to the upregulated (left) and downregulated (right) genes in the white matter and deeper cortex. In 5xFAD mice, both white matter and deep cortex showed the significant increase in gliogenesis and glial cell activation. (DEG: differentially expressed genes; GO: gene ontology)

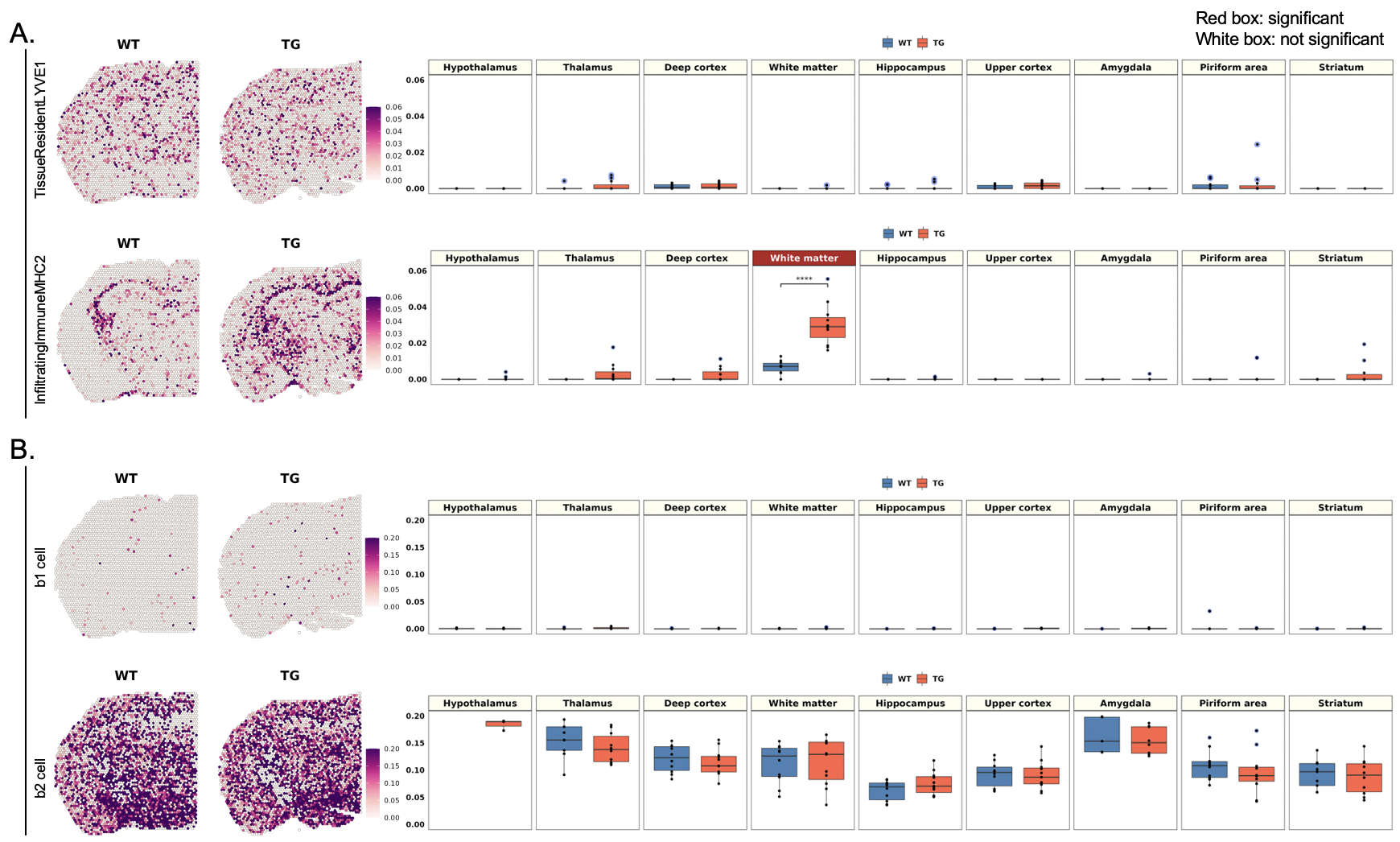

**Supplementary Figure 9. Spatial differences in the distribution of rare brain cell signatures between wild type and 5xFAD mice.**

(A) Spatial pattern of the rare brain cell signatures according to Eraslan et al.^67^ (TissueResidentLYVE1 and InfiltratingImmuneMHC2; left). Representative images of each group were selected among 10 spatial transcriptome data from wild type mice and 11 from 5xFAD mice. Boxplot showing average module scores (right). Each dot represents a mouse in each group. The average module score of LYVE1-expressing tissue resident immune cells showed no significant differences between 5xFAD and wild type mice. MHC2-expressing infiltrating immune cell signature increased significantly in the white matter of 5xFAD mice. (B) Spatial pattern of B cell signatures (b1 cell and b2 cell; left) and boxplot showing average module scores (right). The expression showed no significant differences between 5xFAD and wild type mice. (WT: wild type; TG: 5xFAD mice; LYVE1: lymphatic vessel endothelial hyaluronan receptor 1; MHC2: major histocompatibility complex 2)

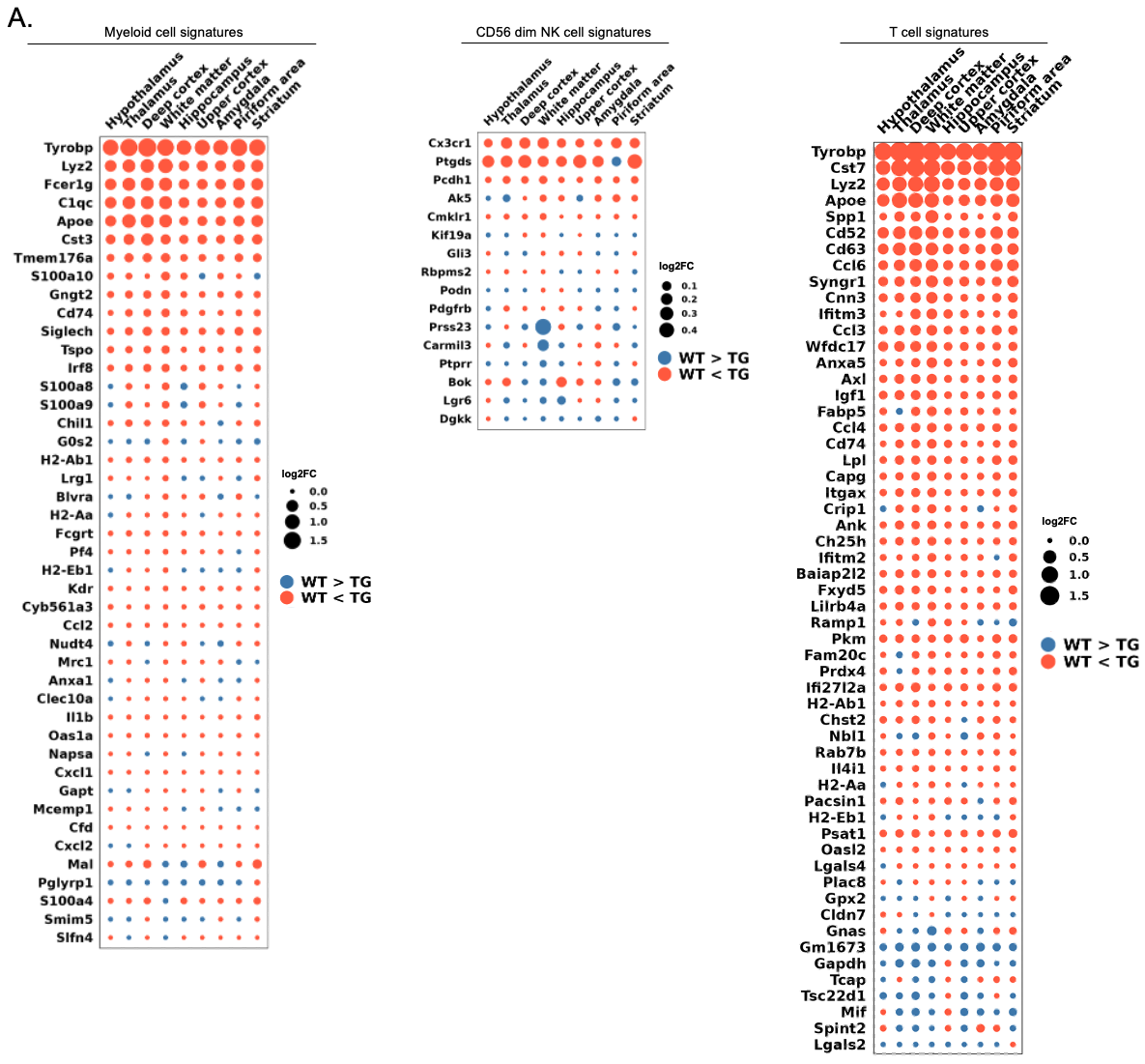

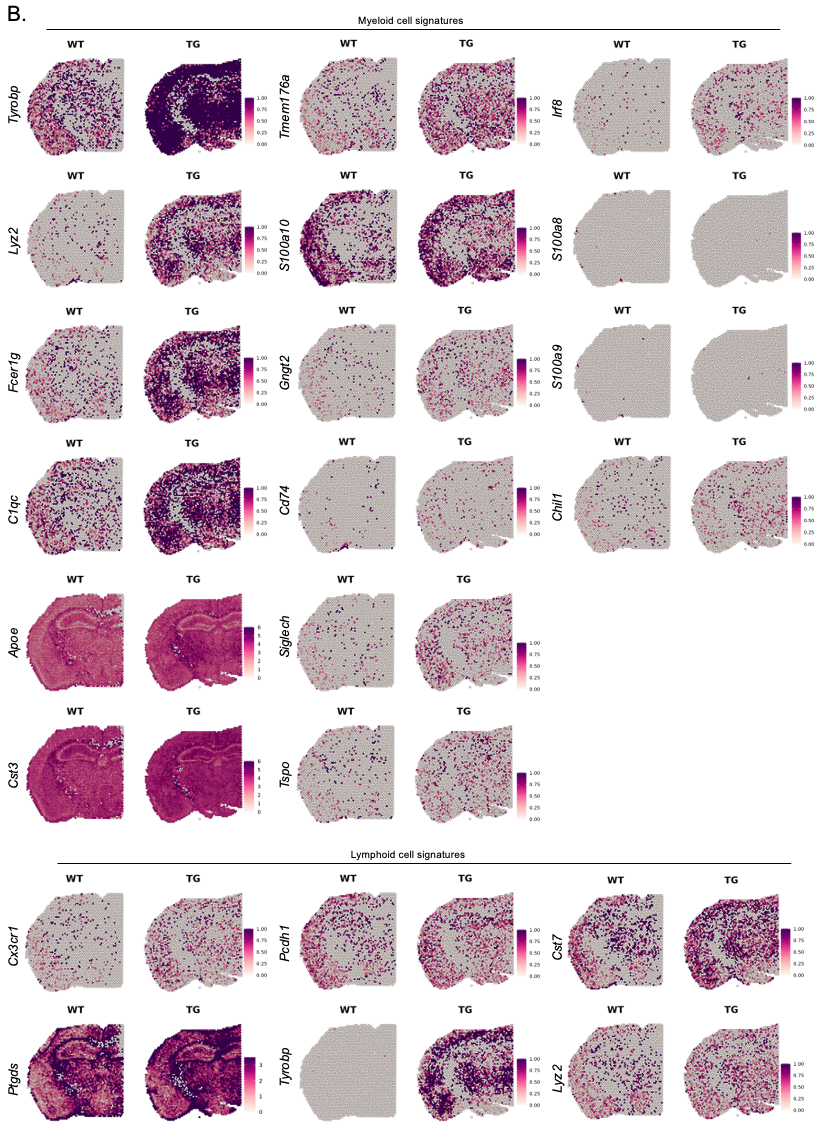

**Supplementary Figure 10.** **Spatial patterns of individual genes involved in differences in type/subtype signatures of rare immune cells between wild type and 5xFAD mice.**

(A) Dot plot showing the average expression differences of individual genes of rare immune cell signatures (myeloid, CD56 dim NK, and T cells) in 9 different regions (hypothalamus, thalamus, deep cortex, white matter, hippocampus, upper cortex, amygdala, piriform area, and striatum). The average expression differences of each brain region between wild type and 5xFAD mice are indicated. For 5xFAD mice compared with wild type mice, the blue dot indicates a decrease, and the orange dot indicates an increase. The size of the dots is proportional to the average expression difference value. (B) Spatial pattern of the selected individual genes showing at least 0.05 of the average expression difference values. The rare immune cell-related genes tended to increase expression across all major brain regions in 5xFAD compared to wild type mice. (WT: wild type; TG: 5xFAD mice)

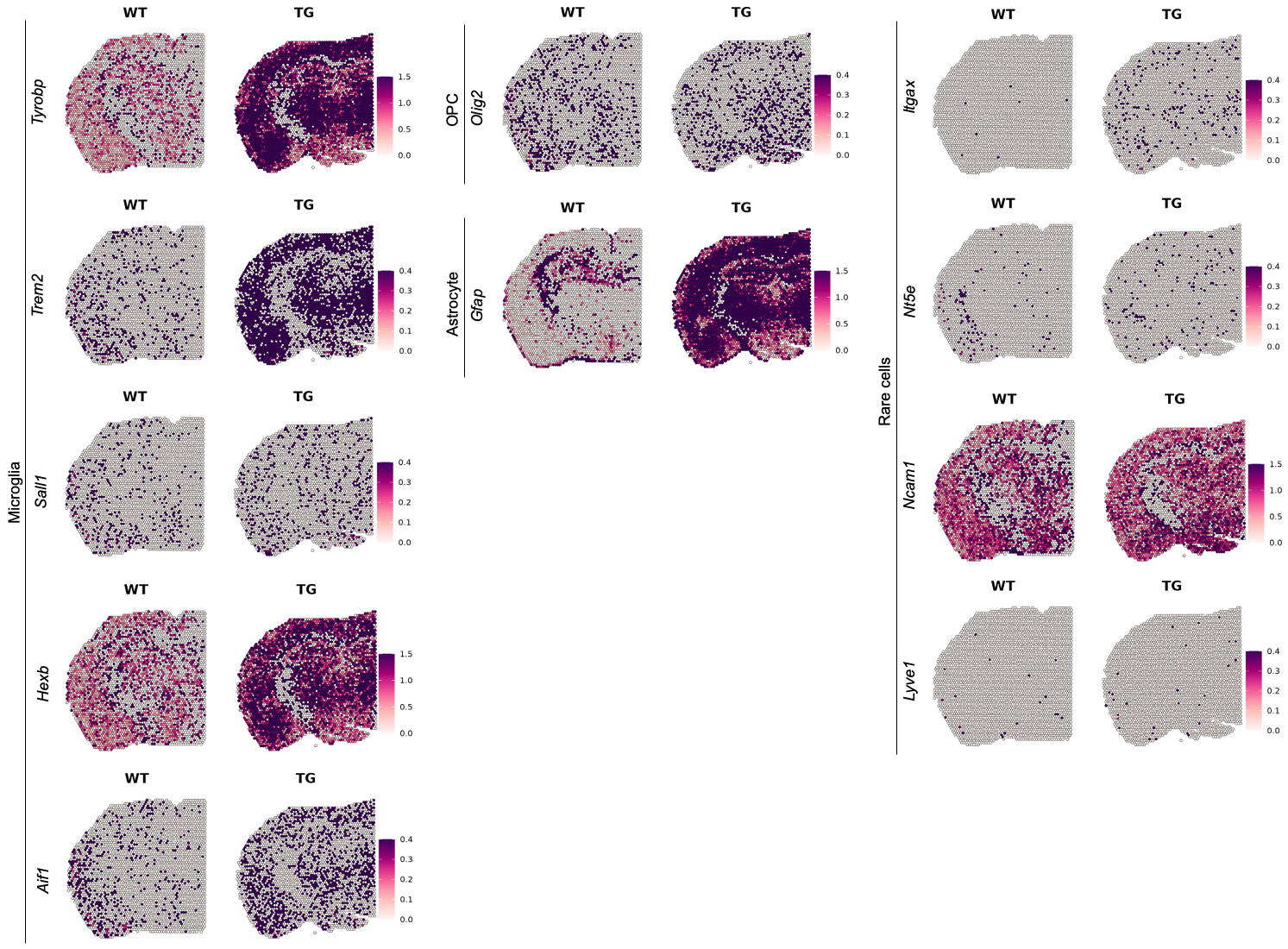

**Supplementary Figure 11.** **Spatial differences of individual genes specifying brain major or rare cell types between wild type and 5xFAD mice.**

Spatial pattern of single gene expression as markers for brain major cells (microglia: *Tyrobp, Trem2, Sall1, Hexb,* and *Aif1*; OPC: *Olig2*; astrocyte: *Gfap*) and rare cells (*Itgax, Nt5e, Ncam1,* and *Lyve1*). Representative images of each group were selected among 10 spatial transcriptome data from wild type mice and 11 from 5xFAD mice. (WT: wild type; TG: 5xFAD mice; OPC: oligodendrocyte precursor cells)

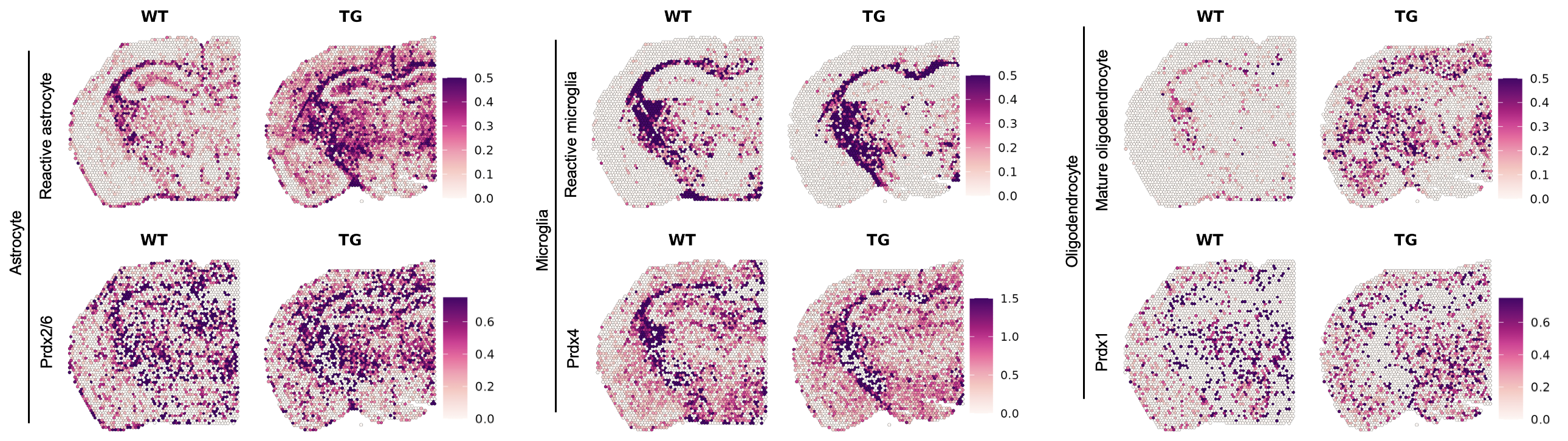

**Supplementary Figure 12.** **Spatial distribution patterns of specific isoforms of peroxiredoxin representing different major brain cells.**

Spatial pattern of the reactive astrocyte signature and *Prdx2/6*, the reactive microglia signature and *Prdx4*, and the mature oligodendrocyte signature and *Prdx1*. Representative images of each group were selected among 10 spatial transcriptome data from wild type mice and 11 from 5xFAD mice. The isoforms of each peroxiredoxin showed different spatial patterns of distribution between wild type and 5xFAD mice. *Prdx6* and *Prdx2* for astrocytes, *Prdx4* for microglia, and *Prdx1* for oligodendrocyte were confirmed to be similar to the expression of the spatial patterns of each cell type, which were different on the comparison of wild type and 5xFAD mice. (WT: wild type; TG: 5xFAD mice; Prdx: Peroxiredoxin)

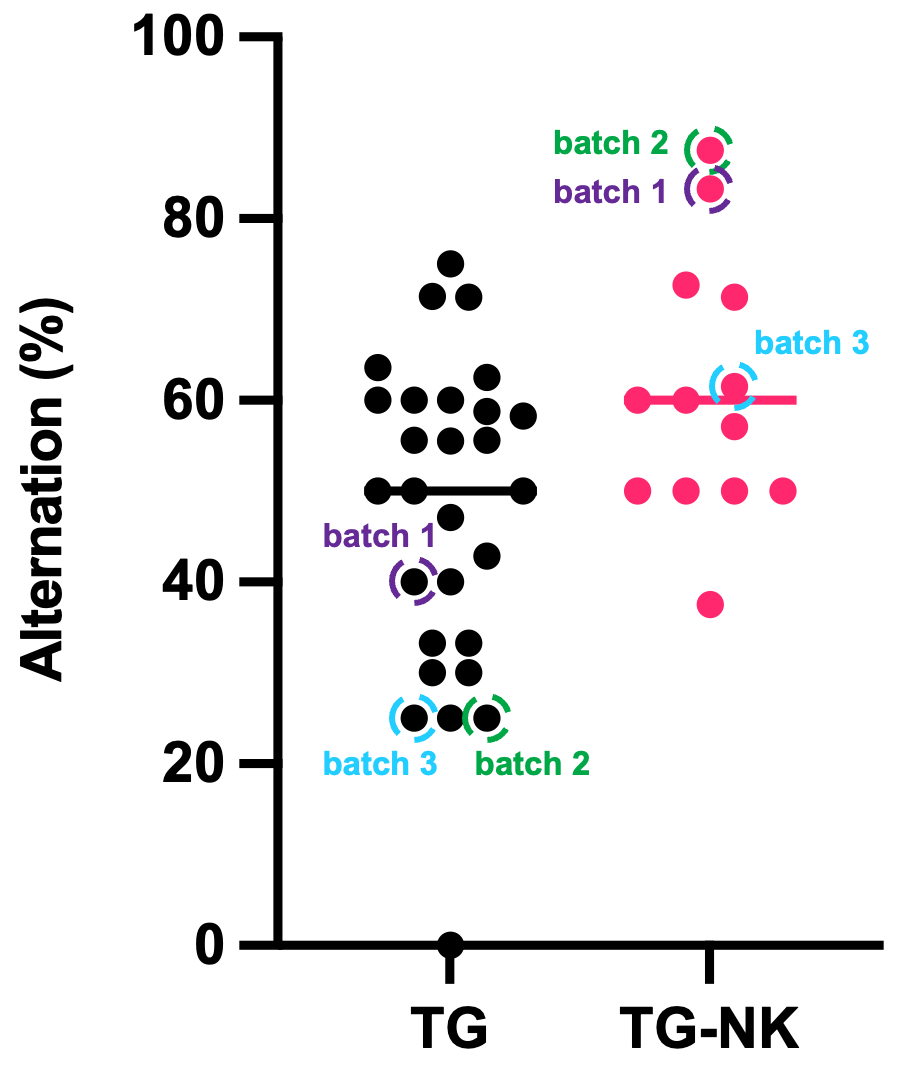

**Supplementary Figure 13. Display of batch-by-batch samples used for ST brain imaging analysis and their Y-maze results after intravenous administration of NK cell supplements in 5xFAD mice.**

Behavior of exploring new environment was examined using the Y-maze test and expressed as alternating percentages. Each dot represents a mouse in each group. Result of Y-maze behavior analysis of saline-treated and NK cell-treated 5xFAD mice. The three batches of mice used for ST brain imaging study were indicated. (NK: natural killer; TG: 5xFAD mice; TG_NK: NK cell-treated 5xFAD)

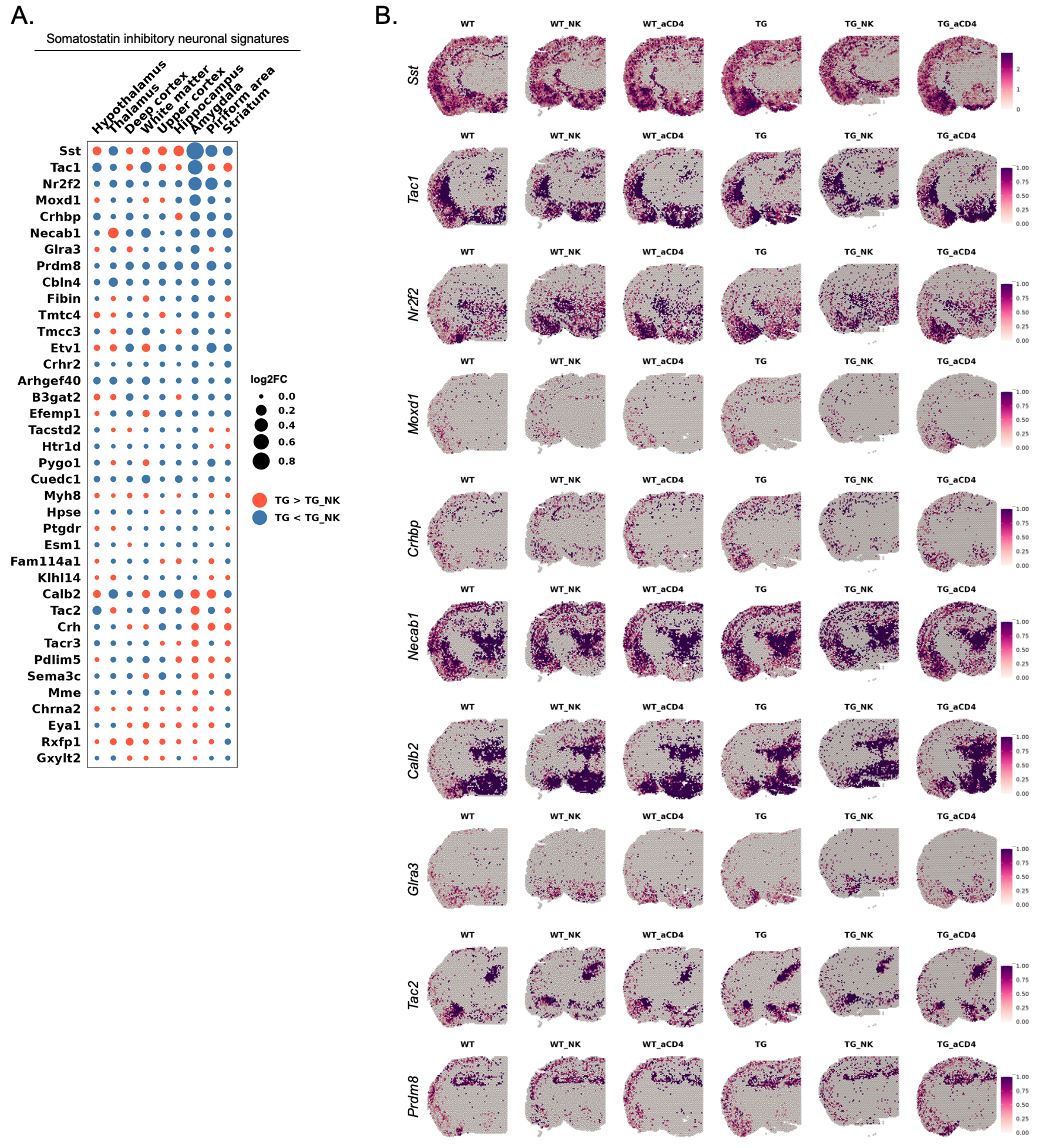

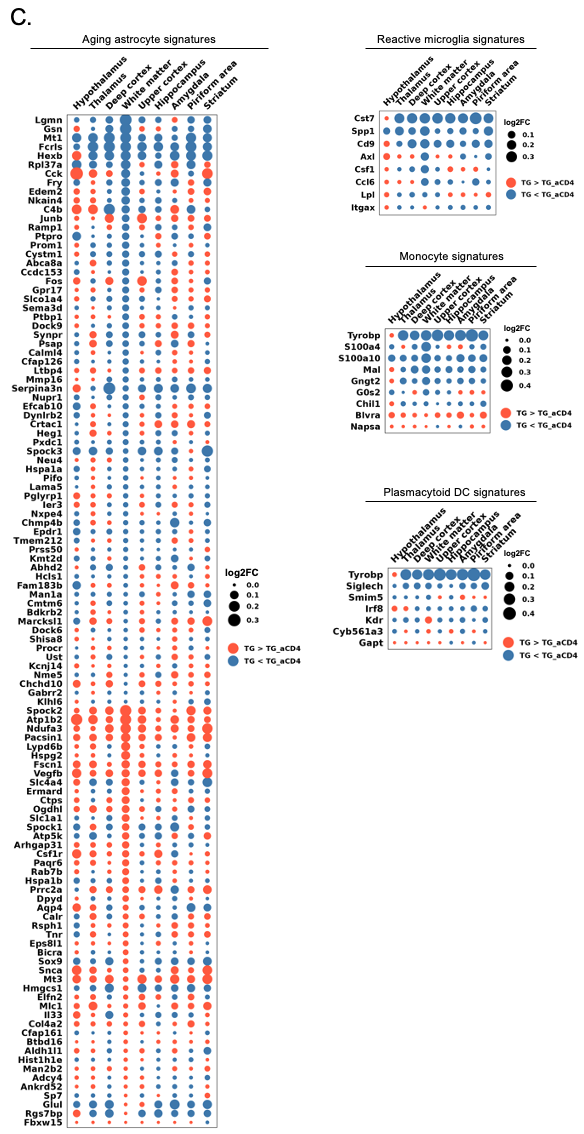

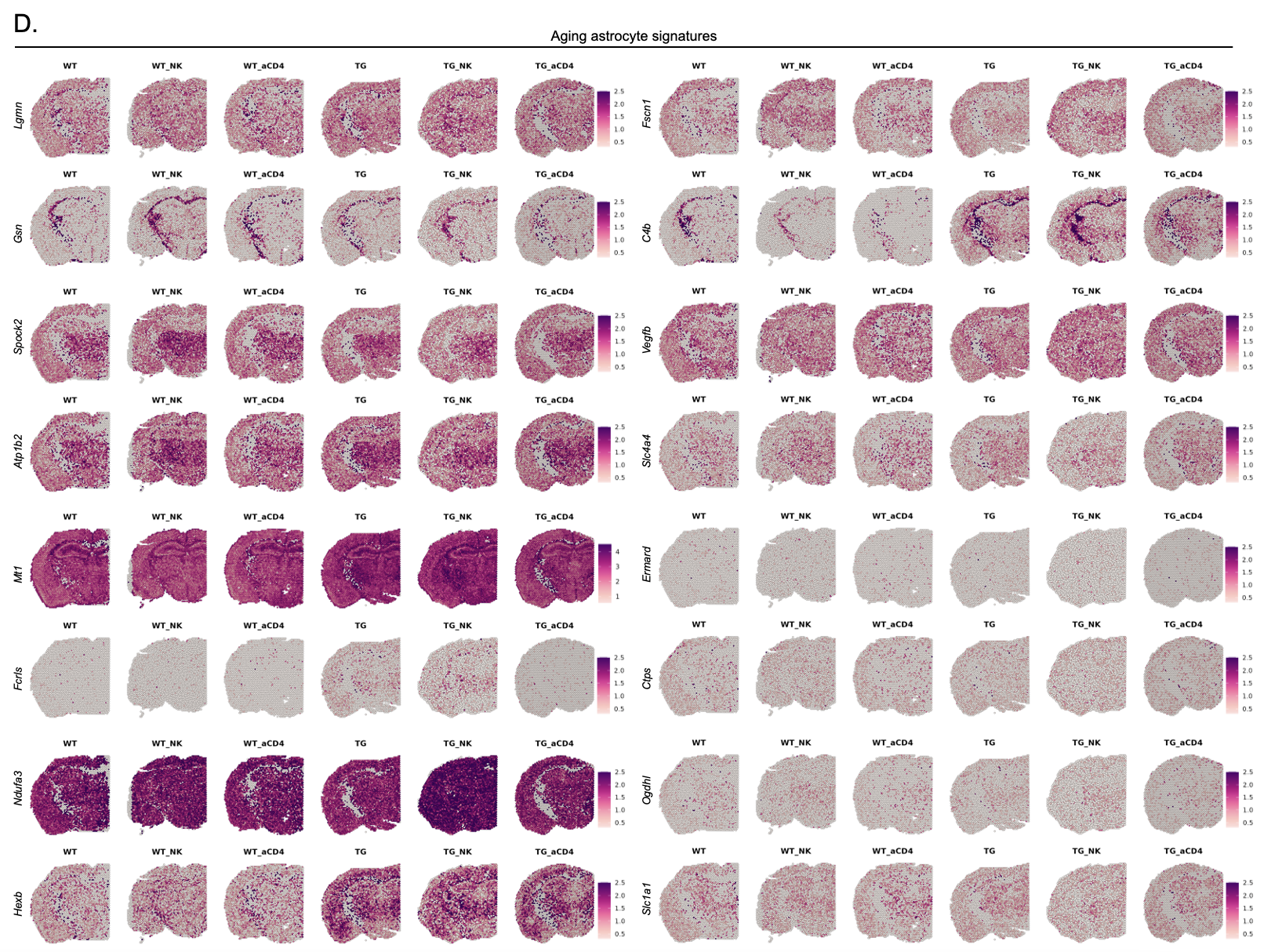

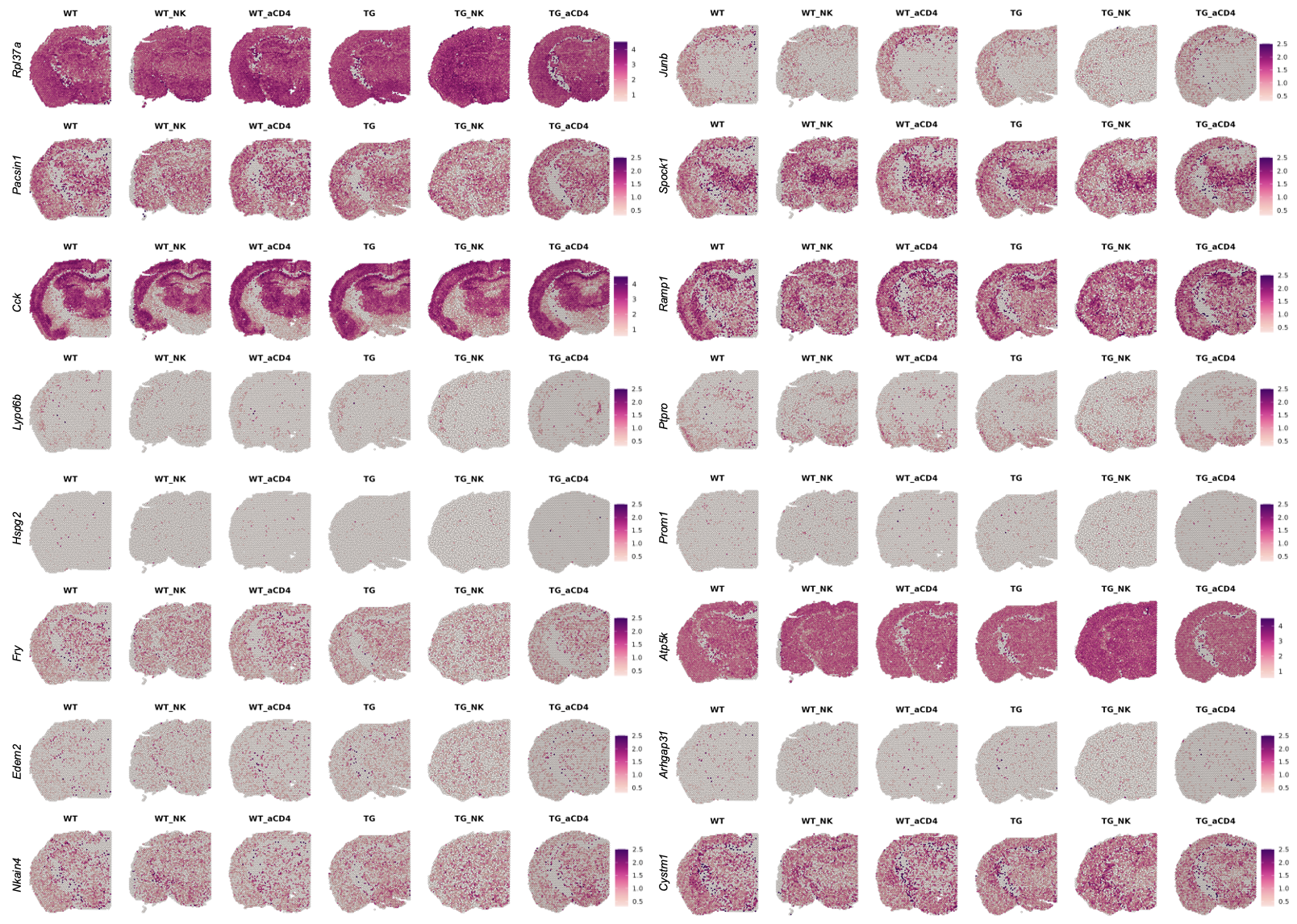

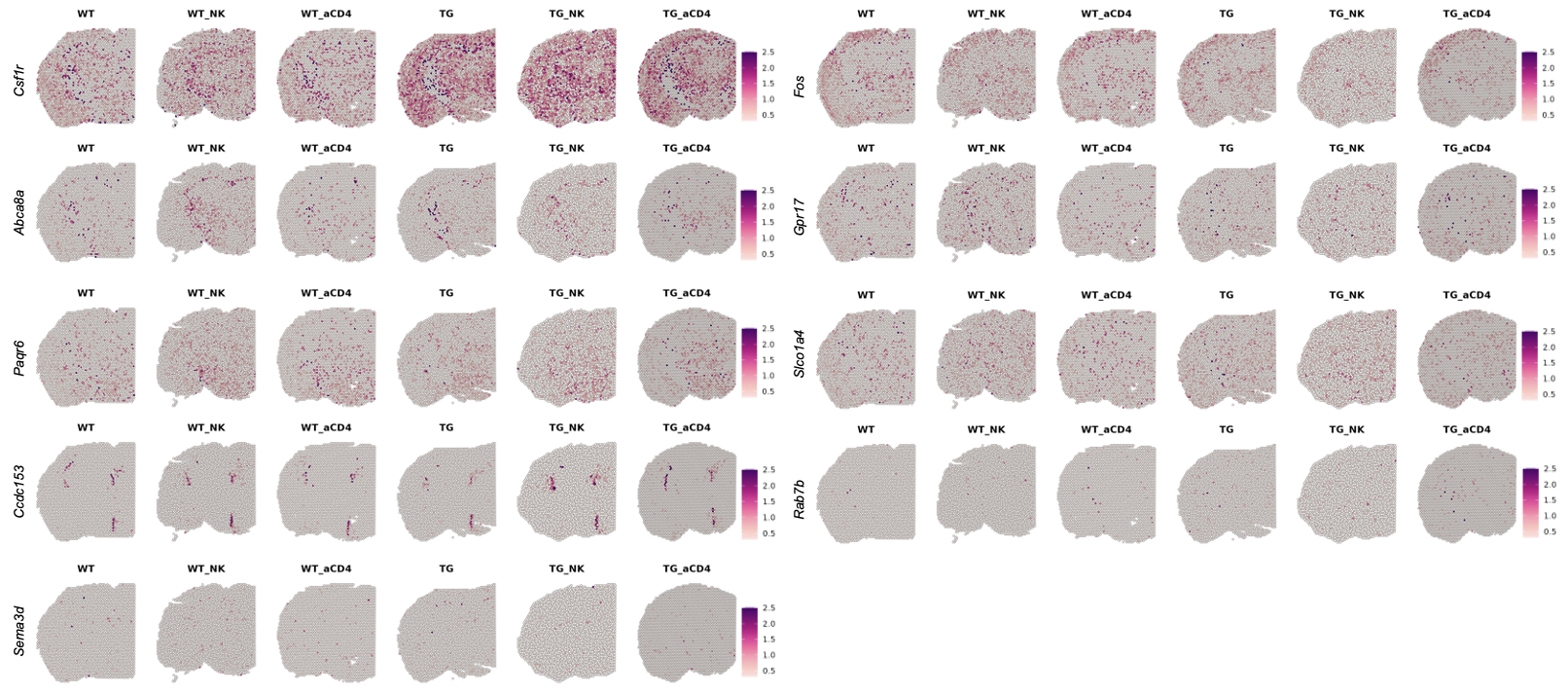

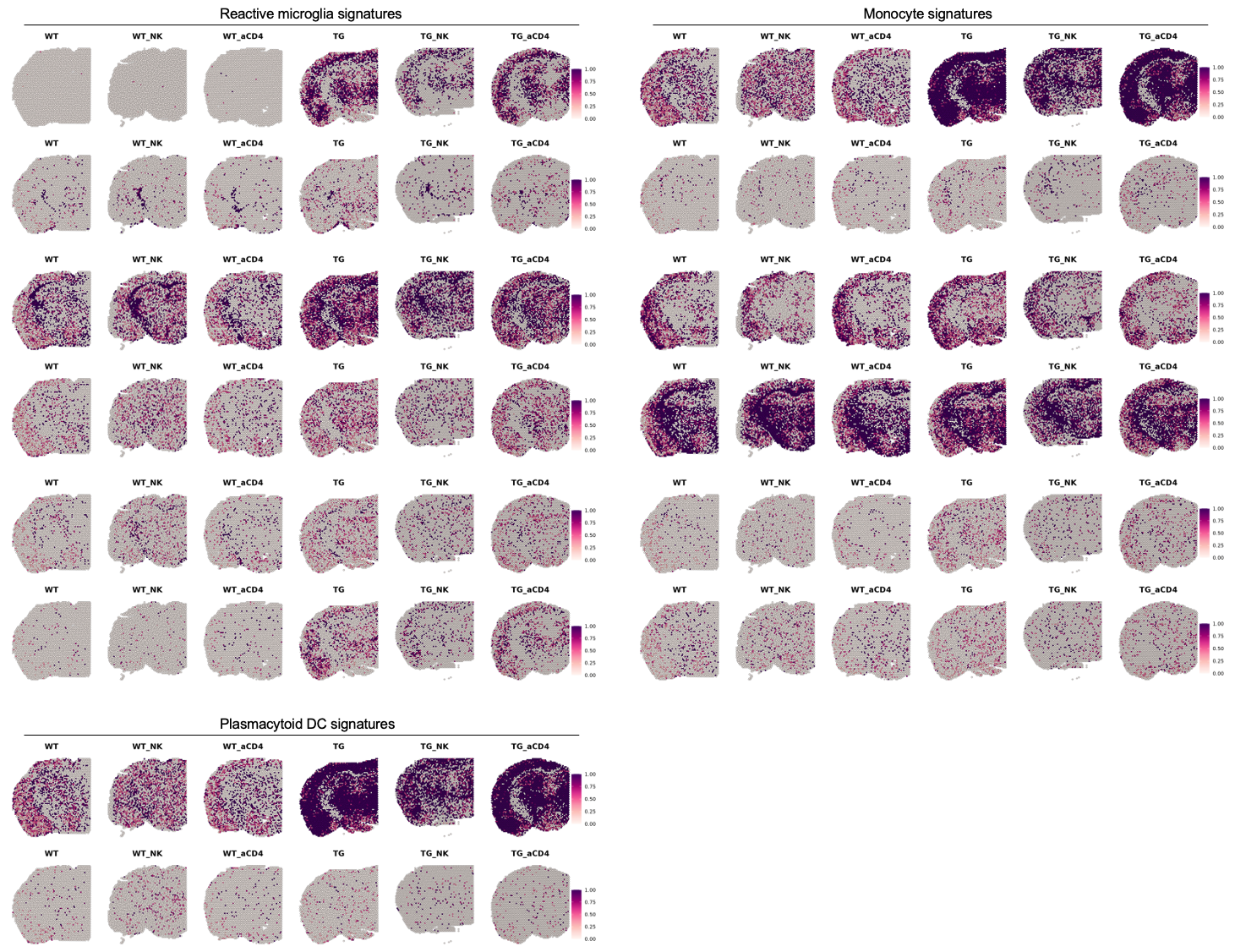

**Supplementary Figure 14. Spatial expression pattern of individual genes involved in changed brain cell signatures after NK cell supplement or anti-CD4 antibody treatment in 5xFAD mice.**

(A) Dot plot showing the average expression differences of individual genes of somatostatin inhibitory neuronal signatures in 9 different regions (hypothalamus, thalamus, deep cortex, white matter, hippocampus, upper cortex, amygdala, piriform area, and striatum). The average expression differences of each brain region between NK cell supplement- and saline-treated 5xFAD mice are indicated. Based on NK cell-treated 5xFAD mice, the blue dot indicates a decrease, and the orange dot indicates an increase. The size of the dots is proportional to the average expression difference value. (B) Spatial pattern of the selected individual genes showing at least 0.05 of the average expression difference values. The genes showed a tendency to decrease expression in the amygdala of 5xFAD mice after administration of NK cell supplement. In contrast, module score did not change by the anti-CD4 antibody treatment. (C) Dot plot showing the average expression differences of individual genes of immune cell signatures (aging astrocyte, reactive microglia, monocyte, and plasmacytoid DC). The average expression differences of each brain region between anti-CD4 antibody- and none-treated 5xFAD mice are indicated. (D) Spatial pattern of the selected individual genes showing at least 0.05 of the average expression difference values. The genes showed a tendency to decrease expression in the white matter of 5xFAD mice after administration of anti-CD4 antibody. Administration of NK cell supplement showed no appreciable changes in immune cell signatures. (aCD4: anti-CD4 antibody; WT: wild type; TG: 5xFAD mice; WT_NK: NK cell-treated wild type; WT_aCD4: anti-CD4 antibody-treated wild type; TG_NK: NK cell-treated 5xFAD; TG_aCD4: anti-CD4 antibody-treated 5xFAD; Sst: Somatostatin; DC: dendritic cells; NK: natural killer)

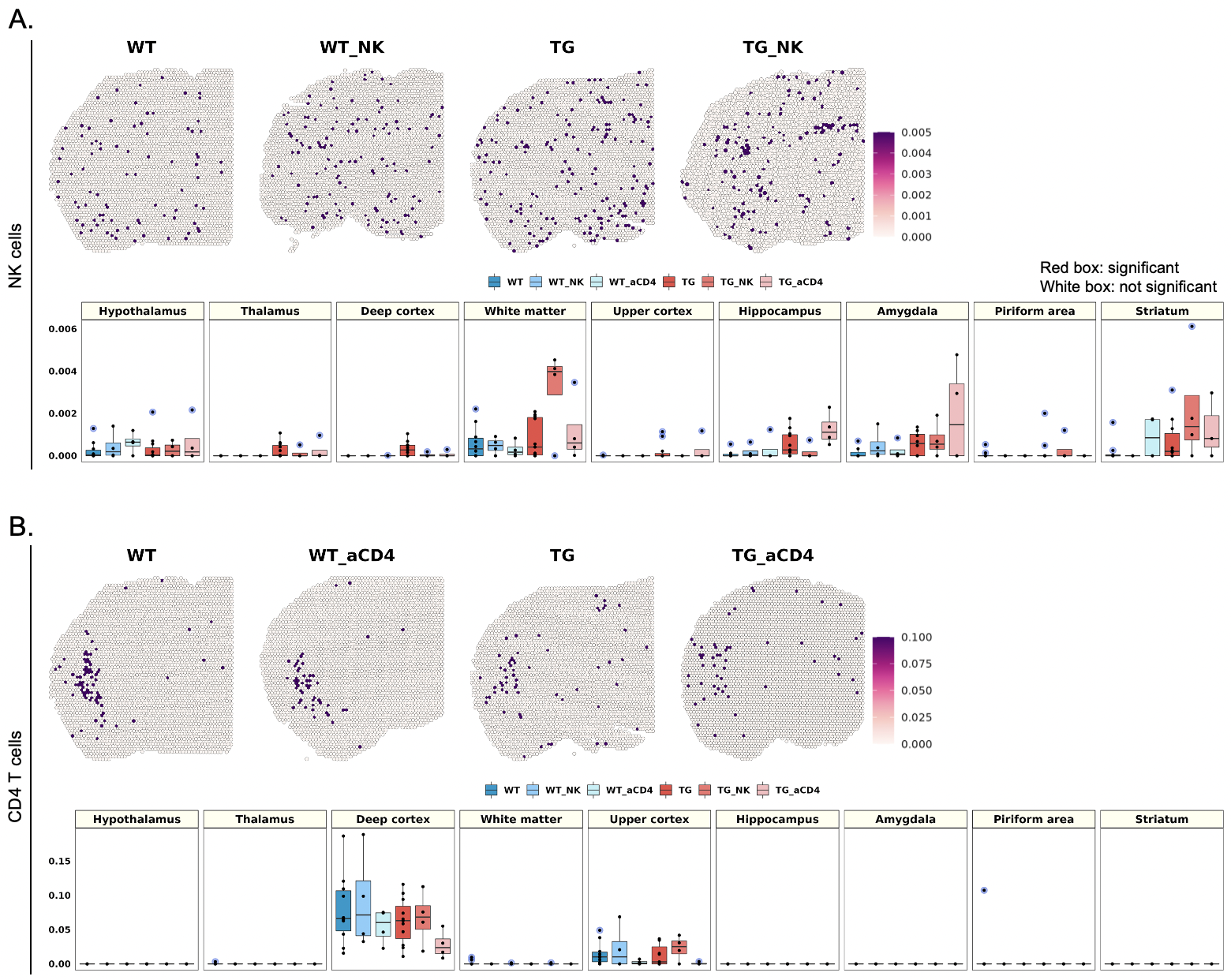

**Supplementary Figure 15. Spatial differences in the distribution of NK and CD4 T cell signatures with or without NK cell supplement or anti-CD4 antibody treatment in 5xFAD mice.**

(A) Spatial pattern of the NK cell signatures (upper row). Representative images of each group were selected among 10 spatial transcriptome data from wild type mice, 4 from NK cell-treated wild type mice, 4 from anti-CD4 antibody-treated wild type mice, 11 from 5xFAD mice, 4 from NK cell-treated 5xFAD mice, and 4 from anti-CD4 antibody-treated 5xFAD mice. Boxplot showing average module scores (lower row). Each dot represents a mouse in each group. Interestingly, the NK cell signature tended to increase after administration of NK cell supplements exclusively in the white matter of 5xFAD mice. In contrast, NK cell module score was not change by the anti-CD4 antibody treatment. (B) Spatial pattern of CD4 T cell signatures (upper row) and boxplot showing average module score levels (lower row). The expression level of the CD4 T cell signature tended to slightly decrease in the cortex after anti-CD4 antibody treatment. Administration of NK cell supplement showed no appreciable changes. (NK: natural killer; aCD4: anti-CD4 antibody; WT: wild type; TG: 5xFAD mice; WT_NK: NK cell-treated wild type; WT_aCD4: anti-CD4 antibody-treated wild type; TG_NK: NK cell-treated 5xFAD; TG_aCD4: anti-CD4 antibody-treated 5xFAD)

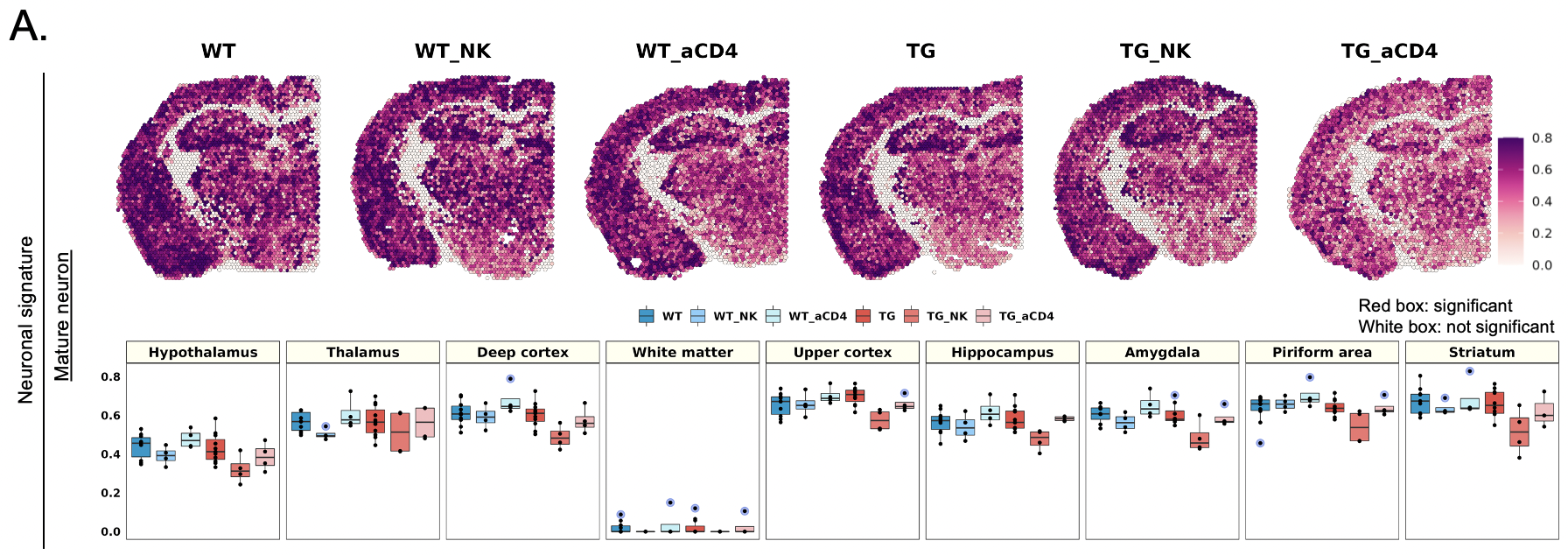

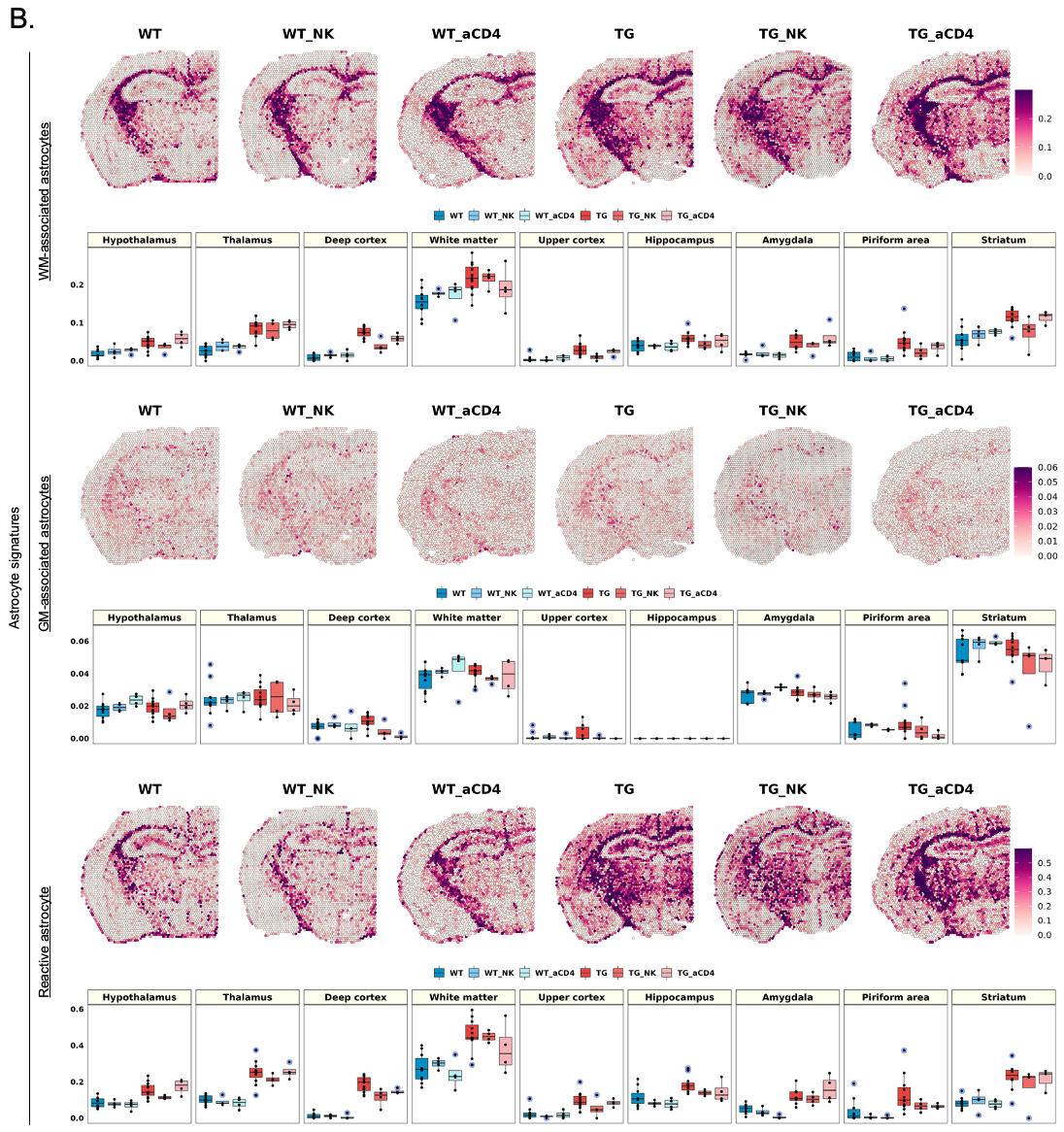

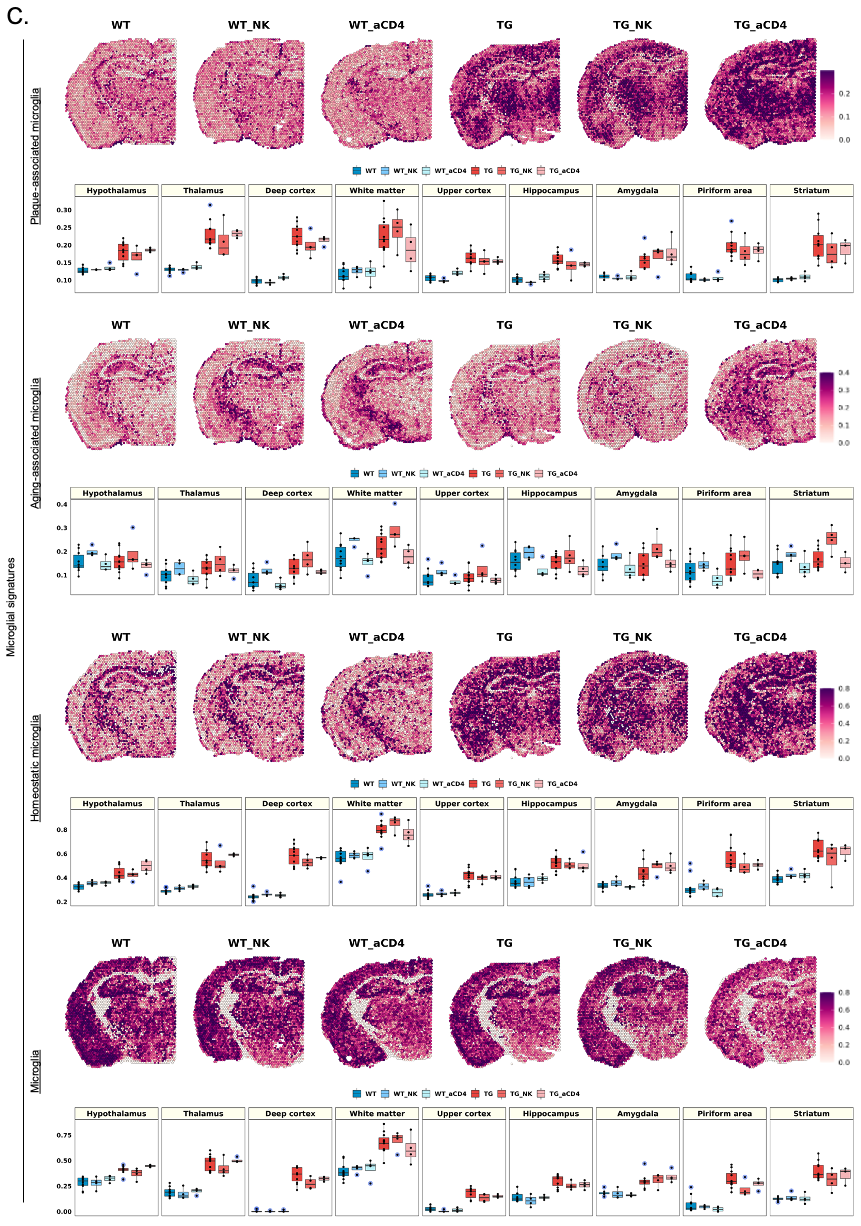

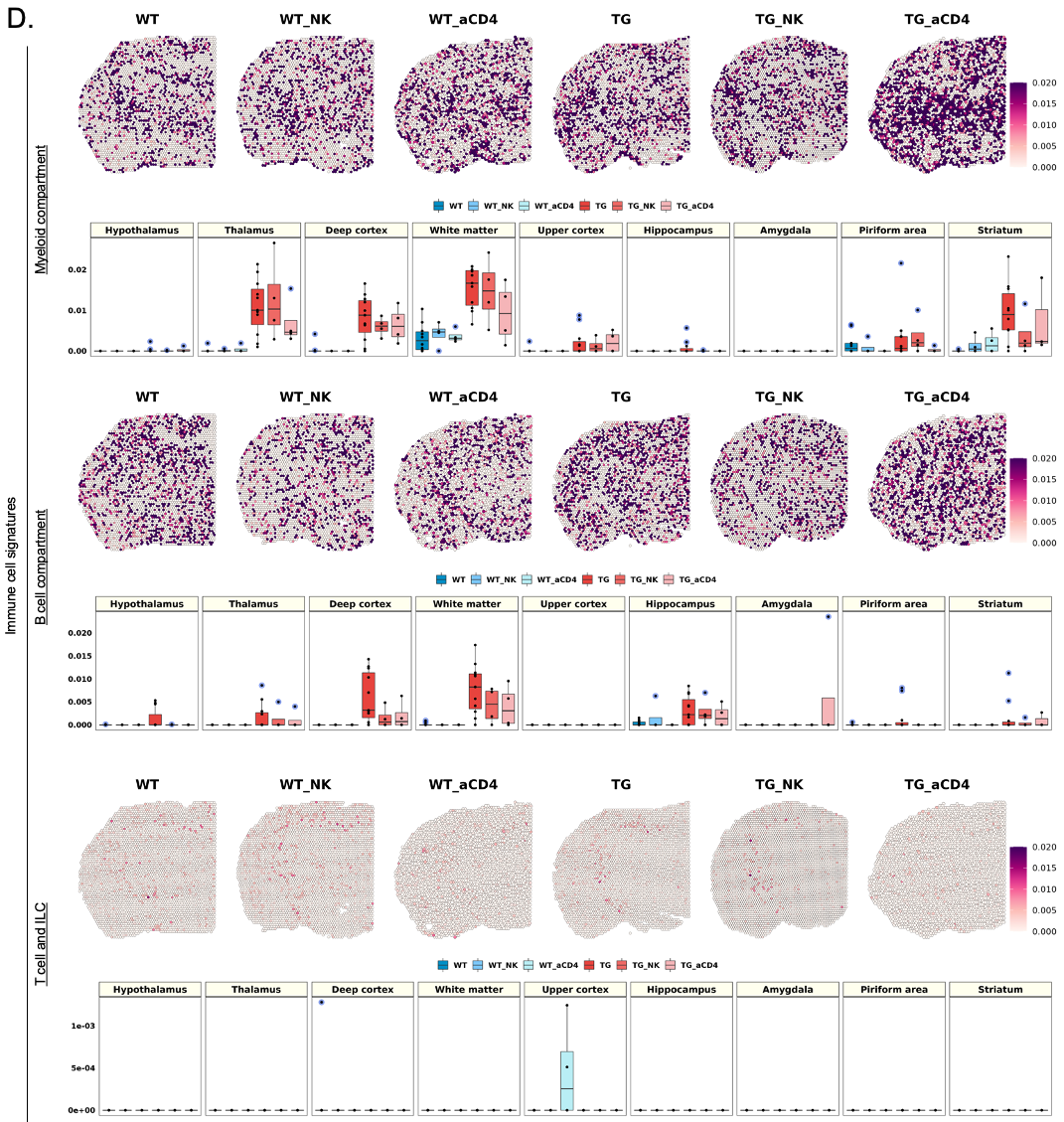

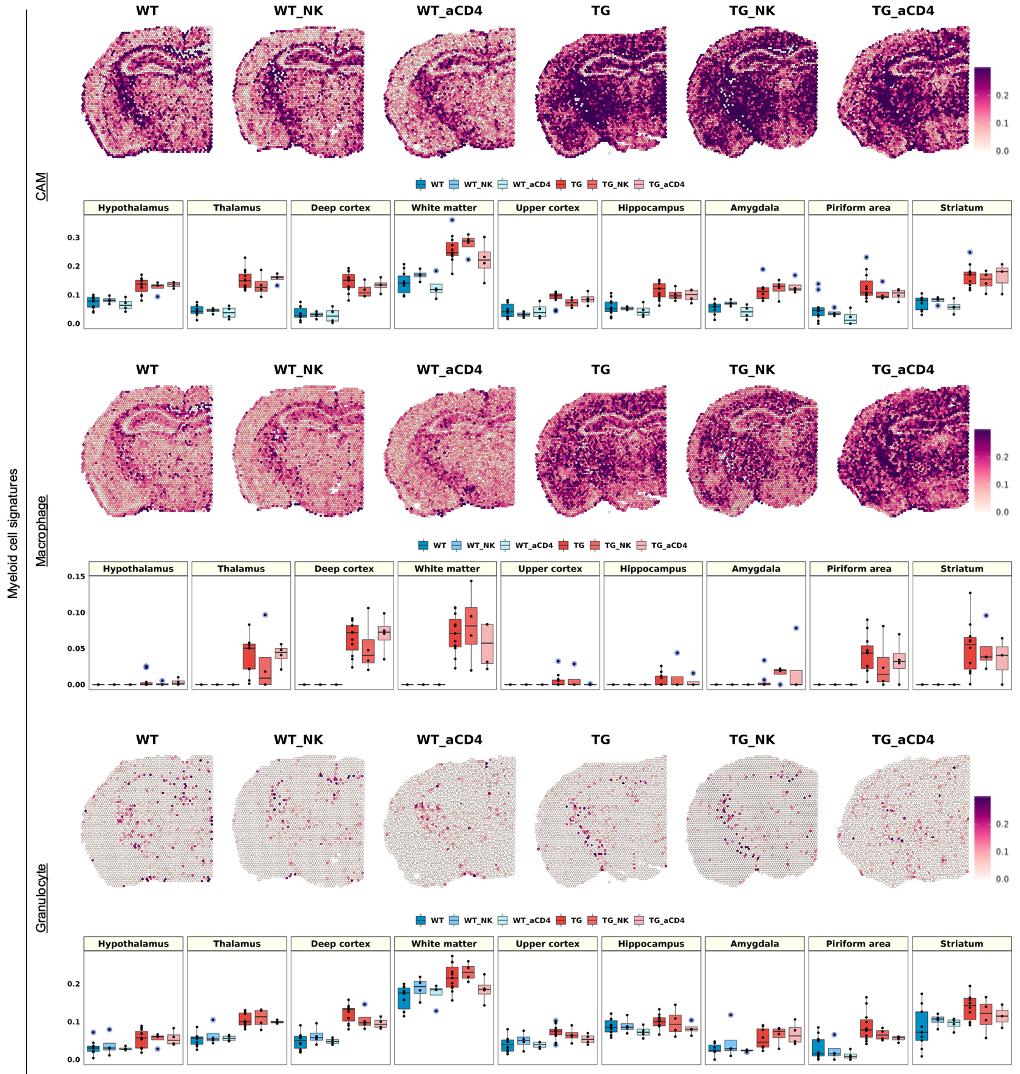

**Supplementary Figure 16. The distribution of brain cell signatures showing no changes in any region after NK cell supplement and anti-CD4 antibody treatment in 5xFAD mice.**

(A) Spatial pattern of the mature neuron signatures (upper row), (B) astrocyte signatures (WM-associated astrocytes, GM-associated astrocytes, and reactive astrocytes), (C) microglia signatures (plaque-associated microglia, aging-associated microglia, homeostatic, and pan microglia), and (D) immune cell signatures (myeloid compartment, B cell compartment, T cell and ILC, CAM, macrophage, and granulocyte) with and without NK cell supplements or anti-CD4 antibody treatment. Each boxplot shows the average module scores for each group (lower row). The expression of these brain cell signatures differed between wild type and 5xFAD mice, but were changed by neither NK cell supplements nor anti-CD4 antibody treatment. These negative findings are in contrast to the findings described in **Figure 5**, that NK cells supplement changed amygdala Sst neuron subtypes and anti-CD4 antibody treatment changed state-specific glial and immune cells (summarized in **Supplementary Table 6**). (NK: natural killer; aCD4: anti-CD4 antibody; WT: wild type; TG: 5xFAD mice; WT_NK: NK cell-treated wild type; WT_aCD4: anti-CD4 antibody-treated wild type; TG_NK: NK cell-treated 5xFAD; TG_aCD4: anti-CD4 antibody-treated 5xFAD; WM: white matter; GM: gray matter; ILC: innate lymphoid cells; CAM: CNS-associated macrophage)

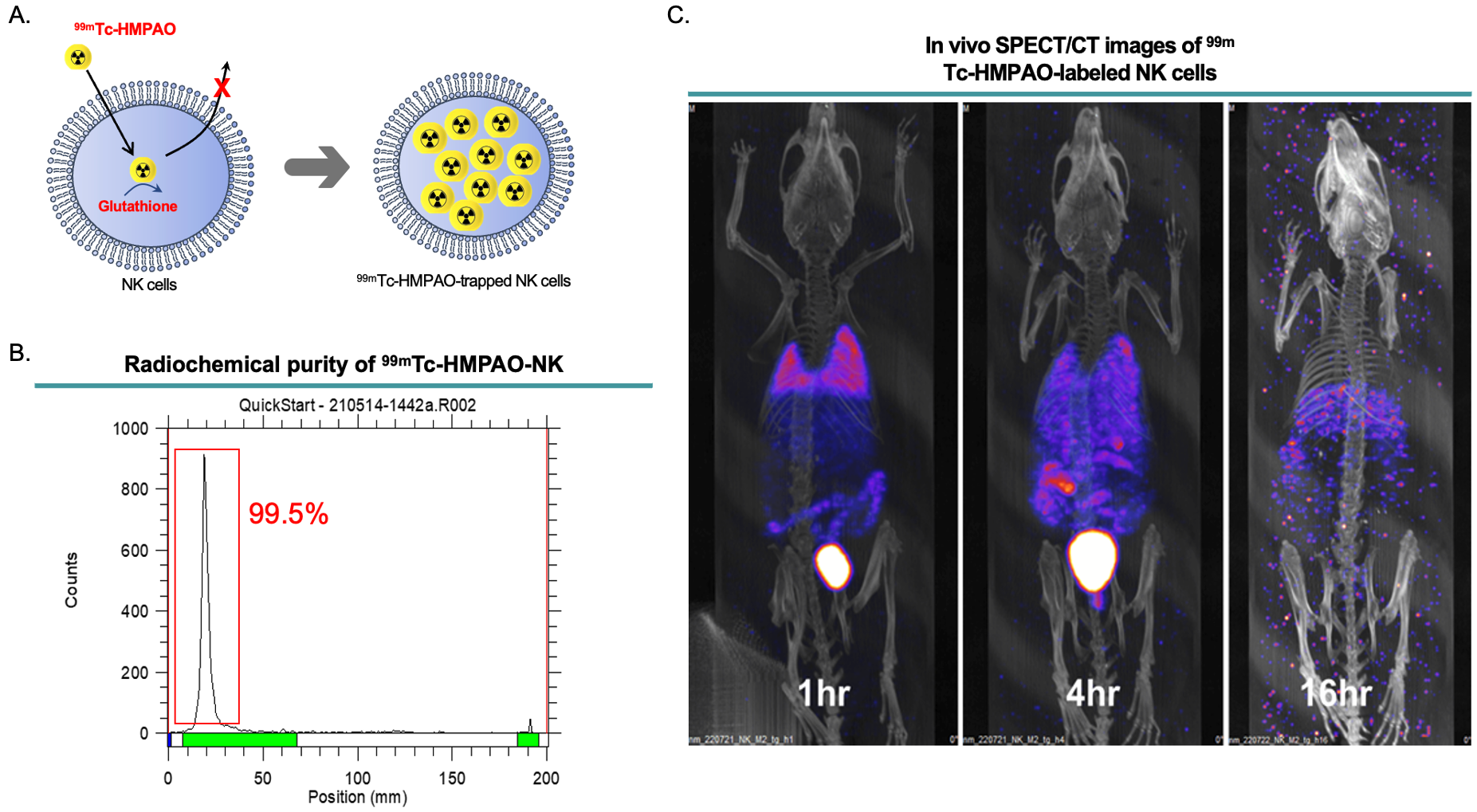

**Supplementary Figure 17. *In vivo* SPECT/CT images and biodistribution of ^99m^Tc-HMPAO-NK cells.**

(A) Schematic diagram for the radiolabeling method of ^99m^Tc-HMPAO-trapped NK cells. After highly lipophilic ^99m^Tc-HMPAO enters the cells, endogenous glutathione in cells begins to convert ^99m^Tc-HMPAO to a hydrophilic form, which is trapped inside the cells. (B) Radiochemical purification of the ^99m^Tc-HMPAO-NK cells. The labeled cells were analyzed by instant thin layered chromatography using Whatman No.1 paper. After radiochemical purification to discard remaining free ^99m^Tc-HMPAO, labeling efficiency was 99.5%. (C) *In vivo* SPECT/CT images of ^99m^Tc-HMPAO-NK cells in mice. After intravenous injection of ^99m^Tc-HMPAO-NK cells, SPECT/CT images were acquired at 1, 4, and 16 h in wild type C57BL/6 mice. Of note, no definite brain uptake of the labeled NK cells was observed. Injected NK cells may have caused, if any, the changes in brain cells at transcriptional levels by indirect pathways such as cytokine release or other secretory factors released by altered peripheral immune cells rather than direct effects. (NK: natural killer; ^99m^Tc-HMPAO: ^99m^technetium-hexamethylpropyleneamine oxime)
